# Fish diversity in a doubly landlocked country - a description of the fish fauna of Uzbekistan using DNA barcoding

**DOI:** 10.1101/2021.05.08.443274

**Authors:** Bakhtiyor Sheraliev, Zuogang Peng

## Abstract

Uzbekistan is one of two doubly landlocked countries in the world, where all rivers are endorheic basins. Although fish diversity is relatively poor in Uzbekistan compared to other regions, the fish fauna of the region has not yet been fully studied. The aim of this study was to establish a reliable barcoding reference database for fish in Uzbekistan. A total of 666 specimens, belonging to 59 species within 39 genera, 16 families, and 9 orders, were subjected to polymerase chain reaction amplification in the barcode region and sequenced. The length of the 666 barcodes was 682 bp. The average K2P distances within species, genera, and families were 0.22%, 6.33%, and 16.46%, respectively. The average interspecific distance was approximately 28.8 times higher than the mean intraspecific distance. The Barcode Index Number (BIN) discordance report showed that 666 specimens represented 55 BINs, of which five were singletons, 45 were taxonomically concordant, and five were taxonomically discordant. The barcode gap analysis demonstrated that 89.3% of the fish species examined could be discriminated by DNA barcoding. These results provide new insights into fish diversity in the inland waters of Uzbekistan and can provide a basis for the development of further studies on fish fauna.

## Introduction

Spanning more than 35,800 species [1], fish account for half of all extant vertebrate species and are well known for their uneven distribution of species diversity [2]. Consequently, fish constitute a significant component of biodiversity in the composition of animal taxa [3, 4]. Additionally, they have direct economic value and are important sources of animal protein for humans [5, 6]. However, every year the richness and abundance of fish biodiversity in aquatic ecosystems become more vulnerable, owing to human disturbances [7, 8]. Although approximately 400 new fish species have been described annually over the past 20 years [1], anthropogenic impacts, such as water pollution from plastic and other household waste, river dams, water withdrawal, overfishing, poaching, and habitat degradation have resulted in a catastrophic loss of fish diversity [9-11]. In-depth taxonomic studies of species are key to conserving biodiversity.

Generally, fish species identification and taxonomy rely on morphometric and meristic characteristics, such as body shape, the number of fin rays or lateral line scales, allometric features, and colour patterns. However, morphological characters are not always stable during various developmental stages and often cannot be assessed in incomplete samples or rare and cryptic species. Moreover, fish identification can be challenging, owing to the similar morphology of congeners during their early life histories as well as due to contradictions in the existing literature and taxonomic history; this is true even if experienced taxonomists work with whole intact adults. In addition, different taxonomists may have different identification abilities and skills, thus even the same specimen may be identified inconsistently, thereby resulting in confusion when summarising and comparing data [12-14]. However, environmental and conservation studies call for a high level of accuracy, requiring specimens to be identified entirely at the species level [15]. The inherent limitations of morphology-based taxonomy and the decreased number of taxonomists require molecular approaches for fish species identification [16].

Molecular identification, which identifies species using molecular markers, is widely used today. Among the various molecular approaches used for species molecular identification, DNA barcoding based on mitochondrial DNA (mtDNA) is one of the most suitable tools for species-level identification [17, 18]. In addition, mtDNA-based molecular identification has several advantages over morphological approaches. First, species identification does not require complete specimens; however, a tiny piece of tissue such as muscle, skin, fin, or teeth is acceptable for DNA extraction [18-20]. Second, DNA is more stable than morphological characters and is more resistant to degradation. For example, DNA can be extracted from water and soil previously occupied by an organism, or from samples that have been processed or digested [21-24]. Third, it is difficult to distinguish some species with similar morphological characteristics, such as cryptic or sibling species. Molecular identification can help accurately distinguish among such species [25, 26]. Fourth, DNA is invariable throughout the developmental stages of an organism. In contrast, morphological characters can change during a life cycle, thereby leading to species misidentification [12]. Therefore, molecular approaches can be applied in the identification of fish eggs, larvae, juveniles, and adults [13, 27]. Fifth, becoming a professional traditional taxonomist requires a lot of time, work, and resources [28, 29]. Advances in technology make it fairly easy to replicate and read DNA sequences, while bioinformatic software can automatically compare the resulting sequences; therefore, the training required to approach molecular identification is much less than that required for morphological identification. Molecular identification is widely used in a number of other fields besides species identification, including illegal species trade, food fraud, biological invasions, and biodiversity monitoring [30-33].

If mitochondrial DNA contains 37 genes, a number of mitochondrial genes, such as 12S ribosomal RNA (12S), 16S rRNA (16S), cytochrome b (CYTB), and control region (D-loop region), have been used as genetic markers for molecular identification [34-36]. Hebert et al. [17] pioneered the use of cytochrome c oxidase subunit I (COI) for molecular species identification, showing that this genetic marker can serve as a DNA barcode for biological identification in both invertebrates and vertebrates [18, 25, 37-39]. The Fish Barcode of Life Initiative (FISH-BOL) is an international research collaboration aimed at creating a standardised reference library of DNA barcodes for all fish species [40, 41]. The main goal of this project is to enable the identification of fish species by comparing the sequence of queries against the database of reference sequences in the Barcode of Life Data Systems (BOLD) [42]. To date, many studies have been carried out worldwide on fish DNA barcoding dedicated to FISH-BOL [3, 4, 18, 43, 44]. Compared to other regions of the world, studies devoted to fish barcoding are almost absent in Central Asia.

Uzbekistan is one of two doubly landlocked countries in the world, where all rivers are endorheic basins; therefore, fish biodiversity is poor. According to Mirabdullaev and Mullabaev [45], the total number of fish species in Uzbekistan exceeds 71, including 26 fish species introduced into the inland waters of the country. At the same time, the drying up of the Aral Sea, which is the largest water basin in the region, global climate change, population growth, river damming, water pollution, water withdrawals for agriculture, poaching, overfishing, and habitat destruction, all affect the fish species in the region [46, 47]. To date, studies on piscifauna have been based mainly on traditional morphological criteria and have not been comprehensively barcoded, except in our recent studies [48-50]. Recently, molecular identification has been applied to identify mainly nematodes among animal species [51].

Consequently, the main aim of the present study was to provide the first inventory of freshwater fish species in Uzbekistan based on DNA barcoding. This inventory could serve as a reference for screening DNA sequences in future studies. Additionally, we assessed the genetic diversity of freshwater fish species. The DNA barcode records generated in this study will be available to researchers for the monitoring and conservation of fish diversity in Uzbekistan.

## Results

### Morphology-based species identification

First, all collected specimens were identified using morphological approaches. Morphological identification classified all samples into 59 species belonging to 39 genera and 16 families that represented nine orders (Table 1). The identified specimens included 50 (84.75%) species identified to the species level and nine (15.25%) species that could not be identified to the species level (Tables 1 and S2). Approximately three-quarters of the species (44 species, 74.58%) belonged to the order Cypriniformes. The remaining eight orders included one or two species.

**Table 1.**
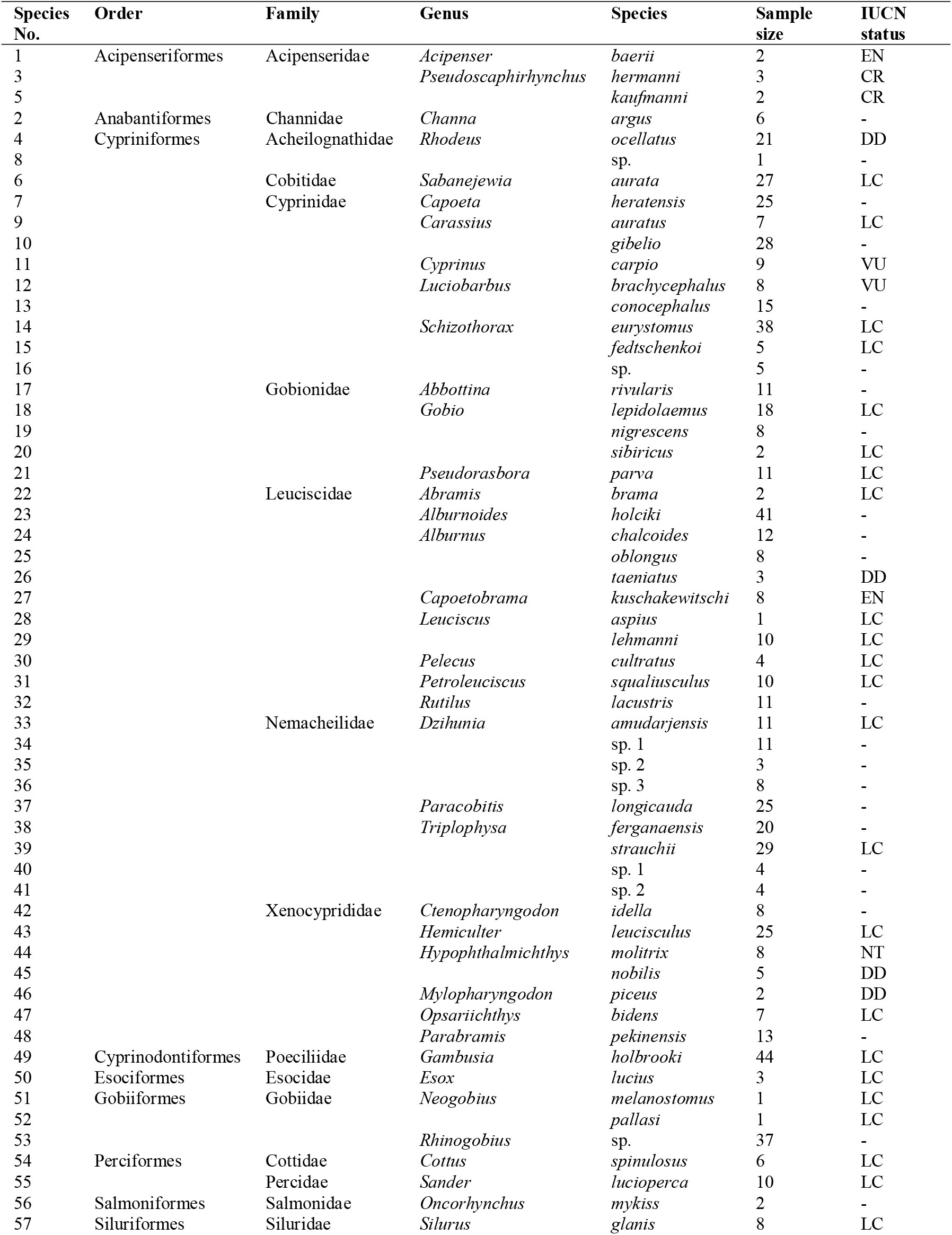
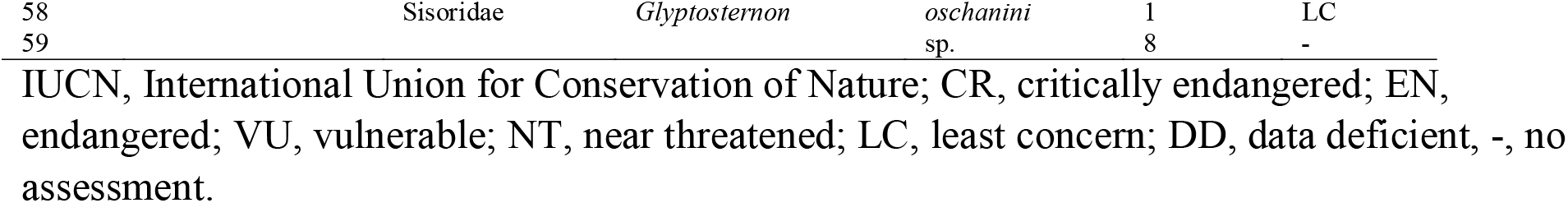
List of the fish species of Uzbekistan using in this study

Of the 59 fish species collected from the inland waters of Uzbekistan, *Pseudoscaphirhynchus hermanni* and *P. kaufmanni* were classified as critically endangered (CR), *Acipenser baerii* and *Capoetobrama kuschakewitschi* were classified as endangered (EN), and *Cyprinus carpio* and *Luciobarbus brachycephalus* were classified as vulnerable (VU) according to International Union for Conservation of Nature’s (IUCN) Red List of Threatened Species. The remaining species were grouped into the least concern (LC) and data deficient (DD) categories (Table 1).

### Identification of fish species using DNA barcodes

A total of 666 fish samples were successfully amplified using three primers and PCR. After editing, all COI barcode sequences were 682 for each sample and the mean nucleotide frequencies of the entire dataset were A (24.49%), T (29.01%), G (18.50%), and C (28.00%). The genetic distance within species ranged from 0.000 to 0.0149.

For species identification at the species level, a total of 666 COI barcode sequences representing 59 different species were employed (mean of 11.3 samples per species). The GenBank and BOLD databases were used for species identification (Table S2). The GenBank-based identification of all species ranged from 98.58% to 100.00%. The COI sequences of 22 fish species had not been reported in the GenBank database. Among them, *P. hermanni* was identified as *P. kaufmanni, Cottus spinulosus* as *C. ricei, L. conocephalus* as *L. capito, Alburnus oblongus* and *A. taeniatus* as *A. escherichii, Leuciscus lehmanni* and *Petroleuciscus squaliusculus* as *L. baicalensis*, and *Triplophysa* sp. 1 as *T. aliensis* with 99.71%, 98.47%, 98.83–100%, 98.39–98.82%, 99.71–99.85%, and 98.37% similarity, respectively.

The BOLD-based identification of 46 fish species ranged from 98.36% to 100%. No matches were found for 13 species. *Pseudoscaphirhynchus. hermanni* was identified as *P. kaufmanni, Cottus spinulosus* as *C. ricei, A. oblongus* and *A. taeniatus* as *A. escherichii, L. lehmanni* and *P. squaliusculus* as *L. baicalensis*, and *Triplophysa* sp. 1 as *T. aliensis* with 99.85– 100%, 98.48%, 98.62–98.92%, 99.8%–100%, and 98.36% similarity, respectively. Despite the GenBank databases, *L. conocephalus, Neogobius pallasi*, and *Rhinogobius* sp. were identified with high similarities (> 99.4%).

The Taxon ID tree shows that the specimens formed phylogenetic clusters that reflected previous taxonomic results based on morphology (Fig. S1). In turn, the barcode gap analysis revealed that five species lacked a barcode gap (intraspecific K2P distance ≥ interspecific one), and four species had a low K2P distance to another species (≤2%), which indicates that the majority of the investigated species could be identified by the DNA barcode approach (Table S3). Generally, the mean K2P distance of a species to its nearest neighbour (NN) was 8.04% (SD: 0.11%).

The mean K2P distances within species, within genera, and within families were 0.22%, 6.33%, and 16.46%, respectively (Table 2; Fig. 1). The largest intraspecific K2P distance was observed in *Opsariichthys bidens* (five specimens; Fig. 2; Table S3). The specimens obtained from several species, such as *Abramis brama* (two specimens), *Capoetobrama kuschakewitschi* (eight specimens), *Gobio nigrescens* (eight specimens), and *Rhinogobius* sp. (37 specimens), carried the same haplotype (Table S3). The average congeneric distance was approximately 28.8 times higher than the mean conspecific distance, but approximately 2.6 times less than the average genetic distance between families, thus the average genetic distance grew based on the taxonomic level.

**Table 2.** Summary of K2P genetic distances (%) calculated for different taxonomic levels

|  | N | Taxa | Comparisons | K2P genetic distance (%) |  |  |
| --- | --- | --- | --- | --- | --- | --- |
|  |  |  |  | Minimum | Maximum | Mean and SD |
| Within species | 661 | 54 | 6608 | 0.00 | 1.49 | 0.22 ± 0.00 |
| Within genus | 309 | 13 | 2534 | 0.00 | 11.78 | 6.33 ± 0.00 |
| Within family | 512 | 3 | 75196 | 0.00 | 22.19 | 16.46 ± 0.00 |

**Figure 1.**
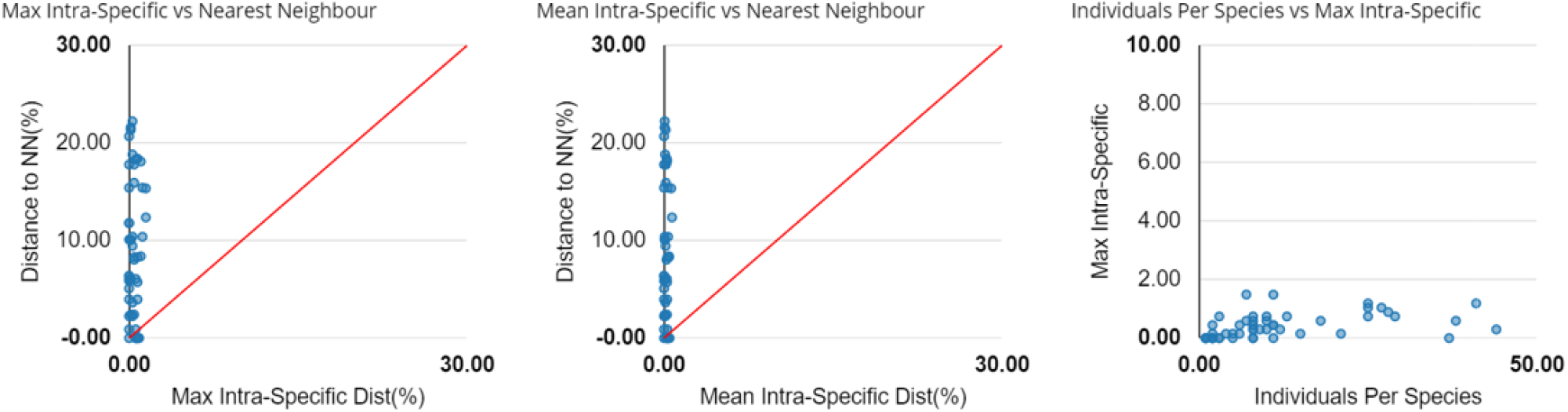
Barcoding gap: Maximum intraspecific Kimura 2-parameter (K2P) distances compared with the minimum interspecific K2P distances recorded in fish from Uzbekistan. The graphs show the overlap of the maximum and mean intra-specific distances with the inter-specific (NN = nearest neighbor) distances.

**Figure 2.**
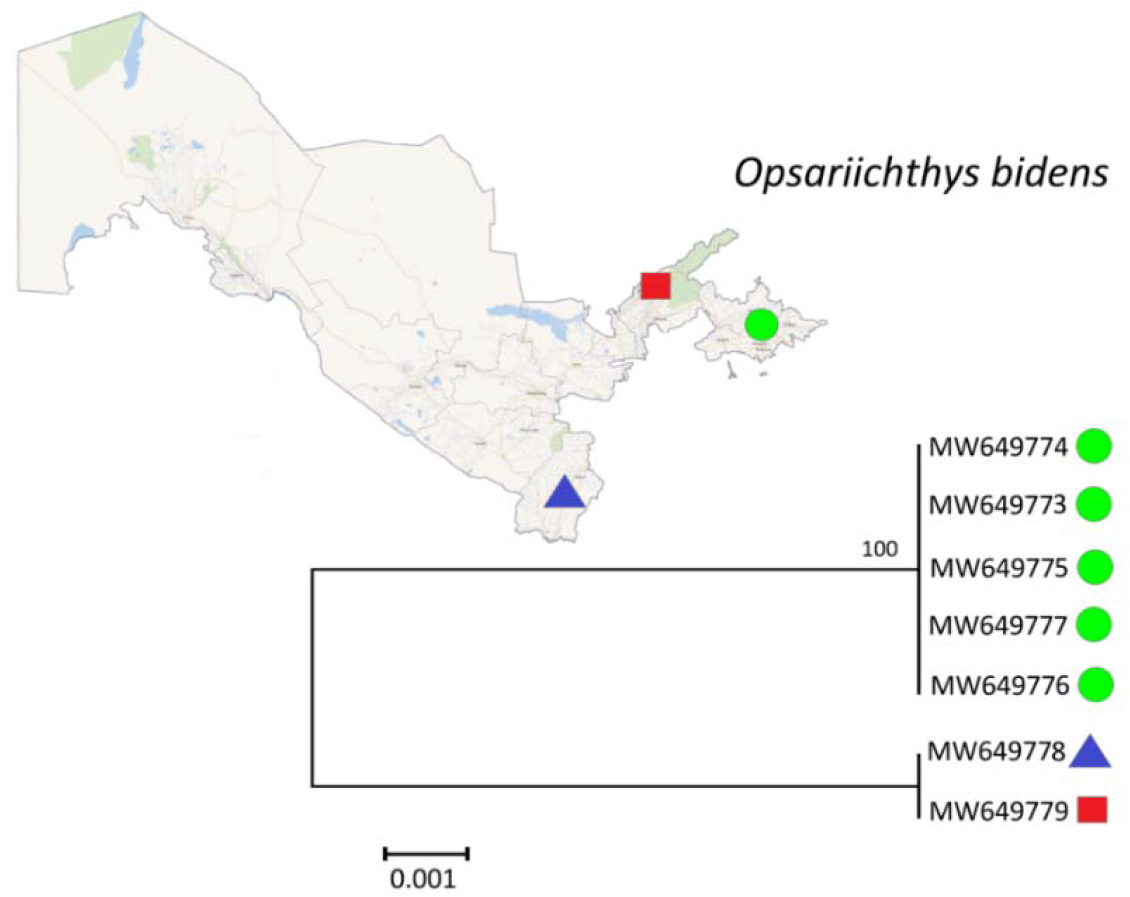
Neighbour-joining tree of *Opsariichthys bidens* from DNA barcode sequences with 100 000 bootstrapping replicates. Sampling localities: Syr Darya (green circle), Chirchik River (red square), and Surkhan Darya (blue triangle).

The Barcode Index Number (BIN) discordance report showed that 666 specimens represented 55 BINs; among them, 45 BINs were taxonomically concordant, five BINs were taxonomically discordant, and five BINs were singletons. For the best match (BM), best close match (BCM), and all species barcodes (ASB) analyses of the 666 sequence data set with singletons, the percentages of correct identification were 94.74%, 94.74%, and 89.03%, respectively; those of ambiguous identification were 4.05%, 4.05%, and 10.51%, respectively; those of incorrect identification were 1.2%, 1.2%, and 0.44%, respectively. Moreover, for the same three analyses of the dataset without singletons (661 sequences), the percentages of correct identification were 95.46%, 95.46%, and 89.71%, respectively; those of ambiguous identification were 3.93%, 3.93%, and 10.13%, respectively; those of incorrect identification were 0.6%, 0. %, and 0.15%, respectively (Table 3).

**Table 3.**
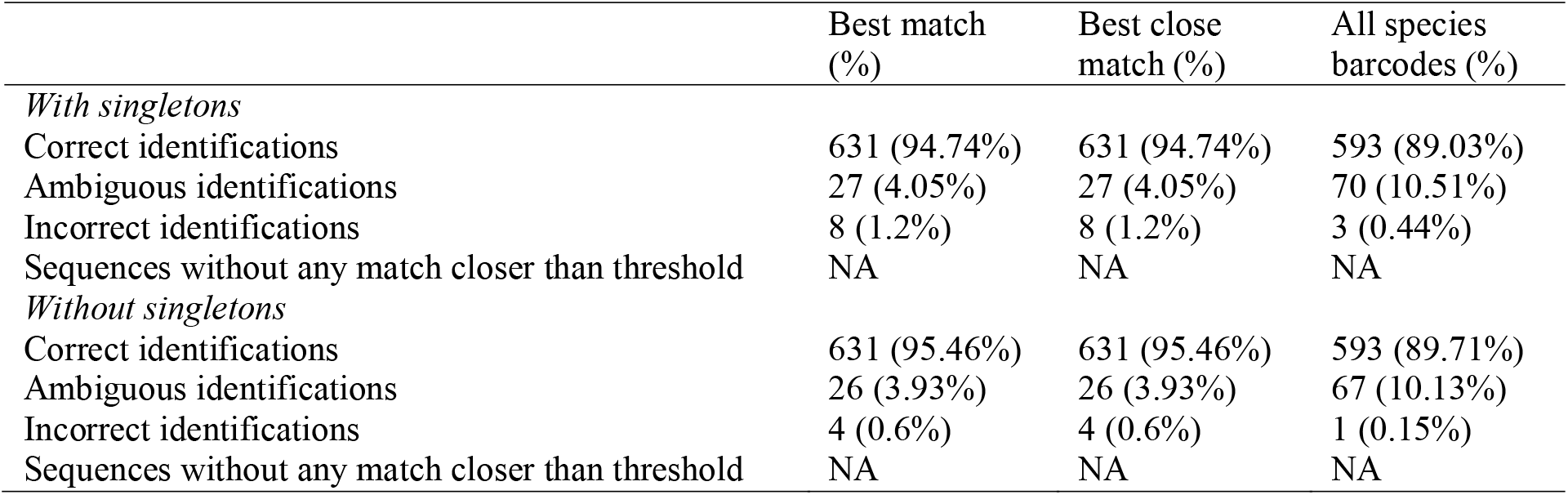
Results of identification success analysis for the criteria: best match, best close match and all species barcodes

|  | Best match (%) | Best close match (%) | All species barcodes (%) |
| --- | --- | --- | --- |
| *With singletons* |  |  |  |
| Correct identifications | 631 (94.74%) | 631 (94.74%) | 593 (89.03%) |
| Ambiguous identifications | 27 (4.05%) | 27 (4.05%) | 70 (10.51%) |
| Incorrect identifications | 8 (1.2%) | 8 (1.2%) | 3 (0.44%) |
| Sequences without any match closer than threshold | NA | NA | NA |
| *Without singletons* |  |  |  |
| Correct identifications | 631 (95.46%) | 631 (95.46%) | 593 (89.71%) |
| Ambiguous identifications | 26 (3.93%) | 26 (3.93%) | 67 (10.13%) |
| Incorrect identifications | 4 (0.6%) | 4 (0.6%) | 1 (0.15%) |
| Sequences without any match closer than threshold | NA | NA | NA |

### Automated barcode gap discovery (ABGD) analyses of species delimitation

The ABGD tool was used for species delimitation. A partition with prior maximal distance P = 0.0359 and 0.0046 delimited the entire dataset into 55 putative species (Fig. 3). Of the 59 morphological-based identified species, 55 (93.22%) were delimited clearly through the ABGD at a prior maximal distance of 0.0359, which was consistent with the observations of genetic distance and neighbour-joining (NJ) and Bayesian inference (BI) analyses (Figs. S1 and 4). Furthermore, at a prior maximal distance of 0.0359, few species, such as *Carassius auratus, C. gibelio, Gobio lepidolaemus, G. sibiricus, L. lehmanni, P. squaliusculus, P. hermanni*, and *P. kaufmanni* could not be delimited into different putative species. No clear divergence between these morphologically distinct species was observed in the NJ and BI analyses, with the exception of *Gobio* species.

**Figure 3.**
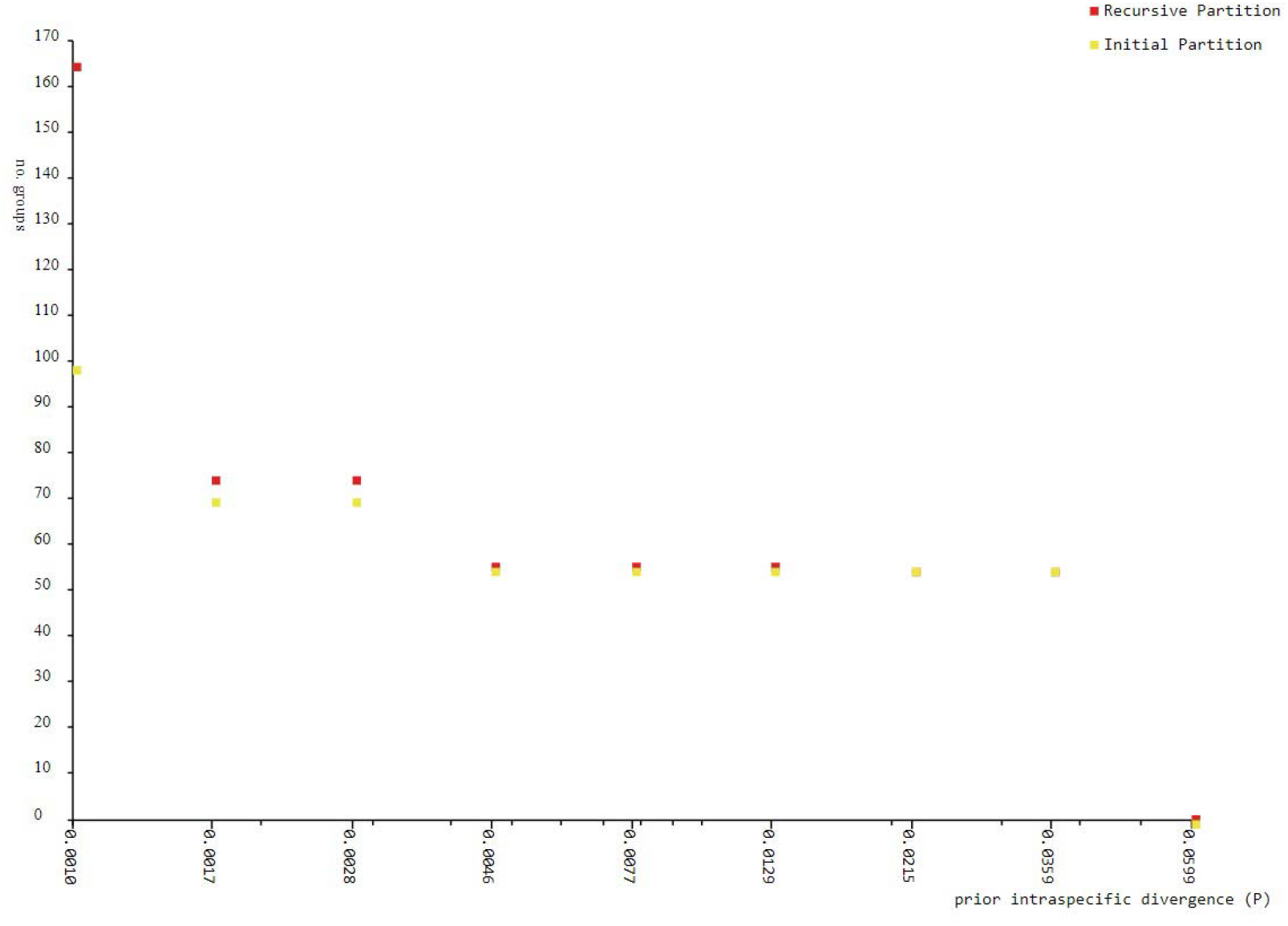
The number of groups inferred from ABGD analysis according to prior intraspecific divergence (*P*)

## Discussion

This study of the fish fauna of the inland waters of Uzbekistan is the first to compile the data in a sequence library, which contributes to the FISH-BOL in the BOLD system. This study included the molecular identification of 59 species. These 59 species included 83.1% of the reported fish fauna of the region [45]. Relationships among species are shown in the topology of the BI tree (Fig. 4).

**Figure 4.**
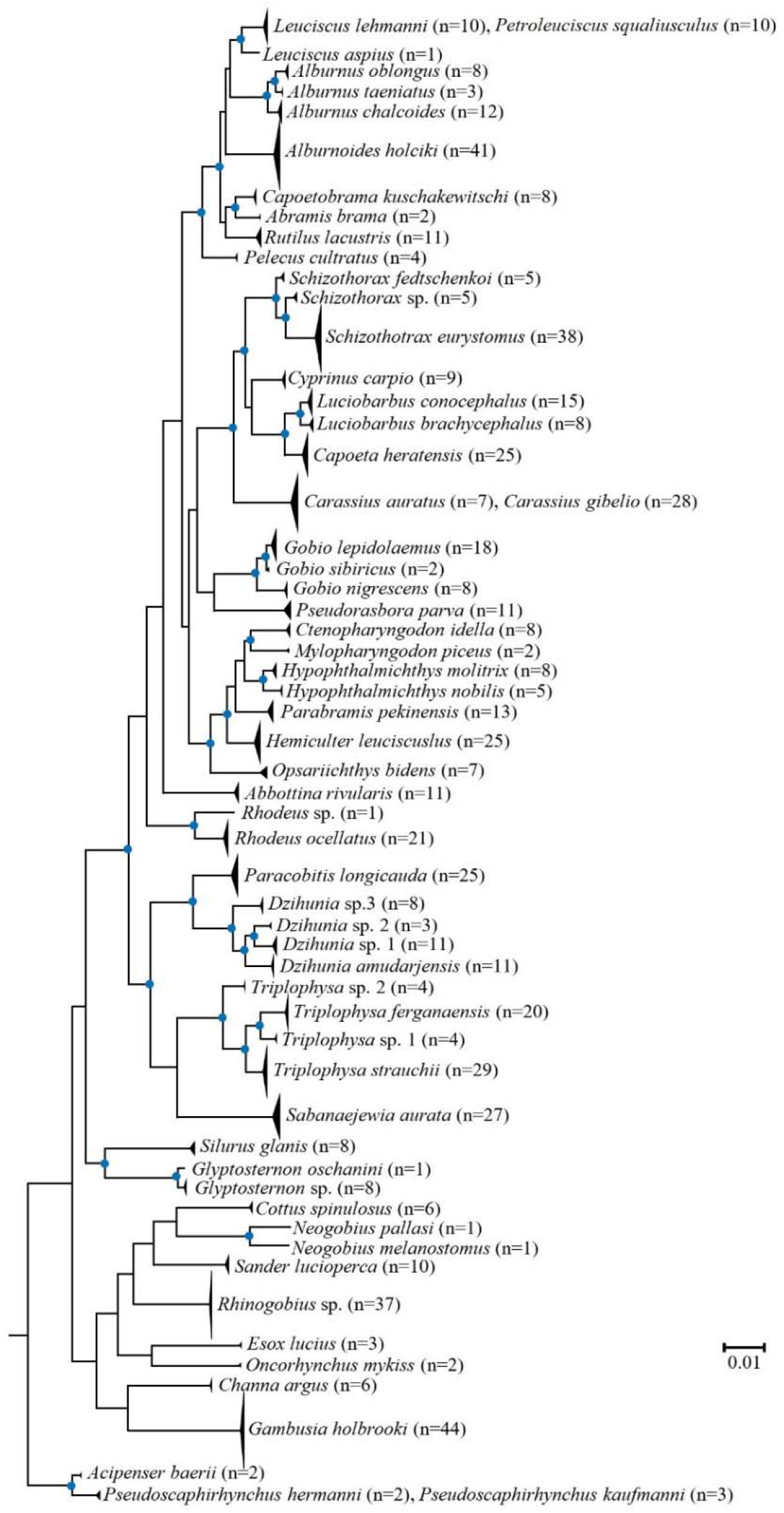
Bayesian inference (BI) consensus tree based on the COI partial gene sequences. The blue circle at nodes represents BI posterior probabilities values >50%. Posterior probability values for all species are >95%.

The gap between COI intraspecific and interspecific diversity is called the ‘barcode gap’, which is decisive for the discriminatory ability of DNA barcoding [52]. The barcode gap can be seen in our study (Table 2), as well as in many other previous studies [3, 44, 53], thereby further confirming that this approach is an effective way to distinguish between fish species.

This study clarified the taxonomic status of a number of taxa, such as *A. oblongus* and *A. taeniatus*, which belong to *Alburnus*, which is consistent with the results of Matveyev et al. [54] and Jouladeh-Roudbar et al. [55]; *Schizothorax fedtschenkoi* is a valid species; another *Schizothorax* sp. from the southern part of the country is an undescribed species; the *Alburnoides* population (previously considered as *A. eichwaldii*) from the inland waters of Uzbekistan, is de facto *A. holciki* [49]; three *Gobio* species occur in the inland waters of the country [50]; *Glyptosternon* and *Rhodeus* each consist of two species and not just one, as previously believed; thus, additional taxonomic research is required; two species of the genus *Neogobius* (*N. melanostomus* and *N. pallasi*) (previously believed to belong to *N. melanostomus* and *N. fluviatilis* [56]) occurred in the lower reaches of the Amu Darya; the population of *Opsariichthys* in Uzbekistan belongs to the same species, and *O. bidens* is not *O. unirostis* as previously believed [56]; the entire *Rhinogobius* population in Uzbekistan belongs to the same species (*Rhinogobius* sp.), which is neither *R. brunneus* nor *R. similis* as previously thought [56, 57]; thus, taxonomic clarification is required; moreover, there is only one species of *Gambusia* (*G. holbrooki*) occurring in the waters of Uzbekistan, while previously it was believed that both *G. affinis* and *G. holbrooki* were found in the waters of the country [56, 58] (Figs S1, 4; Table S2).

Only a single species of *Petroleuciscus* in Central Asia from the upper reaches of the Syr Darya, joined with *L. lehmanni* from the Zeravshan River in our phylogenetic analysis based on the *COI* barcode marker. However, our unpublished work (nuclear molecular and morphology) showed that they are two separate valid species, and *P. squaliusculus* belongs to *Leuciscus*.

Currently, three *Dzihunia* Prokofiev, 2001 species are found in the Amu Darya (*D. amudarjensis*), Zeravshan (*D. ilan*), and Talas (*D. turdakovi*, outside Uzbekistan) rivers [59, 60]. Apparently, the species diversity of *Dzihunia* seems to be much higher than previously thought (Fig. 4). In addition to *D. amudarjensis*, two more undescribed species were found in the upper reaches of Amu Darya. Another undescribed species was found in the Chirchik River; however, members of *Dzihunia* had not previously been found in this river (Fig. 5). On the other hand, *D. ilan* was not found in two of our expeditions to the Zeravshan River; moreover, it is believed that this species may have become extinct [59].

**Figure 5.**
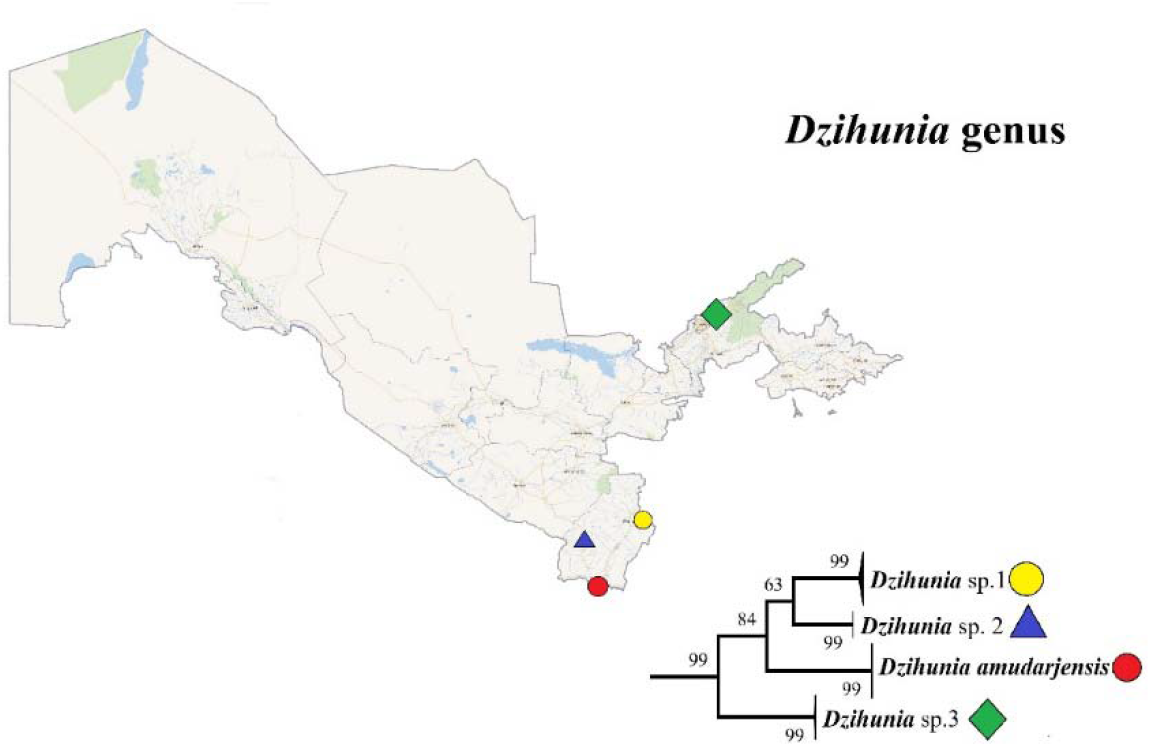
Neighbour-joining tree of *Dzihunia* spp. from DNA barcode sequences with 100 000 bootstrapping replicates. Sampling localities: lower Surkhan Darya (red circle) upper Surkhan Darya (yellow circle), Sherabad River (blue triangle) and Chirchik River (green square).

The inability of DNA barcodes to identify species may be due to incomplete sorting by lineage associated with recent speciation and haplotype sharing as a result of hybridisation. In our study, DNA barcodes of two *Leuciscus* and *Petroleuciscus* (*L. lehmanni* and *P. squaliusculus*), two *Carassius* (*C. auratus* and *C. gibelio*), and two *Pseudoscaphirhynchus* (*P. hermanni* and *P. kaufmanni*) species were sequenced, and the BIN discordance report illustrated that these six species could not be distinguished by the COI barcode gene (Figs. S1 and 4). In this case, a more rapidly evolving DNA fragment, such as the mitochondrial control region (mtCR) or the first internal transcribed ribosomal DNA spacer (ITS1), may be better for identification [3].

A similar situation occurred with *Carassius* species collected in the Mediterranean basin [61]. In addition, among the four *Leuciscus* (*L. baicalensis, L. bergi, L. dzungaricus*, and *L. lindbergi*) species from Central Asia, Russia, and Mongolia, no interspecific differences were found based on the COI gene (J. Freyhof, personal communication). However, in *Pseudoscaphirhynchus* species, no interspecies differences were found either when using other rapidly evolving mtDNA markers [62], the entire mtDNA genome [63], or nDNA markers (our unpublished data). In fact, these two sturgeon species are morphologically easy to distinguish from each other [64]. Thus, the complete genome sequencing of *Pseudoscaphirhynchus* may be important for their molecular authentication.

Unexpectedly, *Abbottina rivularis* from Gobionidae is nested with members of the genus *Rhodeus* from Acheilognathidae in our NJ phylogenetic tree (Fig. S1). Despite the sharp differences in morphology, the fact that these two genera are sister taxa has also been observed in previous studies [65, 66].

The global fish diversity is currently a serious threat. Along with natural limiting factors to native species, the negative impact of introduced species is also increasing [67-70]. At the same time, the negative impact of anthropogenic factors on the biodiversity of freshwater basins is also growing [71]. The number of biological species is declining annually; therefore, DNA barcoding is becoming a versatile approach that can be used to assess fish biodiversity, monitor fish conservation, and manage fishery resources [72-75]. While our DNA barcoding study is beneficial for the taxonomy and phylogenetics of fishes in the Amu Darya and Syr Darya basins, it is also important to clarify the taxonomy of misidentified invasive species acclimatised to Central Asian watersheds [58].

Unfortunately, fish diversity in Uzbekistan has decreased in recent years. A rare sturgeon fish, *Acipenser nudiventris*, is completely extinct in the Aral Sea basin [76]. Another sturgeon species endemic to the Syr Darya, *Pseudoscaphirhynchus fedtschenkoi*, has been possibly extinct since the 1990s [63]. The Syr Darya population of *Capoetobrama kuschakewitschi* has not been recorded in recent decades, and so far, this species has survived only in the lower reaches of the Amu Darya [77]. *Gymnocephalus cernuus* and *Perca fluviatilis* have not been recorded in water bodies in the country since the late 1990s [45]. Monitoring the existing populations of other rare native fish species and studying the negative impact of invasive species on them is advisable. The traditional monitoring of fish diversity is usually time-consuming, expensive, and labour intensive. However, with an ever-expanding barcode database and advances in biotechnology (such as environmental DNA analysis), the assessment of fish diversity is becoming more efficient [78-80]. As our molecular study of fishes develops in Uzbekistan, data on fish species in this region will become more readily available than ever.

## Methods

### Ethical Statement

Fish sampling for this research has complied with the Law of the Republic of Uzbekistan ‘On the protection and use of wildlife’ (No. 545-I 26.12.1997). No experimentation was conducted on live specimens in the laboratory, and the work performed in the laboratory followed the rules in the Guide for the Use and Care of Laboratory Animals of Southwest University.

### Sample collection and morphological identification

A total of 666 fish samples were collected from February 2016 to August 2020 using gill nets or cast nets from 53 distant locations in different rivers, tributaries, canals, springs, and lakes. Information about the sampling stations, along with geographical coordinates and sampling dates, is given in Table S1.

Initially, all specimens were identified to the species level based on morphological characteristics following the identification keys of Berg [64, 81] and Mirabdullaev et al. [82]. If identification was not correctly assigned to a specific species, the ‘sp.’ and ‘cf. abbreviations were applied [83]. Two pieces of right pectoral fin tissue and muscle tissue were dissected from each fish specimen and stored in 99% ethanol at -20 °C. Fin-clipped whole specimens and excess specimens for further morphological analyses were fixed in 10% formalin. After 5–7 days they were transferred to 70% ethanol for long-term storage and deposited in the Key Laboratory of Freshwater Fish Reproduction and Development at the Southwest University, School of Life Sciences (China), respectively, with the exception of sturgeon species, which were deposited in the Department of Biology at the Fergana State University, Faculty of Life Sciences (Uzbekistan).

### DNA extraction, COI amplification, and DNA sequencing

Genomic DNA was extracted from muscle or fin tissues by proteinase K digestion followed by a standard phenol-chloroform method. The DNA concentration was estimated using a nano-volume spectrophotometer (NanoDrop 2000; Thermo Fisher Scientific Inc., Waltham, MA, USA) and stored at -20 °C for further use. Approximately 680 bp were amplified from the 5’
s region of the COI gene using the fish-specific primers described by Ivanova et al. [84]: FishF2_t1 TGT AAA ACG ACG GCC AGT CGA CTA ATC ATA AAG ATA TCG GCA C and FishR2_t1 CAG GAA ACA GCT ATG ACA CTT CAG GGT GAC CGA AGA ATC AGA A, respectively. The following primers [18] were used for *Gambusia holbrooki*: FishF2-TCG ACT AAT CAT AAA GAT ATC GGC AC and FishR2-ACT TCA GGG TGA CCG AAG AAT CAG AA. The following primers [85] were used for sisorid catfishes: catF-TCT CAA CCA ACC ATA AAG ACA TTG G and catR-TAT ACT TCT GGG TGC CCA AAG AAT CA.

The PCR reactions were performed in a final volume of 25 µL, containing 10–100 ng template DNA, five µmol of each forward and reserve primer, while 12.5 µL of 2× *Taq* Master Mix (Novoprotein, Guangdong, China) and double-distilled water were also used. The thermal conditions consisted of an initial step of 3 min at 94 °C followed by 35 cycles of 0.5 min at 94 °C, 45 s at 54 °C, and 1 min 10 s at 72 °C, followed by a final extension of 7 min at 72 °C. The reactions were performed in an Applied Biosystems thermocycler (Veriti™ 96-Well Thermal Cycler, Singapore), and the PCR products were evaluated by electrophoresis using 1% agarose gel stained with BioRAD (Universal Hood II; Des Plaines, IL, USA). The PCR products were sent to TsingKe Biological Technology Co., Ltd. (Chongqing) for sequencing.

### Molecular data analysis

All sequences were manually edited using the SeqMan program (DNAStar software) combined with manual proofreading; all contig sequences started at the first codon position and ended at the third position; no stop codons were also detected. All obtained barcodes were uploaded to the BOLD and GenBank databases, and the details are given in Table S1.

The COI barcode sequence of each sample was identified by the scientific name or species using the BLAST and BOLD databases. Specimens were classified by family, genus, and species according to the fish taxonomic systems of Fricke et al. [60], and their status was checked in the IUCN Red List of Threatened Species v. 2020-3. The results of species identification based on the BLAST and BOLD databases are presented in Table S2.

We uploaded the entire data set to BOLD under project title ‘Freshwater fishes of Uzbekistan’. BOLD version 4 analytical tools were used for the following analyses. The distance summary with the parameter setting the Kalign alignment option [86] and pairwise deletion (ambiguous base/gap handling) was employed to estimate the Kimura 2-parameter (K2P) distances for taxonomic ranks at the species, genus, and family levels. Barcode gap analysis was carried out with the setting of the parameter ‘K2P; kalign alignment option; pairwise deletion (ambiguous base/gap handling)’ to construct the distribution of intraspecific and interspecific genetic distances [nearest neighbour (NN) analysis]. The BIN discordance report was employed to confirm the exactness of species identification, as well as to check for cases of low levels of genetic differentiation between different species. The Taxon ID tree was used to construct an NJ tree of the entire 666 sequences with the parameter-setting K2P distance model, the Kalign alignment algorithm [86], and pairwise deletion (ambiguous base/gap handling).

To verify intraspecific and interspecific genetic distances, we also used barcode gap analyses in ABGD. ABGD was used with K2P with the transition/transversion ratio (TS/TV) set to 2.0, 10 recursive steps, X (relative gap width) = 1.0; the remaining parameters were set to default values (Pmin = 0.001, Pmax = 0.1, Nb bins = 20).

We also used SPECIESIDENTIFIER v1.7.8 to verify species identification success by applying three criteria (BM, BCM, and ASB) to the entire barcode dataset, following Meier et al. [87]. Fish species that had only one sequence (singletons) were automatically assigned as ‘incorrectly identified’ under the BM and BCM criteria, as there were no conspecific barcoding sequences to match.

For phylogenetic reconstructions, the datasets were analysed based on the BI methodology using MrBayes 3.2. MrBayes was run with six substitution types (nst = 6), and we considered the gamma-distributed rate variation and the proportion of invariable positions (GTR+G+I) for the *COI* datasets. For BI, we ran four simultaneous Monte Carlo Markov chains for 25,000,000 generations, with sampling every 1,000 generations. The chain temperature was set at 0.2. Log-likelihood stability was determined after 10,000 generations, and we excluded the first 1,000 trees as burn-in. The remaining trees were used to compute a 50% majority-rule consensus tree. Moreover, to reveal the phylogenetic relationship of some fish species, the NJ tree of the K2P distance was constructed using MEGA7. Phylogenetic trees were visualised and edited using FigTree 1.4.2.

## Acknowledgements

We are thankful to Elbek Jalolov, Zuhriddin Obloqulov, Akbarjon Rozimov, Sirojiddin Allayarov, and Bahriddin Karimov for assistance in the field work. This work was funded by the grant from the National Natural Science Foundation of China (No. 31872204).

## Author contributions

Z.P. initiated the project, acquired funding, and managed project administration. B.S. collected specimens, preformed DNA extraction, analyzed the results and wrote the manuscript. Z.P. reviewed the manuscript. All authors read and approved the final version of the manuscript.

## Data availability

All sequences and associated voucher data are available from BOLD and GenBank. Voucher metadata are available in Supplementary Information.

## Additional information

Supplementary Information The online version contains supplementary material available at …

