## Supplementary material for "Fish diversity in a doubly landlocked country - a description of the fish fauna of Uzbekistan using DNA barcoding": Supplemetal Figure 1

| | |
|---|---|
| **Title** | **Fish diversity in a doubly landlocked country - a description of the fish fauna of Uzbekistan using DNA barcoding** |
| Authors | Bakhtiyor SHERALIEV[1,2], Zuogang PENG[1] |
| Affiliation | [1]Key Laboratory of Freshwater Fish Reproduction and Development (Ministry of Education), Southwest University, School of Life Sciences, Chongqing 400715, China |
| | [2]Faculty of Life Sciences, Fergana State University, Fergana, Uzbekistan |
| *Corresponding author | Professor Zuogang PENG, Ph.D. E-mail: |
| | |

**Figure S1.** Neighbor-joining tree based on the *COI* partial gene sequences

### BOLD TaxonID Tree

| | | |
|---|---|---|
| Title | : | Tree Result - FFU |
| Date | : | 26-Apr-2021 |
| Data Type | : | Nucleotide |
| Distance Model | : | Kimura 2 Parameter |
| Marker | : | COI-5P |
| Colourization | : | [blue]=Stop Codons [red]=Contamination or misidentification |
| Label | : | Process ID |
| Label | : | Taxon |
| Sequence Count | : | 666 |
| Species count | : | 59 |
| Genus count | : | 39 |
| Family count | : | 14 |
| Unidentified | : | 0 |
| BIN Count | : | 55 |

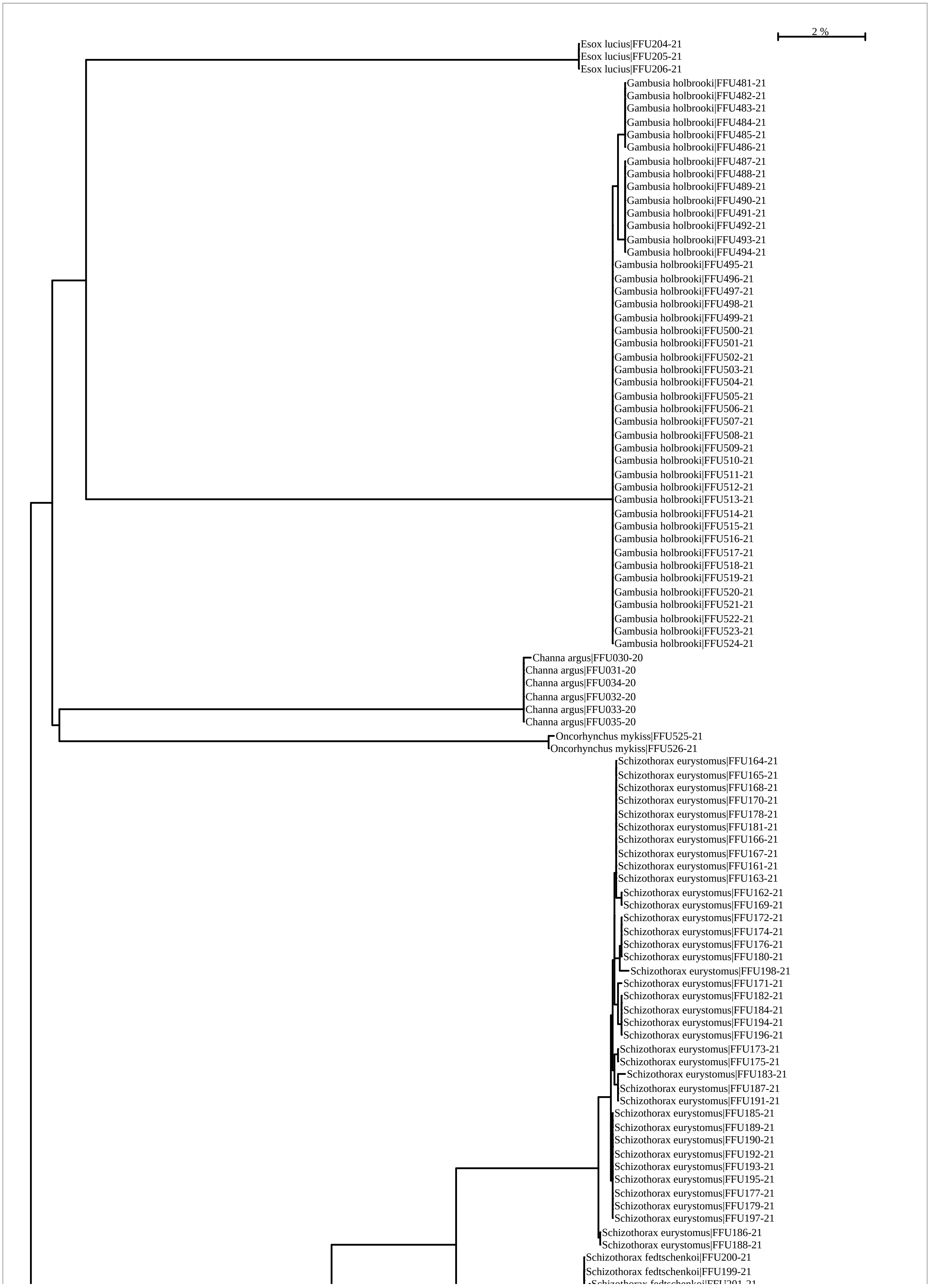
2 %
Esox lucius|FFU204-21
Esox lucius|FFU205-21
Esox lucius|FFU206-21
Gambusia holbrooki|FFU481-21
Gambusia holbrooki|FFU482-21
Gambusia holbrooki|FFU483-21
Gambusia holbrooki|FFU484-21
Gambusia holbrooki|FFU485-21
Gambusia holbrooki|FFU486-21
Gambusia holbrooki|FFU487-21
Gambusia holbrooki|FFU488-21
Gambusia holbrooki|FFU489-21
Gambusia holbrooki|FFU490-21
Gambusia holbrooki|FFU491-21
Gambusia holbrooki|FFU492-21
Gambusia holbrooki|FFU493-21
Gambusia holbrooki|FFU494-21
Gambusia holbrooki|FFU495-21
Gambusia holbrooki|FFU496-21
Gambusia holbrooki|FFU497-21
Gambusia holbrooki|FFU498-21
Gambusia holbrooki|FFU499-21
Gambusia holbrooki|FFU500-21
Gambusia holbrooki|FFU501-21
Gambusia holbrooki|FFU502-21
Gambusia holbrooki|FFU503-21
Gambusia holbrooki|FFU504-21
Gambusia holbrooki|FFU505-21
Gambusia holbrooki|FFU506-21
Gambusia holbrooki|FFU507-21
Gambusia holbrooki|FFU508-21
Gambusia holbrooki|FFU509-21
Gambusia holbrooki|FFU510-21
Gambusia holbrooki|FFU511-21
Gambusia holbrooki|FFU512-21
Gambusia holbrooki|FFU513-21
Gambusia holbrooki|FFU514-21
Gambusia holbrooki|FFU515-21
Gambusia holbrooki|FFU516-21
Gambusia holbrooki|FFU517-21
Gambusia holbrooki|FFU518-21
Gambusia holbrooki|FFU519-21
Gambusia holbrooki|FFU520-21
Gambusia holbrooki|FFU521-21
Gambusia holbrooki|FFU522-21
Gambusia holbrooki|FFU523-21
Gambusia holbrooki|FFU524-21
Channa argus|FFU030-20
Channa argus|FFU031-20
Channa argus|FFU034-20
Channa argus|FFU032-20
Channa argus|FFU033-20
Channa argus|FFU035-20
Oncorhynchus mykiss|FFU525-21
Oncorhynchus mykiss|FFU526-21
Schizothorax eurystomus|FFU164-21
Schizothorax eurystomus|FFU165-21
Schizothorax eurystomus|FFU168-21
Schizothorax eurystomus|FFU170-21
Schizothorax eurystomus|FFU178-21
Schizothorax eurystomus|FFU181-21
Schizothorax eurystomus|FFU166-21
Schizothorax eurystomus|FFU167-21
Schizothorax eurystomus|FFU161-21
Schizothorax eurystomus|FFU163-21
Schizothorax eurystomus|FFU162-21
Schizothorax eurystomus|FFU169-21
Schizothorax eurystomus|FFU172-21
Schizothorax eurystomus|FFU174-21
Schizothorax eurystomus|FFU176-21
Schizothorax eurystomus|FFU180-21
Schizothorax eurystomus|FFU198-21
Schizothorax eurystomus|FFU171-21
Schizothorax eurystomus|FFU182-21
Schizothorax eurystomus|FFU184-21
Schizothorax eurystomus|FFU194-21
Schizothorax eurystomus|FFU196-21
Schizothorax eurystomus|FFU173-21
Schizothorax eurystomus|FFU175-21
Schizothorax eurystomus|FFU183-21
Schizothorax eurystomus|FFU187-21
Schizothorax eurystomus|FFU191-21
Schizothorax eurystomus|FFU185-21
Schizothorax eurystomus|FFU189-21
Schizothorax eurystomus|FFU190-21
Schizothorax eurystomus|FFU192-21
Schizothorax eurystomus|FFU193-21
Schizothorax eurystomus|FFU195-21
Schizothorax eurystomus|FFU177-21
Schizothorax eurystomus|FFU179-21
Schizothorax eurystomus|FFU197-21
Schizothorax eurystomus|FFU186-21
Schizothorax eurystomus|FFU188-21
Schizothorax fedtschenkoi|FFU200-21
Schizothorax fedtschenkoi|FFU199-21

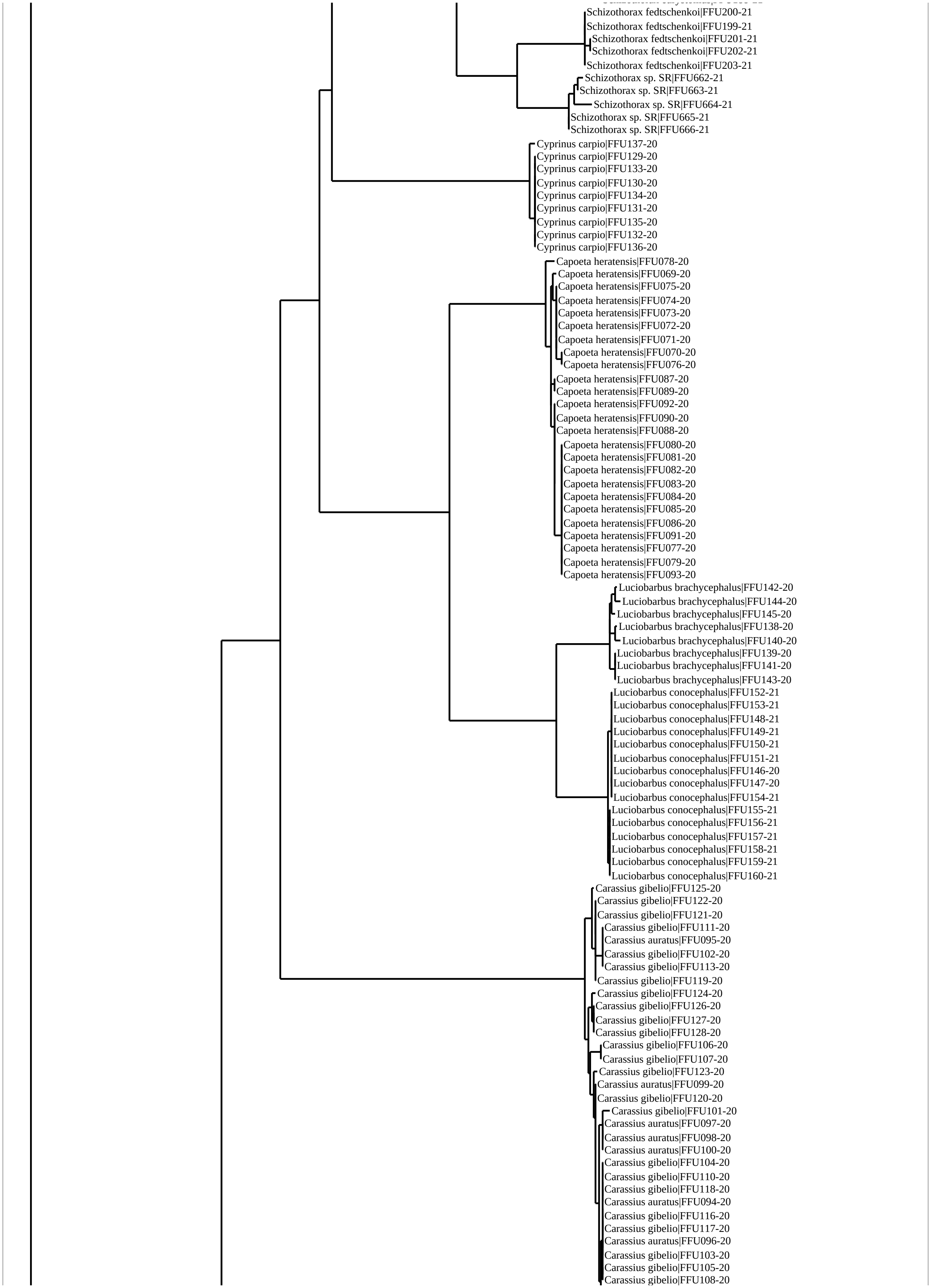
Schizothorax fedtschenkoi|FFU200-21
Schizothorax fedtschenkoi|FFU199-21
Schizothorax fedtschenkoi|FFU201-21
Schizothorax fedtschenkoi|FFU202-21
Schizothorax fedtschenkoi|FFU203-21
Schizothorax sp. SR|FFU662-21
Schizothorax sp. SR|FFU663-21
Schizothorax sp. SR|FFU664-21
Schizothorax sp. SR|FFU665-21
Schizothorax sp. SR|FFU666-21
Cyprinus carpio|FFU137-20
Cyprinus carpio|FFU129-20
Cyprinus carpio|FFU133-20
Cyprinus carpio|FFU130-20
Cyprinus carpio|FFU134-20
Cyprinus carpio|FFU131-20
Cyprinus carpio|FFU135-20
Cyprinus carpio|FFU132-20
Cyprinus carpio|FFU136-20
Capoeta heratensis|FFU078-20
Capoeta heratensis|FFU069-20
Capoeta heratensis|FFU075-20
Capoeta heratensis|FFU074-20
Capoeta heratensis|FFU073-20
Capoeta heratensis|FFU072-20
Capoeta heratensis|FFU071-20
Capoeta heratensis|FFU070-20
Capoeta heratensis|FFU076-20
Capoeta heratensis|FFU087-20
Capoeta heratensis|FFU089-20
Capoeta heratensis|FFU092-20
Capoeta heratensis|FFU090-20
Capoeta heratensis|FFU088-20
Capoeta heratensis|FFU080-20
Capoeta heratensis|FFU081-20
Capoeta heratensis|FFU082-20
Capoeta heratensis|FFU083-20
Capoeta heratensis|FFU084-20
Capoeta heratensis|FFU085-20
Capoeta heratensis|FFU086-20
Capoeta heratensis|FFU091-20
Capoeta heratensis|FFU077-20
Capoeta heratensis|FFU079-20
Capoeta heratensis|FFU093-20
Luciobarbus brachycephalus|FFU142-20
Luciobarbus brachycephalus|FFU144-20
Luciobarbus brachycephalus|FFU145-20
Luciobarbus brachycephalus|FFU138-20
Luciobarbus brachycephalus|FFU140-20
Luciobarbus brachycephalus|FFU139-20
Luciobarbus brachycephalus|FFU141-20
Luciobarbus brachycephalus|FFU143-20
Luciobarbus conocephalus|FFU152-21
Luciobarbus conocephalus|FFU153-21
Luciobarbus conocephalus|FFU148-21
Luciobarbus conocephalus|FFU149-21
Luciobarbus conocephalus|FFU150-21
Luciobarbus conocephalus|FFU151-21
Luciobarbus conocephalus|FFU146-20
Luciobarbus conocephalus|FFU147-20
Luciobarbus conocephalus|FFU154-21
Luciobarbus conocephalus|FFU155-21
Luciobarbus conocephalus|FFU156-21
Luciobarbus conocephalus|FFU157-21
Luciobarbus conocephalus|FFU158-21
Luciobarbus conocephalus|FFU159-21
Luciobarbus conocephalus|FFU160-21
Carassius gibelio|FFU125-20
Carassius gibelio|FFU122-20
Carassius gibelio|FFU121-20
Carassius gibelio|FFU111-20
Carassius auratus|FFU095-20
Carassius gibelio|FFU102-20
Carassius gibelio|FFU113-20
Carassius gibelio|FFU119-20
Carassius gibelio|FFU124-20
Carassius gibelio|FFU126-20
Carassius gibelio|FFU127-20
Carassius gibelio|FFU128-20
Carassius gibelio|FFU106-20
Carassius gibelio|FFU107-20
Carassius gibelio|FFU123-20
Carassius auratus|FFU099-20
Carassius gibelio|FFU120-20
Carassius gibelio|FFU101-20
Carassius auratus|FFU097-20
Carassius auratus|FFU098-20
Carassius auratus|FFU100-20
Carassius gibelio|FFU104-20
Carassius gibelio|FFU110-20
Carassius gibelio|FFU118-20
Carassius auratus|FFU094-20
Carassius gibelio|FFU116-20
Carassius gibelio|FFU117-20
Carassius auratus|FFU096-20
Carassius gibelio|FFU103-20
Carassius gibelio|FFU105-20
Carassius gibelio|FFU108-20

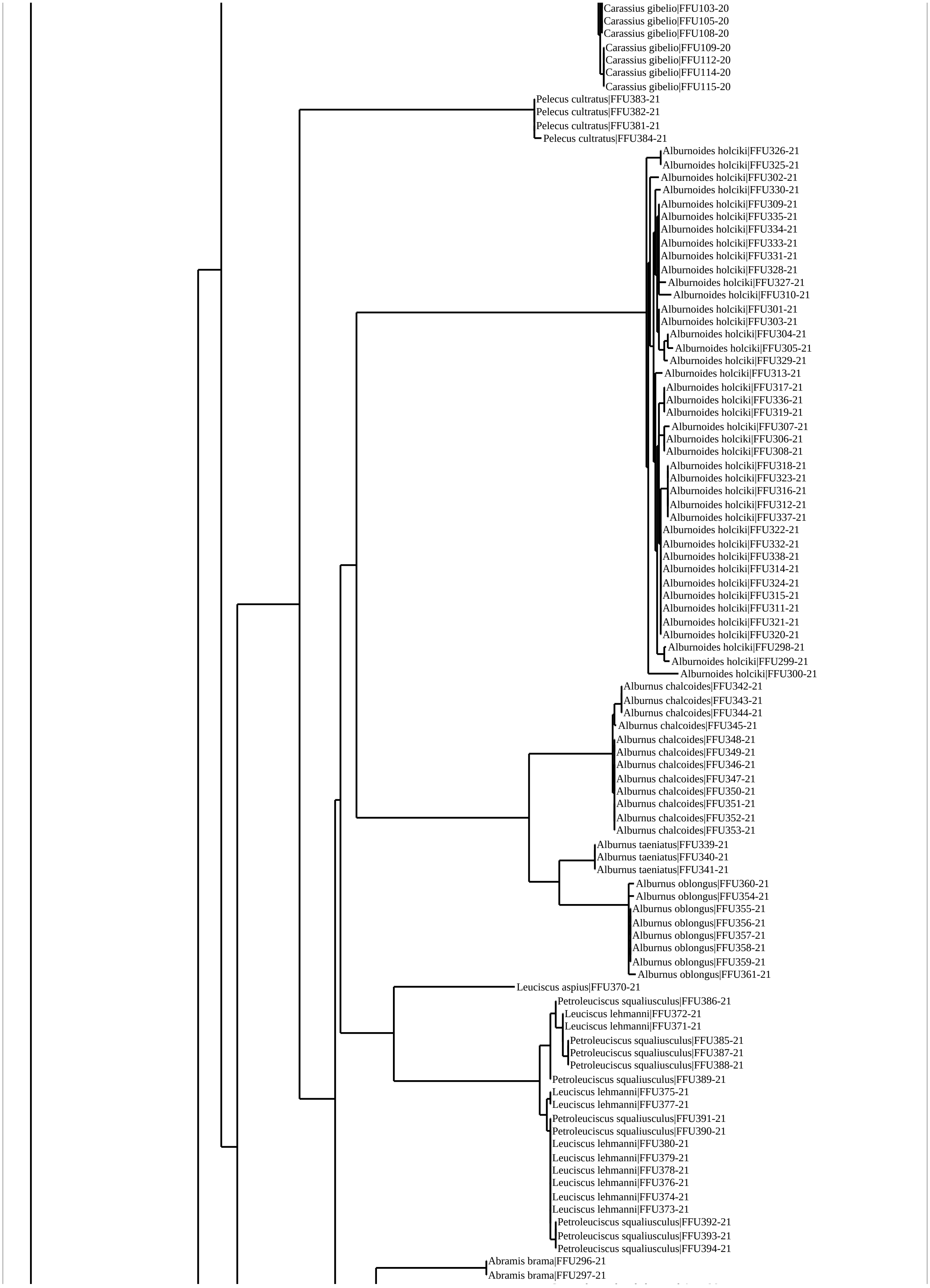
Carassius gibelio|FFU103-20
Carassius gibelio|FFU105-20
Carassius gibelio|FFU108-20
Carassius gibelio|FFU109-20
Carassius gibelio|FFU112-20
Carassius gibelio|FFU114-20
Carassius gibelio|FFU115-20
Pelecus cultratus|FFU383-21
Pelecus cultratus|FFU382-21
Pelecus cultratus|FFU381-21
Pelecus cultratus|FFU384-21
Alburnoides holciki|FFU326-21
Alburnoides holciki|FFU325-21
Alburnoides holciki|FFU302-21
Alburnoides holciki|FFU330-21
Alburnoides holciki|FFU309-21
Alburnoides holciki|FFU335-21
Alburnoides holciki|FFU334-21
Alburnoides holciki|FFU333-21
Alburnoides holciki|FFU331-21
Alburnoides holciki|FFU328-21
Alburnoides holciki|FFU327-21
Alburnoides holciki|FFU310-21
Alburnoides holciki|FFU301-21
Alburnoides holciki|FFU303-21
Alburnoides holciki|FFU304-21
Alburnoides holciki|FFU305-21
Alburnoides holciki|FFU329-21
Alburnoides holciki|FFU313-21
Alburnoides holciki|FFU317-21
Alburnoides holciki|FFU336-21
Alburnoides holciki|FFU319-21
Alburnoides holciki|FFU307-21
Alburnoides holciki|FFU306-21
Alburnoides holciki|FFU308-21
Alburnoides holciki|FFU318-21
Alburnoides holciki|FFU323-21
Alburnoides holciki|FFU316-21
Alburnoides holciki|FFU312-21
Alburnoides holciki|FFU337-21
Alburnoides holciki|FFU322-21
Alburnoides holciki|FFU332-21
Alburnoides holciki|FFU338-21
Alburnoides holciki|FFU314-21
Alburnoides holciki|FFU324-21
Alburnoides holciki|FFU315-21
Alburnoides holciki|FFU311-21
Alburnoides holciki|FFU321-21
Alburnoides holciki|FFU320-21
Alburnoides holciki|FFU298-21
Alburnoides holciki|FFU299-21
Alburnoides holciki|FFU300-21
Alburnus chalcoides|FFU342-21
Alburnus chalcoides|FFU343-21
Alburnus chalcoides|FFU344-21
Alburnus chalcoides|FFU345-21
Alburnus chalcoides|FFU348-21
Alburnus chalcoides|FFU349-21
Alburnus chalcoides|FFU346-21
Alburnus chalcoides|FFU347-21
Alburnus chalcoides|FFU350-21
Alburnus chalcoides|FFU351-21
Alburnus chalcoides|FFU352-21
Alburnus chalcoides|FFU353-21
Alburnus taeniatus|FFU339-21
Alburnus taeniatus|FFU340-21
Alburnus taeniatus|FFU341-21
Alburnus oblongus|FFU360-21
Alburnus oblongus|FFU354-21
Alburnus oblongus|FFU355-21
Alburnus oblongus|FFU356-21
Alburnus oblongus|FFU357-21
Alburnus oblongus|FFU358-21
Alburnus oblongus|FFU359-21
Alburnus oblongus|FFU361-21
Leuciscus aspius|FFU370-21
Petroleuciscus squaliusculus|FFU386-21
Leuciscus lehmanni|FFU372-21
Leuciscus lehmanni|FFU371-21
Petroleuciscus squaliusculus|FFU385-21
Petroleuciscus squaliusculus|FFU387-21
Petroleuciscus squaliusculus|FFU388-21
Petroleuciscus squaliusculus|FFU389-21
Leuciscus lehmanni|FFU375-21
Leuciscus lehmanni|FFU377-21
Petroleuciscus squaliusculus|FFU391-21
Petroleuciscus squaliusculus|FFU390-21
Leuciscus lehmanni|FFU380-21
Leuciscus lehmanni|FFU379-21
Leuciscus lehmanni|FFU378-21
Leuciscus lehmanni|FFU376-21
Leuciscus lehmanni|FFU374-21
Leuciscus lehmanni|FFU373-21
Petroleuciscus squaliusculus|FFU392-21
Petroleuciscus squaliusculus|FFU393-21
Petroleuciscus squaliusculus|FFU394-21
Abramis brama|FFU296-21
Abramis brama|FFU297-21

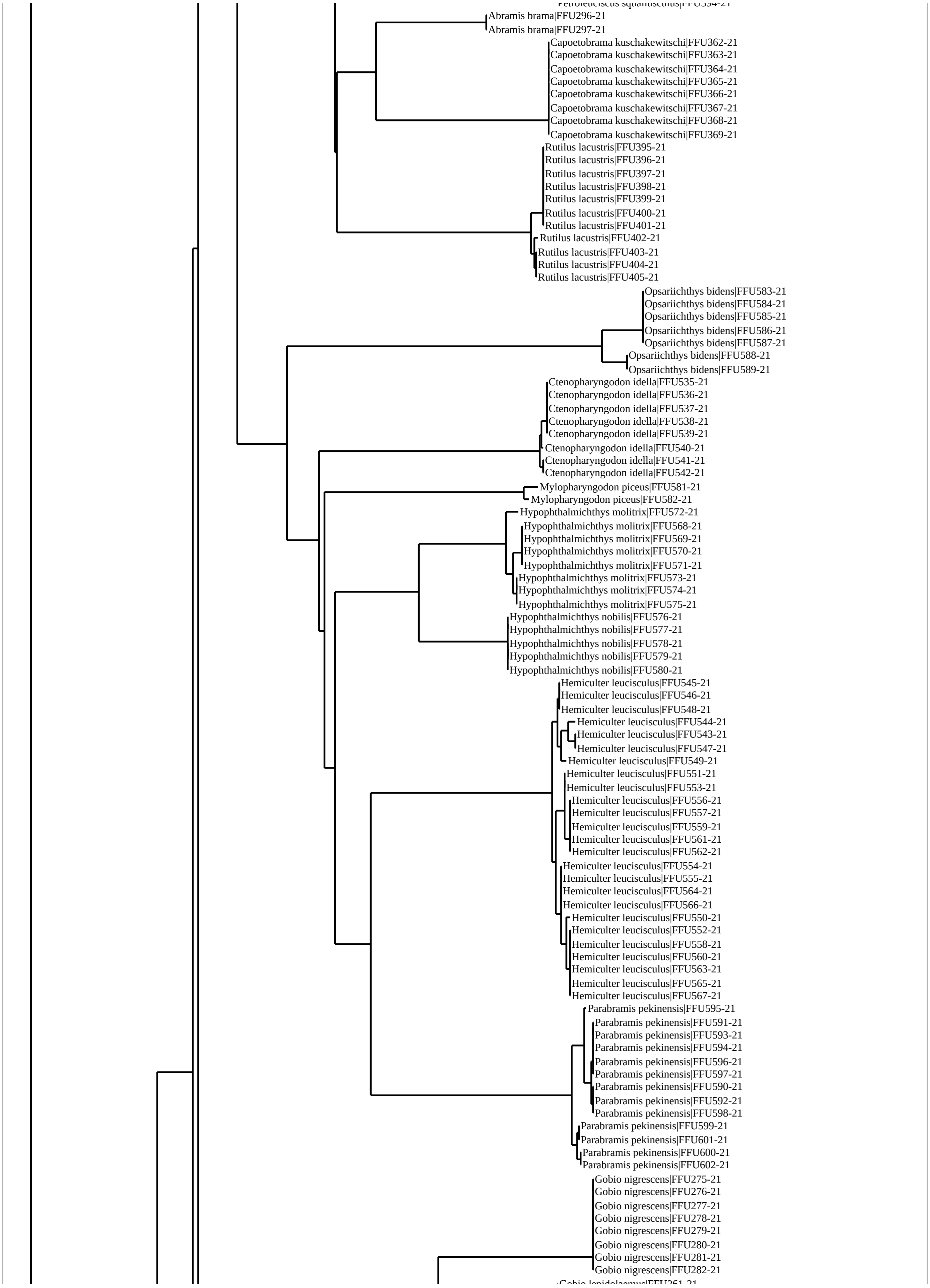
Abramis brama|FFU296-21
Abramis brama|FFU297-21
Capoetobrama kuschakewitschi|FFU362-21
Capoetobrama kuschakewitschi|FFU363-21
Capoetobrama kuschakewitschi|FFU364-21
Capoetobrama kuschakewitschi|FFU365-21
Capoetobrama kuschakewitschi|FFU366-21
Capoetobrama kuschakewitschi|FFU367-21
Capoetobrama kuschakewitschi|FFU368-21
Capoetobrama kuschakewitschi|FFU369-21
Rutilus lacustris|FFU395-21
Rutilus lacustris|FFU396-21
Rutilus lacustris|FFU397-21
Rutilus lacustris|FFU398-21
Rutilus lacustris|FFU399-21
Rutilus lacustris|FFU400-21
Rutilus lacustris|FFU401-21
Rutilus lacustris|FFU402-21
Rutilus lacustris|FFU403-21
Rutilus lacustris|FFU404-21
Rutilus lacustris|FFU405-21
Opsariichthys bidens|FFU583-21
Opsariichthys bidens|FFU584-21
Opsariichthys bidens|FFU585-21
Opsariichthys bidens|FFU586-21
Opsariichthys bidens|FFU587-21
Opsariichthys bidens|FFU588-21
Opsariichthys bidens|FFU589-21
Ctenopharyngodon idella|FFU535-21
Ctenopharyngodon idella|FFU536-21
Ctenopharyngodon idella|FFU537-21
Ctenopharyngodon idella|FFU538-21
Ctenopharyngodon idella|FFU539-21
Ctenopharyngodon idella|FFU540-21
Ctenopharyngodon idella|FFU541-21
Ctenopharyngodon idella|FFU542-21
Mylopharyngodon piceus|FFU581-21
Mylopharyngodon piceus|FFU582-21
Hypophthalmichthys molitrix|FFU572-21
Hypophthalmichthys molitrix|FFU568-21
Hypophthalmichthys molitrix|FFU569-21
Hypophthalmichthys molitrix|FFU570-21
Hypophthalmichthys molitrix|FFU571-21
Hypophthalmichthys molitrix|FFU573-21
Hypophthalmichthys molitrix|FFU574-21
Hypophthalmichthys molitrix|FFU575-21
Hypophthalmichthys nobilis|FFU576-21
Hypophthalmichthys nobilis|FFU577-21
Hypophthalmichthys nobilis|FFU578-21
Hypophthalmichthys nobilis|FFU579-21
Hypophthalmichthys nobilis|FFU580-21
Hemiculter leucisculus|FFU545-21
Hemiculter leucisculus|FFU546-21
Hemiculter leucisculus|FFU548-21
Hemiculter leucisculus|FFU544-21
Hemiculter leucisculus|FFU543-21
Hemiculter leucisculus|FFU547-21
Hemiculter leucisculus|FFU549-21
Hemiculter leucisculus|FFU551-21
Hemiculter leucisculus|FFU553-21
Hemiculter leucisculus|FFU556-21
Hemiculter leucisculus|FFU557-21
Hemiculter leucisculus|FFU559-21
Hemiculter leucisculus|FFU561-21
Hemiculter leucisculus|FFU562-21
Hemiculter leucisculus|FFU554-21
Hemiculter leucisculus|FFU555-21
Hemiculter leucisculus|FFU564-21
Hemiculter leucisculus|FFU566-21
Hemiculter leucisculus|FFU550-21
Hemiculter leucisculus|FFU552-21
Hemiculter leucisculus|FFU558-21
Hemiculter leucisculus|FFU560-21
Hemiculter leucisculus|FFU563-21
Hemiculter leucisculus|FFU565-21
Hemiculter leucisculus|FFU567-21
Parabramis pekinensis|FFU595-21
Parabramis pekinensis|FFU591-21
Parabramis pekinensis|FFU593-21
Parabramis pekinensis|FFU594-21
Parabramis pekinensis|FFU596-21
Parabramis pekinensis|FFU597-21
Parabramis pekinensis|FFU590-21
Parabramis pekinensis|FFU592-21
Parabramis pekinensis|FFU598-21
Parabramis pekinensis|FFU599-21
Parabramis pekinensis|FFU601-21
Parabramis pekinensis|FFU600-21
Parabramis pekinensis|FFU602-21
Gobio nigrescens|FFU275-21
Gobio nigrescens|FFU276-21
Gobio nigrescens|FFU277-21
Gobio nigrescens|FFU278-21
Gobio nigrescens|FFU279-21
Gobio nigrescens|FFU280-21
Gobio nigrescens|FFU281-21
Gobio nigrescens|FFU282-21

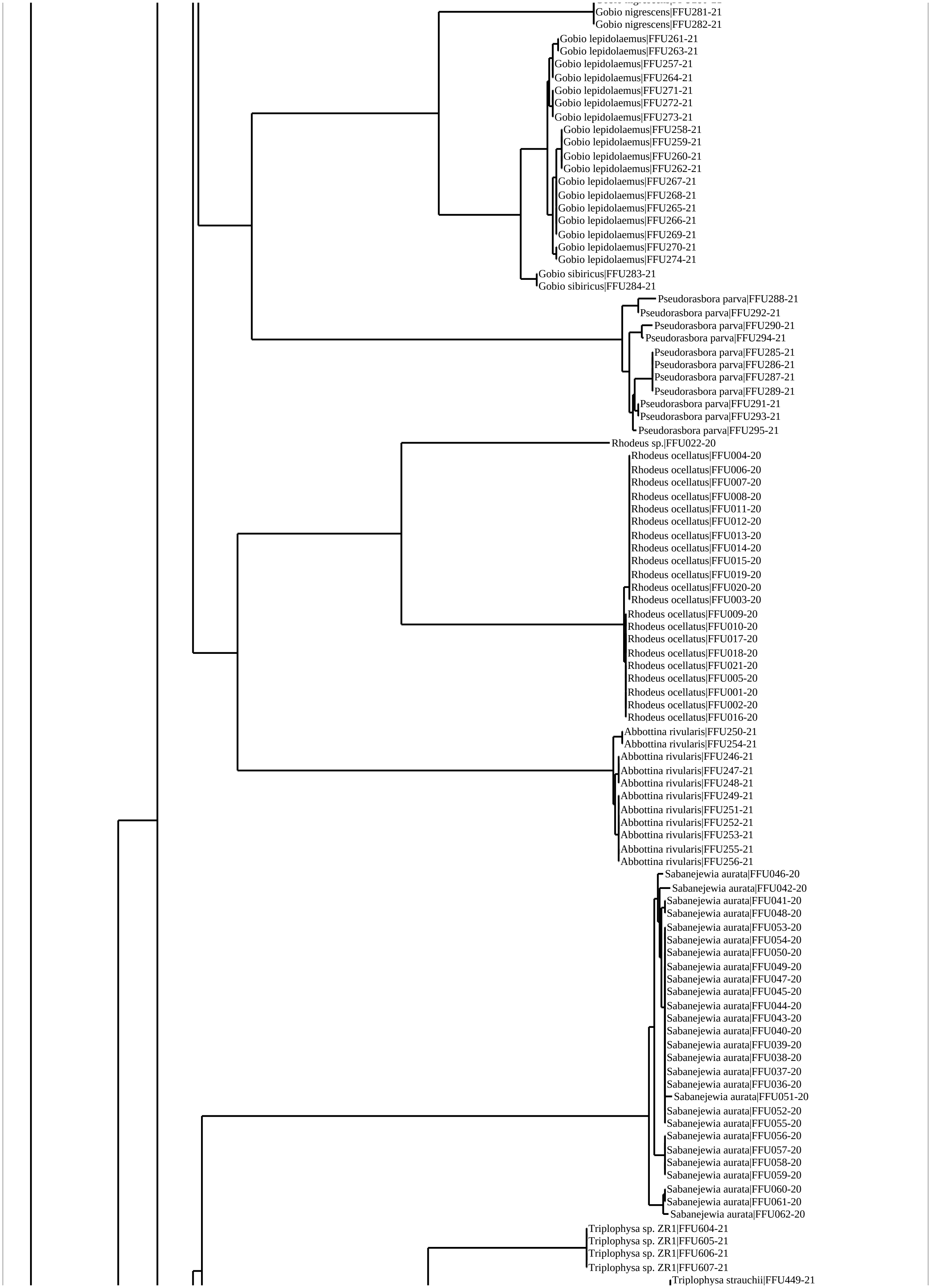
Gobio nigrescens|FFU281-21
Gobio nigrescens|FFU282-21
Gobio lepidolaemus|FFU261-21
Gobio lepidolaemus|FFU263-21
Gobio lepidolaemus|FFU257-21
Gobio lepidolaemus|FFU264-21
Gobio lepidolaemus|FFU271-21
Gobio lepidolaemus|FFU272-21
Gobio lepidolaemus|FFU273-21
Gobio lepidolaemus|FFU258-21
Gobio lepidolaemus|FFU259-21
Gobio lepidolaemus|FFU260-21
Gobio lepidolaemus|FFU262-21
Gobio lepidolaemus|FFU267-21
Gobio lepidolaemus|FFU268-21
Gobio lepidolaemus|FFU265-21
Gobio lepidolaemus|FFU266-21
Gobio lepidolaemus|FFU269-21
Gobio lepidolaemus|FFU270-21
Gobio lepidolaemus|FFU274-21
Gobio sibiricus|FFU283-21
Gobio sibiricus|FFU284-21
Pseudorasbora parva|FFU288-21
Pseudorasbora parva|FFU292-21
Pseudorasbora parva|FFU290-21
Pseudorasbora parva|FFU294-21
Pseudorasbora parva|FFU285-21
Pseudorasbora parva|FFU286-21
Pseudorasbora parva|FFU287-21
Pseudorasbora parva|FFU289-21
Pseudorasbora parva|FFU291-21
Pseudorasbora parva|FFU293-21
Pseudorasbora parva|FFU295-21
Rhodeus sp.|FFU022-20
Rhodeus ocellatus|FFU004-20
Rhodeus ocellatus|FFU006-20
Rhodeus ocellatus|FFU007-20
Rhodeus ocellatus|FFU008-20
Rhodeus ocellatus|FFU011-20
Rhodeus ocellatus|FFU012-20
Rhodeus ocellatus|FFU013-20
Rhodeus ocellatus|FFU014-20
Rhodeus ocellatus|FFU015-20
Rhodeus ocellatus|FFU019-20
Rhodeus ocellatus|FFU020-20
Rhodeus ocellatus|FFU003-20
Rhodeus ocellatus|FFU009-20
Rhodeus ocellatus|FFU010-20
Rhodeus ocellatus|FFU017-20
Rhodeus ocellatus|FFU018-20
Rhodeus ocellatus|FFU021-20
Rhodeus ocellatus|FFU005-20
Rhodeus ocellatus|FFU001-20
Rhodeus ocellatus|FFU002-20
Rhodeus ocellatus|FFU016-20
Abbottina rivularis|FFU250-21
Abbottina rivularis|FFU254-21
Abbottina rivularis|FFU246-21
Abbottina rivularis|FFU247-21
Abbottina rivularis|FFU248-21
Abbottina rivularis|FFU249-21
Abbottina rivularis|FFU251-21
Abbottina rivularis|FFU252-21
Abbottina rivularis|FFU253-21
Abbottina rivularis|FFU255-21
Abbottina rivularis|FFU256-21
Sabanejewia aurata|FFU046-20
Sabanejewia aurata|FFU042-20
Sabanejewia aurata|FFU041-20
Sabanejewia aurata|FFU048-20
Sabanejewia aurata|FFU053-20
Sabanejewia aurata|FFU054-20
Sabanejewia aurata|FFU050-20
Sabanejewia aurata|FFU049-20
Sabanejewia aurata|FFU047-20
Sabanejewia aurata|FFU045-20
Sabanejewia aurata|FFU044-20
Sabanejewia aurata|FFU043-20
Sabanejewia aurata|FFU040-20
Sabanejewia aurata|FFU039-20
Sabanejewia aurata|FFU038-20
Sabanejewia aurata|FFU037-20
Sabanejewia aurata|FFU036-20
Sabanejewia aurata|FFU051-20
Sabanejewia aurata|FFU052-20
Sabanejewia aurata|FFU055-20
Sabanejewia aurata|FFU056-20
Sabanejewia aurata|FFU057-20
Sabanejewia aurata|FFU058-20
Sabanejewia aurata|FFU059-20
Sabanejewia aurata|FFU060-20
Sabanejewia aurata|FFU061-20
Sabanejewia aurata|FFU062-20
Triplophysa sp. ZR1|FFU604-21
Triplophysa sp. ZR1|FFU605-21
Triplophysa sp. ZR1|FFU606-21
Triplophysa sp. ZR1|FFU607-21
Triplophysa strauchii|FFU449-21

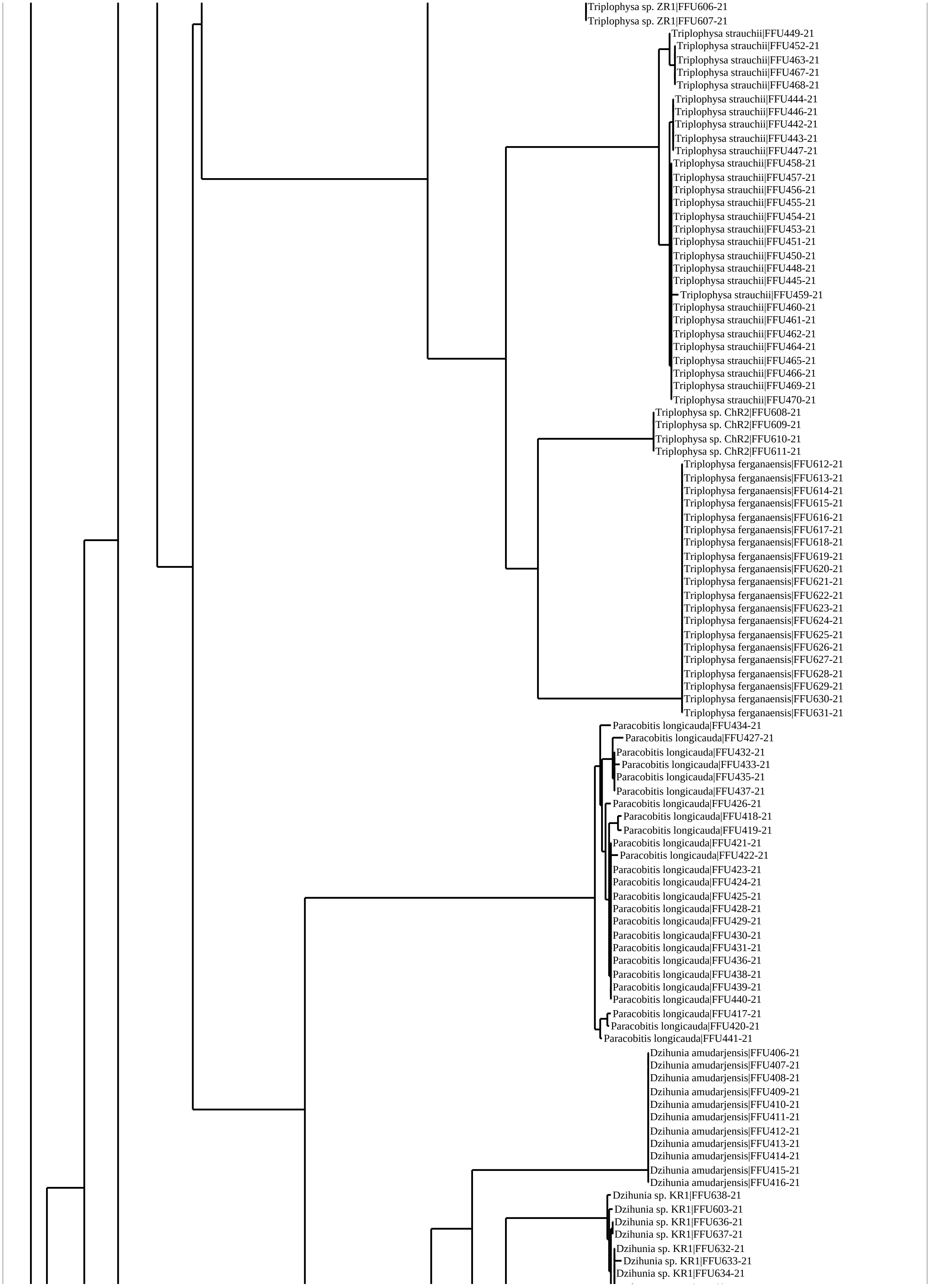
Triplophysa sp. ZR1|FFU606-21
Triplophysa sp. ZR1|FFU607-21
Triplophysa strauchii|FFU449-21
Triplophysa strauchii|FFU452-21
Triplophysa strauchii|FFU463-21
Triplophysa strauchii|FFU467-21
Triplophysa strauchii|FFU468-21
Triplophysa strauchii|FFU444-21
Triplophysa strauchii|FFU446-21
Triplophysa strauchii|FFU442-21
Triplophysa strauchii|FFU443-21
Triplophysa strauchii|FFU447-21
Triplophysa strauchii|FFU458-21
Triplophysa strauchii|FFU457-21
Triplophysa strauchii|FFU456-21
Triplophysa strauchii|FFU455-21
Triplophysa strauchii|FFU454-21
Triplophysa strauchii|FFU453-21
Triplophysa strauchii|FFU451-21
Triplophysa strauchii|FFU450-21
Triplophysa strauchii|FFU448-21
Triplophysa strauchii|FFU445-21
Triplophysa strauchii|FFU459-21
Triplophysa strauchii|FFU460-21
Triplophysa strauchii|FFU461-21
Triplophysa strauchii|FFU462-21
Triplophysa strauchii|FFU464-21
Triplophysa strauchii|FFU465-21
Triplophysa strauchii|FFU466-21
Triplophysa strauchii|FFU469-21
Triplophysa strauchii|FFU470-21
Triplophysa sp. ChR2|FFU608-21
Triplophysa sp. ChR2|FFU609-21
Triplophysa sp. ChR2|FFU610-21
Triplophysa sp. ChR2|FFU611-21
Triplophysa ferganaensis|FFU612-21
Triplophysa ferganaensis|FFU613-21
Triplophysa ferganaensis|FFU614-21
Triplophysa ferganaensis|FFU615-21
Triplophysa ferganaensis|FFU616-21
Triplophysa ferganaensis|FFU617-21
Triplophysa ferganaensis|FFU618-21
Triplophysa ferganaensis|FFU619-21
Triplophysa ferganaensis|FFU620-21
Triplophysa ferganaensis|FFU621-21
Triplophysa ferganaensis|FFU622-21
Triplophysa ferganaensis|FFU623-21
Triplophysa ferganaensis|FFU624-21
Triplophysa ferganaensis|FFU625-21
Triplophysa ferganaensis|FFU626-21
Triplophysa ferganaensis|FFU627-21
Triplophysa ferganaensis|FFU628-21
Triplophysa ferganaensis|FFU629-21
Triplophysa ferganaensis|FFU630-21
Triplophysa ferganaensis|FFU631-21
Paracobitis longicauda|FFU434-21
Paracobitis longicauda|FFU427-21
Paracobitis longicauda|FFU432-21
Paracobitis longicauda|FFU433-21
Paracobitis longicauda|FFU435-21
Paracobitis longicauda|FFU437-21
Paracobitis longicauda|FFU426-21
Paracobitis longicauda|FFU418-21
Paracobitis longicauda|FFU419-21
Paracobitis longicauda|FFU421-21
Paracobitis longicauda|FFU422-21
Paracobitis longicauda|FFU423-21
Paracobitis longicauda|FFU424-21
Paracobitis longicauda|FFU425-21
Paracobitis longicauda|FFU428-21
Paracobitis longicauda|FFU429-21
Paracobitis longicauda|FFU430-21
Paracobitis longicauda|FFU431-21
Paracobitis longicauda|FFU436-21
Paracobitis longicauda|FFU438-21
Paracobitis longicauda|FFU439-21
Paracobitis longicauda|FFU440-21
Paracobitis longicauda|FFU417-21
Paracobitis longicauda|FFU420-21
Paracobitis longicauda|FFU441-21
Dzihunia amudarjensis|FFU406-21
Dzihunia amudarjensis|FFU407-21
Dzihunia amudarjensis|FFU408-21
Dzihunia amudarjensis|FFU409-21
Dzihunia amudarjensis|FFU410-21
Dzihunia amudarjensis|FFU411-21
Dzihunia amudarjensis|FFU412-21
Dzihunia amudarjensis|FFU413-21
Dzihunia amudarjensis|FFU414-21
Dzihunia amudarjensis|FFU415-21
Dzihunia amudarjensis|FFU416-21
Dzihunia sp. KR1|FFU638-21
Dzihunia sp. KR1|FFU603-21
Dzihunia sp. KR1|FFU636-21
Dzihunia sp. KR1|FFU637-21
Dzihunia sp. KR1|FFU632-21
Dzihunia sp. KR1|FFU633-21
Dzihunia sp. KR1|FFU634-21

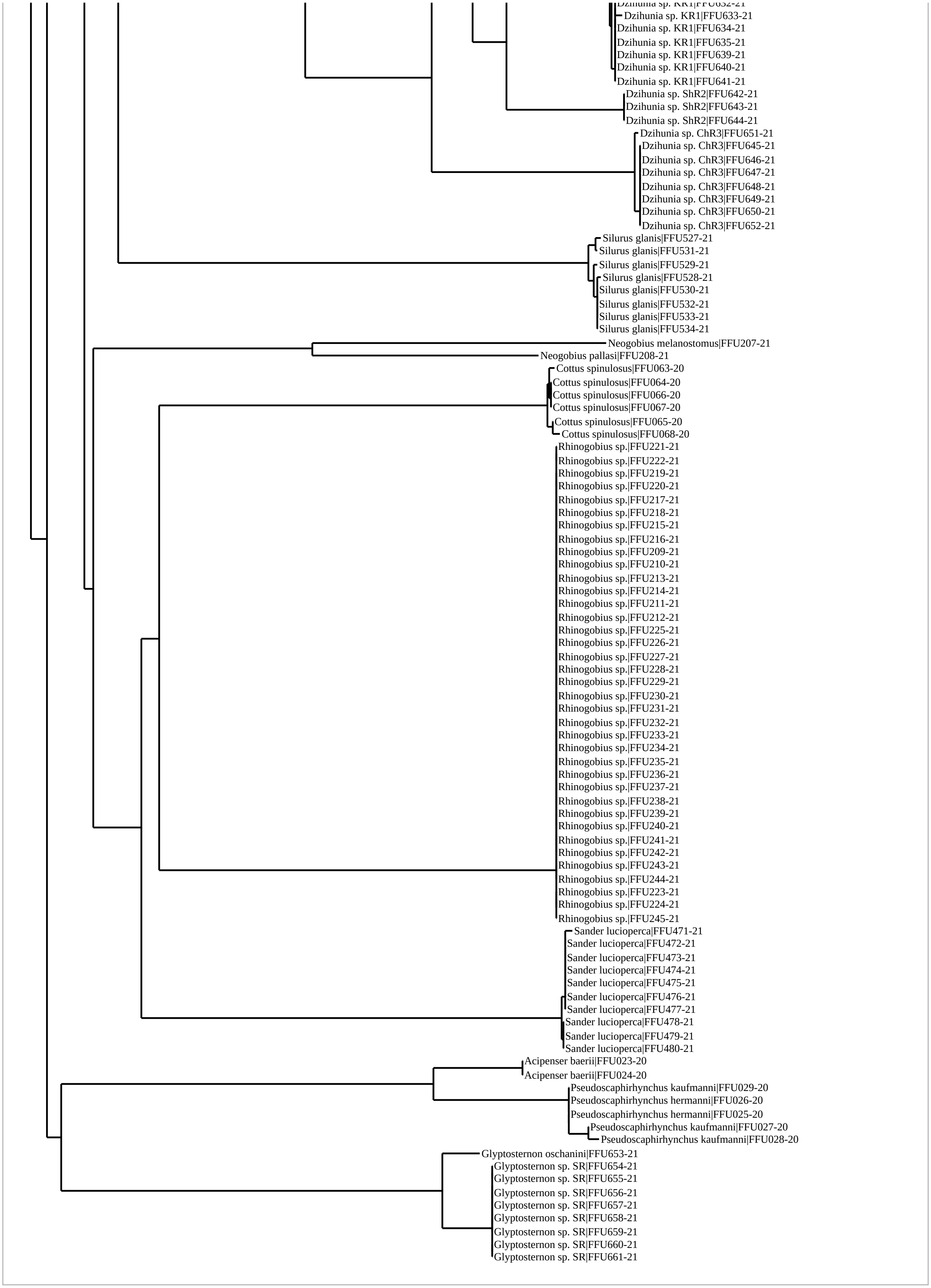
Dzihunia sp. KR1|FFU633-21
Dzihunia sp. KR1|FFU634-21
Dzihunia sp. KR1|FFU635-21
Dzihunia sp. KR1|FFU639-21
Dzihunia sp. KR1|FFU640-21
Dzihunia sp. KR1|FFU641-21
Dzihunia sp. ShR2|FFU642-21
Dzihunia sp. ShR2|FFU643-21
Dzihunia sp. ShR2|FFU644-21
Dzihunia sp. ChR3|FFU651-21
Dzihunia sp. ChR3|FFU645-21
Dzihunia sp. ChR3|FFU646-21
Dzihunia sp. ChR3|FFU647-21
Dzihunia sp. ChR3|FFU648-21
Dzihunia sp. ChR3|FFU649-21
Dzihunia sp. ChR3|FFU650-21
Dzihunia sp. ChR3|FFU652-21
Silurus glanis|FFU527-21
Silurus glanis|FFU531-21
Silurus glanis|FFU529-21
Silurus glanis|FFU528-21
Silurus glanis|FFU530-21
Silurus glanis|FFU532-21
Silurus glanis|FFU533-21
Silurus glanis|FFU534-21
Neogobius melanostomus|FFU207-21
Neogobius pallasi|FFU208-21
Cottus spinulosus|FFU063-20
Cottus spinulosus|FFU064-20
Cottus spinulosus|FFU066-20
Cottus spinulosus|FFU067-20
Cottus spinulosus|FFU065-20
Cottus spinulosus|FFU068-20
Rhinogobius sp.|FFU221-21
Rhinogobius sp.|FFU222-21
Rhinogobius sp.|FFU219-21
Rhinogobius sp.|FFU220-21
Rhinogobius sp.|FFU217-21
Rhinogobius sp.|FFU218-21
Rhinogobius sp.|FFU215-21
Rhinogobius sp.|FFU216-21
Rhinogobius sp.|FFU209-21
Rhinogobius sp.|FFU210-21
Rhinogobius sp.|FFU213-21
Rhinogobius sp.|FFU214-21
Rhinogobius sp.|FFU211-21
Rhinogobius sp.|FFU212-21
Rhinogobius sp.|FFU225-21
Rhinogobius sp.|FFU226-21
Rhinogobius sp.|FFU227-21
Rhinogobius sp.|FFU228-21
Rhinogobius sp.|FFU229-21
Rhinogobius sp.|FFU230-21
Rhinogobius sp.|FFU231-21
Rhinogobius sp.|FFU232-21
Rhinogobius sp.|FFU233-21
Rhinogobius sp.|FFU234-21
Rhinogobius sp.|FFU235-21
Rhinogobius sp.|FFU236-21
Rhinogobius sp.|FFU237-21
Rhinogobius sp.|FFU238-21
Rhinogobius sp.|FFU239-21
Rhinogobius sp.|FFU240-21
Rhinogobius sp.|FFU241-21
Rhinogobius sp.|FFU242-21
Rhinogobius sp.|FFU243-21
Rhinogobius sp.|FFU244-21
Rhinogobius sp.|FFU223-21
Rhinogobius sp.|FFU224-21
Rhinogobius sp.|FFU245-21
Sander lucioperca|FFU471-21
Sander lucioperca|FFU472-21
Sander lucioperca|FFU473-21
Sander lucioperca|FFU474-21
Sander lucioperca|FFU475-21
Sander lucioperca|FFU476-21
Sander lucioperca|FFU477-21
Sander lucioperca|FFU478-21
Sander lucioperca|FFU479-21
Sander lucioperca|FFU480-21
Acipenser baerii|FFU023-20
Acipenser baerii|FFU024-20
Pseudoscaphirhynchus kaufmanni|FFU029-20
Pseudoscaphirhynchus hermanni|FFU026-20
Pseudoscaphirhynchus hermanni|FFU025-20
Pseudoscaphirhynchus kaufmanni|FFU027-20
Pseudoscaphirhynchus kaufmanni|FFU028-20
Glyptosternon oschanini|FFU653-21
Glyptosternon sp. SR|FFU654-21
Glyptosternon sp. SR|FFU655-21
Glyptosternon sp. SR|FFU656-21
Glyptosternon sp. SR|FFU657-21
Glyptosternon sp. SR|FFU658-21
Glyptosternon sp. SR|FFU659-21
Glyptosternon sp. SR|FFU660-21
Glyptosternon sp. SR|FFU661-21
