## Supplemental Tables 1,2,3 for "Fish diversity in a doubly landlocked country - a description of the fish fauna of Uzbekistan using DNA barcoding"

| | |
|---|---|
| **Title** | **Fish diversity in a doubly landlocked country - a description of the fish fauna of Uzbekistan using DNA barcoding** |
| Authors | Bakhtiyor SHERALIEV[1,2], Zuogang PENG[1] |
| Affiliation | [1]Key Laboratory of Freshwater Fish Reproduction and Development (Ministry of Education), Southwest University, School of Life Sciences, Chongqing 400715, China |
| | [2]Faculty of Life Sciences, Fergana State University, Fergana, Uzbekistan |
| *Corresponding author | Professor Zuogang PENG, Ph.D.<br>E-mail: |
| | |

**TABLE S1.** Voucher metadata.

Identifications and metadata are subject to revision and may change

| No. | Sample ID | Voucher number | BOLD process ID | GenBank Acc. No. | Family | Species | Collected by | Identified by | Region | Drainage | River/stream/ hill/lake | Latitude | Longitude | Date |
|---|---|---|---|---|---|---|---|---|---|---|---|---|---|---|
| 1. | Rhodeus_ocellatus_261 | SWU19072019261 | FFU001-20 | MW649153 | Acheilognathidae | *Rhodeus ocellatus* | Sheraliev, BM | Sheraliev, BM | Surxondaryo | Amu Darya | Surkhan Darya | 37.255833 | 67.338611 | 19/07/2019 |
| 2. | Rhodeus_ocellatus_284 | SWU20072019284 | FFU002-20 | MW649154 | Acheilognathidae | *Rhodeus ocellatus* | Sheraliev, BM | Sheraliev, BM | Surxondaryo | Amu Darya | Amu Darya | 37.235555 | 67.660555 | 20/07/2019 |
| 3. | Rhodeus_ocellatus_353 | SWU22072019353 | FFU003-20 | MW649155 | Acheilognathidae | *Rhodeus ocellatus* | Sheraliev, BM | Sheraliev, BM | Surxondaryo | Amu Darya | Karatag river | 38.363055 | 68.072059 | 22/07/2019 |
| 4. | Rhodeus_ocellatus_354 | SWU22072019354 | FFU004-20 | MW649156 | Acheilognathidae | *Rhodeus ocellatus* | Sheraliev, BM | Sheraliev, BM | Surxondaryo | Amu Darya | Karatag river | 38.363055 | 68.072059 | 22/07/2019 |
| 5. | Rhodeus_ocellatus_355 | SWU22072019355 | FFU005-20 | MW649157 | Acheilognathidae | *Rhodeus ocellatus* | Sheraliev, BM | Sheraliev, BM | Surxondaryo | Amu Darya | Karatag river | 38.363055 | 68.072059 | 22/07/2019 |
| 6. | Rhodeus_ocellatus_356 | SWU22072019356 | FFU006-20 | MW649158 | Acheilognathidae | *Rhodeus ocellatus* | Sheraliev, BM | Sheraliev, BM | Surxondaryo | Amu Darya | Karatag river | 38.363055 | 68.072059 | 22/07/2019 |
| 7. | Rhodeus_ocellatus_357 | SWU22072019357 | FFU007-20 | MW649159 | Acheilognathidae | *Rhodeus ocellatus* | Sheraliev, BM | Sheraliev, BM | Surxondaryo | Amu Darya | Karatag river | 38.363055 | 68.072059 | 22/07/2019 |
| 8. | Rhodeus_ocellatus_390 | SWU24072019390 | FFU008-20 | MW649160 | Acheilognathidae | *Rhodeus ocellatus* | Sheraliev, BM | Sheraliev, BM | Surxondaryo | Amu Darya | Surkhan Darya | 37.823333 | 67.618611 | 24/07/2019 |
| 9. | Rhodeus_ocellatus_460 | SWU28072019460 | FFU009-20 | MW649161 | Acheilognathidae | *Rhodeus ocellatus* | Sheraliev, BM | Sheraliev, BM | Bukhara | Zeravshan | Zeravshan | 40.071012 | 64.778429 | 28/07/2019 |
| 10. | Rhodeus_ocellatus_461 | SWU28072019461 | FFU010-20 | MW649162 | Acheilognathidae | *Rhodeus ocellatus* | Sheraliev, BM | Sheraliev, BM | Bukhara | Zeravshan | Zeravshan | 40.071012 | 64.778429 | 28/07/2019 |
| 11. | Rhodeus_ocellatus_462 | SWU28072019462 | FFU011-20 | MW649163 | Acheilognathidae | *Rhodeus ocellatus* | Sheraliev, BM | Sheraliev, BM | Bukhara | Zeravshan | Zeravshan | 40.071012 | 64.778429 | 28/07/2019 |
| 12. | Rhodeus_ocellatus_550 | SWU09082019550 | FFU012-20 | MW649164 | Acheilognathidae | *Rhodeus ocellatus* | Sheraliev, BM | Sheraliev, BM | Namangan | Syr Darya | Kara Darya | 40.918813 | 71.828896 | 09/08/2019 |
| 13. | Rhodeus_ocellatus_570 | SWU09082019570 | FFU013-20 | MW649165 | Acheilognathidae | *Rhodeus ocellatus* | Sheraliev, BM | Sheraliev, BM | Namangan | Syr Darya | Kara Darya | 40.918813 | 71.828896 | 09/08/2019 |
| 14. | Rhodeus_ocellatus_572 | SWU09082019572 | FFU014-20 | MW649166 | Acheilognathidae | *Rhodeus ocellatus* | Sheraliev, BM | Sheraliev, BM | Namangan | Syr Darya | Kara Darya | 40.918813 | 71.828896 | 09/08/2019 |
| 15. | Rhodeus_ocellatus_573 | SWU09082019573 | FFU015-20 | MW649167 | Acheilognathidae | *Rhodeus ocellatus* | Sheraliev, BM | Sheraliev, BM | Namangan | Syr Darya | Kara Darya | 40.918813 | 71.828896 | 09/08/2019 |
| 16. | Rhodeus_ocellatus_585 | SWU09082019585 | FFU016-20 | MW649168 | Acheilognathidae | *Rhodeus ocellatus* | Sheraliev, BM | Sheraliev, BM | Namangan | Syr Darya | Naryn River | 40.940911 | 71.853631 | 09/08/2019 |
| 17. | Rhodeus_ocellatus_614 | SWU18082019614 | FFU017-20 | MW649169 | Acheilognathidae | *Rhodeus ocellatus* | Sheraliev, BM | Sheraliev, BM | Tashkent | Syr Darya | Chirchik River | 41.274444 | 69.389166 | 18/08/2019 |
| 18. | Rhodeus_ocellatus_615 | SWU18082019615 | FFU018-20 | MW649170 | Acheilognathidae | *Rhodeus ocellatus* | Sheraliev, BM | Sheraliev, BM | Tashkent | Syr Darya | Chirchik River | 41.274444 | 69.389166 | 18/08/2019 |
| 19. | Rhodeus_ocellatus_616 | SWU18082019616 | FFU019-20 | MW649171 | Acheilognathidae | *Rhodeus ocellatus* | Sheraliev, BM | Sheraliev, BM | Tashkent | Syr Darya | Chirchik River | 41.274444 | 69.389166 | 18/08/2019 |
| 20. | Rhodeus_ocellatus_712 | SWU16052020712 | FFU020-20 | MW649172 | Acheilognathidae | *Rhodeus ocellatus* | Rozimov, A | Sheraliev, BM | Khorazm | Amu Darya | Unnamed stream | 41.540222 | 60.166666 | 16/05/2020 |
| 21. | Rhodeus_ocellatus_713 | SWU17052020713 | FFU021-20 | MW649173 | Acheilognathidae | *Rhodeus ocellatus* | Rozimov, A | Sheraliev, BM | Khorazm | Amu Darya | Unnamed stream | 41.540222 | 60.166666 | 17/05/2020 |
| 22. | Rhodeus_sp_571 | SWU09082019571 | FFU022-20 | MW649174 | Acheilognathidae | *Rhodeus* sp. | Sheraliev, BM | Sheraliev, BM | Namangan | Syr Darya | Kara Darya | 40.918813 | 71.828896 | 09/08/2019 |
| 23. | Acipenser_baerii_708 | FSU28122019708 | FFU023-20 | MW649175 | Acipenseridae | *Acipenser baerii* | Rozimov, A | Sheraliev, BM | Khorazm | Amu Darya | Amu Darya | 41.603361 | 60.790055 | 28/12/2019 |
| 24. | Acipenser_baerii_709 | FSU28122019709 | FFU024-20 | MW649176 | Acipenseridae | *Acipenser baerii* | Rozimov, A | Sheraliev, BM | Khorazm | Amu Darya | Amu Darya | 41.603361 | 60.790055 | 28/12/2019 |
| 25. | Pseudoscaphirhynchus_hermanni_706 | FSU19012020706 | FFU025-20 | MW649177 | Acipenseridae | *Pseudoscaphirhynchus hermanni* | Rozimov, A | Sheraliev, BM | Khorazm | Amu Darya | Amu Darya | 41.489638 | 60.955777 | 19/01/2020 |
| 26. | Pseudoscaphirhynchus_hermanni_707 | FSU21012020707 | FFU026-20 | MW649178 | Acipenseridae | *Pseudoscaphirhynchus hermanni* | Rozimov, A | Sheraliev, BM | Khorazm | Amu Darya | Amu Darya | 41.489638 | 60.955777 | 21/01/2020 |
| 27. | Pseudoscaphirhynchus_kaufmanni_190 | FSU19072018190 | FFU027-20 | MW649179 | Acipenseridae | *Pseudoscaphirhynchus kaufmanni* | Sheraliev, BM | Sheraliev, BM | Khorazm | Amu Darya | Amu Darya | 41.466583 | 60.961305 | 19/07/2018 |
| 28. | Pseudoscaphirhynchus_kaufmanni_678 | FSU21082019678 | FFU028-20 | MW649180 | Acipenseridae | *Pseudoscaphirhynchus kaufmanni* | Rozimov, A | Sheraliev, BM | Khorazm | Amu Darya | Amu Darya | 41.344972 | 61.233583 | 21/08/2019 |
| 29. | Pseudoscaphirhynchus_kaufmanni_710 | FSU21052020710 | FFU029-20 | MW649181 | Acipenseridae | *Pseudoscaphirhynchus kaufmanni* | Rozimov, A | Sheraliev, BM | Khorazm | Amu Darya | Amu Darya | 41.660805 | 60.689305 | 21/05/2020 |
| 30. | Channa_argus_014 | SWU14082016014 | FFU030-20 | MW649182 | Channidae | *Channa argus* | Sheraliev, BM | Sheraliev, BM | Surxondaryo | Amu Darya | South Surkhan R. | 37.854013 | 67.628373 | 14/08/2016 |
| 31. | Channa_argus_092 | SWU23012018092 | FFU031-20 | MW649183 | Channidae | *Channa argus* | Sheraliev, BM | Sheraliev, BM | Fergana | Syr Darya | Local fish market | | | 23/01/2018 |
| 32. | Channa_argus_093 | SWU23012018093 | FFU032-20 | MW649184 | Channidae | *Channa argus* | Sheraliev, BM | Sheraliev, BM | Fergana | Syr Darya | Local fish market | | | 23/01/2018 |
| 33. | Channa_argus_094 | SWU23012018094 | FFU033-20 | MW649185 | Channidae | *Channa argus* | Sheraliev, BM | Sheraliev, BM | Fergana | Syr Darya | Local fish market | | | 23/01/2018 |
| 34. | Channa_argus_095 | SWU23012018095 | FFU034-20 | MW649186 | Channidae | *Channa argus* | Sheraliev, BM | Sheraliev, BM | Fergana | Syr Darya | Local fish market | | | 23/01/2018 |
| 35. | Channa_argus_096 | SWU23012018096 | FFU035-20 | MW649187 | Channidae | *Channa argus* | Sheraliev, BM | Sheraliev, BM | Fergana | Syr Darya | Local fish market | | | 23/01/2018 |
| 36. | Sabanejewia_aurata_253 | SWU19072019253 | FFU036-20 | MW649188 | Cobitidae | *Sabanejewia aurata* | Sheraliev, BM | Sheraliev, BM | Surxondaryo | Amu Darya | Surkhan Darya | 37.255833 | 67.338611 | 19/07/2019 |
| 37. | Sabanejewia_aurata_254 | SWU19072019254 | FFU037-20 | MW649189 | Cobitidae | *Sabanejewia aurata* | Sheraliev, BM | Sheraliev, BM | Surxondaryo | Amu Darya | Surkhan Darya | 37.255833 | 67.338611 | 19/07/2019 |
| 38. | Sabanejewia_aurata_255 | SWU19072019255 | FFU038-20 | MW649190 | Cobitidae | *Sabanejewia aurata* | Sheraliev, BM | Sheraliev, BM | Surxondaryo | Amu Darya | Surkhan Darya | 37.255833 | 67.338611 | 19/07/2019 |
| 39. | Sabanejewia_aurata_256 | SWU19072019256 | FFU039-20 | MW649191 | Cobitidae | *Sabanejewia aurata* | Sheraliev, BM | Sheraliev, BM | Surxondaryo | Amu Darya | Surkhan Darya | 37.255833 | 67.338611 | 19/07/2019 |
| 40. | Sabanejewia_aurata_300 | SWU21072019300 | FFU040-20 | MW649192 | Cobitidae | *Sabanejewia aurata* | Sheraliev, BM | Sheraliev, BM | Surxondaryo | Amu Darya | Surkhan Darya | 37.238888 | 67.329722 | 21/07/2019 |
| 41. | Sabanejewia_aurata_301 | SWU21072019301 | FFU041-20 | MW649193 | Cobitidae | *Sabanejewia aurata* | Sheraliev, BM | Sheraliev, BM | Surxondaryo | Amu Darya | Surkhan Darya | 37.238888 | 67.329722 | 21/07/2019 |
| 42. | Sabanejewia_aurata_302 | SWU21072019302 | FFU042-20 | MW649194 | Cobitidae | *Sabanejewia aurata* | Sheraliev, BM | Sheraliev, BM | Surxondaryo | Amu Darya | Surkhan Darya | 37.238888 | 67.329722 | 21/07/2019 |
| 43. | Sabanejewia_aurata_308 | SWU21072019308 | FFU043-20 | MW649195 | Cobitidae | *Sabanejewia aurata* | Sheraliev, BM | Sheraliev, BM | Surxondaryo | Amu Darya | Surkhan Darya | 37.238888 | 67.329722 | 21/07/2019 |
| 44. | Sabanejewia_aurata_326 | SWU22072019326 | FFU044-20 | MW649196 | Cobitidae | *Sabanejewia aurata* | Sheraliev, BM | Sheraliev, BM | Surxondaryo | Amu Darya | Karatag river | 38.363055 | 68.072059 | 22/07/2019 |
| 45. | Sabanejewia_aurata_327 | SWU22072019327 | FFU045-20 | MW649197 | Cobitidae | *Sabanejewia aurata* | Sheraliev, BM | Sheraliev, BM | Surxondaryo | Amu Darya | Karatag river | 38.363055 | 68.072059 | 22/07/2019 |
| 46. | Sabanejewia_aurata_328 | SWU22072019328 | FFU046-20 | MW649198 | Cobitidae | *Sabanejewia aurata* | Sheraliev, BM | Sheraliev, BM | Surxondaryo | Amu Darya | Karatag river | 38.363055 | 68.072059 | 22/07/2019 |
| 47. | Sabanejewia_aurata_329 | SWU22072019329 | FFU047-20 | MW649199 | Cobitidae | *Sabanejewia aurata* | Sheraliev, BM | Sheraliev, BM | Surxondaryo | Amu Darya | Karatag river | 38.363055 | 68.072059 | 22/07/2019 |
| 48. | Sabanejewia_aurata_330 | SWU22072019330 | FFU048-20 | MW649200 | Cobitidae | *Sabanejewia aurata* | Sheraliev, BM | Sheraliev, BM | Surxondaryo | Amu Darya | Karatag river | 38.363055 | 68.072059 | 22/07/2019 |
| 49. | Sabanejewia_aurata_384 | SWU23072019384 | FFU049-20 | MW649201 | Cobitidae | *Sabanejewia aurata* | Sheraliev, BM | Sheraliev, BM | Surxondaryo | Amu Darya | Tupalang River | 38.318966 | 68.006169 | 23/07/2019 |
| 50. | Sabanejewia_aurata_385 | SWU23072019385 | FFU050-20 | MW649202 | Cobitidae | *Sabanejewia aurata* | Sheraliev, BM | Sheraliev, BM | Surxondaryo | Amu Darya | Tupalang River | 38.318966 | 68.006169 | 23/07/2019 |
| 51. | Sabanejewia_aurata_386 | SWU23072019386 | FFU051-20 | MW649203 | Cobitidae | *Sabanejewia aurata* | Sheraliev, BM | Sheraliev, BM | Surxondaryo | Amu Darya | Tupalang River | 38.318966 | 68.006169 | 23/07/2019 |
| 52. | Sabanejewia_aurata_387 | SWU23072019387 | FFU052-20 | MW649204 | Cobitidae | *Sabanejewia aurata* | Sheraliev, BM | Sheraliev, BM | Surxondaryo | Amu Darya | Tupalang River | 38.318966 | 68.006169 | 23/07/2019 |
| 53. | Sabanejewia_aurata_388 | SWU23072019388 | FFU053-20 | MW649205 | Cobitidae | *Sabanejewia aurata* | Sheraliev, BM | Sheraliev, BM | Surxondaryo | Amu Darya | Tupalang River | 38.318966 | 68.006169 | 23/07/2019 |
| 54. | Sabanejewia_aurata_403 | SWU25072019403 | FFU054-20 | MW649206 | Cobitidae | *Sabanejewia aurata* | Sheraliev, BM | Sheraliev, BM | Surxondaryo | Amu Darya | Sherobod river | 37.747222 | 66.996666 | 25/07/2019 |
| 55. | Sabanejewia_aurata_416 | SWU27072019416 | FFU055-20 | MW649207 | Cobitidae | *Sabanejewia aurata* | Sheraliev, BM | Sheraliev, BM | Bukhara | Zeravshan | Echkiliksoy | 40.267551 | 64.743333 | 27/07/2019 |
| 56. | Sabanejewia_aurata_507 | SWU02082019507 | FFU056-20 | MW649208 | Cobitidae | *Sabanejewia aurata* | Sheraliev, BM | Sheraliev, BM | Samarkand | Zeravshan | Zeravshan | 39.676388 | 67.076111 | 02/08/2019 |
| 57. | Sabanejewia_aurata_508 | SWU02082019508 | FFU057-20 | MW649209 | Cobitidae | *Sabanejewia aurata* | Sheraliev, BM | Sheraliev, BM | Samarkand | Zeravshan | Zeravshan | 39.676388 | 67.076111 | 02/08/2019 |
| 58. | Sabanejewia_aurata_509 | SWU02082019509 | FFU058-20 | MW649210 | Cobitidae | *Sabanejewia aurata* | Sheraliev, BM | Sheraliev, BM | Samarkand | Zeravshan | Zeravshan | 39.676388 | 67.076111 | 02/08/2019 |
| 59. | Sabanejewia_aurata_510 | SWU02082019510 | FFU059-20 | MW649211 | Cobitidae | *Sabanejewia aurata* | Sheraliev, BM | Sheraliev, BM | Samarkand | Zeravshan | Zeravshan | 39.676388 | 67.076111 | 02/08/2019 |
| 60. | Sabanejewia_aurata_610 | SWU18082019610 | FFU060-20 | MW649212 | Cobitidae | *Sabanejewia aurata* | Sheraliev, BM | Sheraliev, BM | Tashkent | Syr Darya | Chirchik River | 41.274444 | 69.389166 | 18/08/2019 |
| 61. | Sabanejewia_aurata_611 | SWU18082019611 | FFU061-20 | MW649213 | Cobitidae | *Sabanejewia aurata* | Sheraliev, BM | Sheraliev, BM | Tashkent | Syr Darya | Chirchik River | 41.274444 | 69.389166 | 18/08/2019 |
| 62. | Sabanejewia_aurata_612 | SWU18082019612 | FFU062-20 | MW649214 | Cobitidae | *Sabanejewia aurata* | Sheraliev, BM | Sheraliev, BM | Tashkent | Syr Darya | Chirchik River | 41.274444 | 69.389166 | 18/08/2019 |
| 63. | Cottus_spinulosus_553 | SWU09082019553 | FFU063-20 | MW649215 | Cottidae | *Cottus spinulosus* | Sheraliev, BM | Sheraliev, BM | Namangan | Syr Darya | Kara Darya | 40.918813 | 71.828896 | 09/08/2019 |
| 64. | Cottus_spinulosus_554 | SWU09082019554 | FFU064-20 | MW649216 | Cottidae | *Cottus spinulosus* | Sheraliev, BM | Sheraliev, BM | Namangan | Syr Darya | Kara Darya | 40.918813 | 71.828896 | 09/08/2019 |

| | | | | | | | | | | | | | | |
|---|---|---|---|---|---|---|---|---|---|---|---|---|---|---|
| 65. | Cottus_spinulosus_555 | SWU09082019555 | FFU065-20 | MW649217 | Cottidae | *Cottus spinulosus* | Sheraliev, BM | Sheraliev, BM | Namangan | Syr Darya | Kara Darya | 40.918813 | 71.828896 | 09/08/2019 |
| 66. | Cottus_spinulosus_556 | SWU09082019556 | FFU066-20 | MW649218 | Cottidae | *Cottus spinulosus* | Sheraliev, BM | Sheraliev, BM | Namangan | Syr Darya | Kara Darya | 40.918813 | 71.828896 | 09/08/2019 |
| 67. | Cottus_spinulosus_557 | SWU09082019557 | FFU067-20 | MW649219 | Cottidae | *Cottus spinulosus* | Sheraliev, BM | Sheraliev, BM | Namangan | Syr Darya | Kara Darya | 40.918813 | 71.828896 | 09/08/2019 |
| 68. | Cottus_spinulosus_558 | SWU09082019558 | FFU068-20 | MW649220 | Cottidae | *Cottus spinulosus* | Sheraliev, BM | Sheraliev, BM | Namangan | Syr Darya | Kara Darya | 40.918813 | 71.828896 | 09/08/2019 |
| 69. | Capoeta_heratensis_008 | SWU10082016008 | FFU069-20 | MW649221 | Cyprinidae | *Capoeta heratensis* | Sheraliev, BM | Sheraliev, BM | Surxondaryo | Amu Darya | Tupalang river | 38.399802 | 67.944766 | 10/08/2016 |
| 70. | Capoeta_heratensis_145 | SWU16072018145 | FFU070-20 | MW649222 | Cyprinidae | *Capoeta heratensis* | Sheraliev, BM | Sheraliev, BM | Bukhara | Zeravshan | Zeravshan | 40.146944 | 64.891666 | 16/07/2018 |
| 71. | Capoeta_heratensis_146 | SWU16072018146 | FFU071-20 | MW649223 | Cyprinidae | *Capoeta heratensis* | Sheraliev, BM | Sheraliev, BM | Bukhara | Zeravshan | Zeravshan | 40.146944 | 64.891666 | 16/07/2018 |
| 72. | Capoeta_heratensis_147 | SWU16072018147 | FFU072-20 | MW649224 | Cyprinidae | *Capoeta heratensis* | Sheraliev, BM | Sheraliev, BM | Bukhara | Zeravshan | Zeravshan | 40.146944 | 64.891666 | 16/07/2018 |
| 73. | Capoeta_heratensis_148 | SWU16072018148 | FFU073-20 | MW649225 | Cyprinidae | *Capoeta heratensis* | Sheraliev, BM | Sheraliev, BM | Bukhara | Zeravshan | Zeravshan | 40.146944 | 64.891666 | 16/07/2018 |
| 74. | Capoeta_heratensis_149 | SWU16072018149 | FFU074-20 | MW649226 | Cyprinidae | *Capoeta heratensis* | Sheraliev, BM | Sheraliev, BM | Bukhara | Zeravshan | Zeravshan | 40.146944 | 64.891666 | 16/07/2018 |
| 75. | Capoeta_heratensis_169 | SWU17072018169 | FFU075-20 | MW649227 | Cyprinidae | *Capoeta heratensis* | Sheraliev, BM | Sheraliev, BM | Bukhara | Zeravshan | Dzhalvan canal | 40.149666 | 64.885400 | 17/07/2018 |
| 76. | Capoeta_heratensis_170 | SWU17072018170 | FFU076-20 | MW649228 | Cyprinidae | *Capoeta heratensis* | Sheraliev, BM | Sheraliev, BM | Bukhara | Zeravshan | Dzhalvan canal | 40.149666 | 64.885400 | 17/07/2018 |
| 77. | Capoeta_heratensis_258 | SWU19072019258 | FFU077-20 | MW649229 | Cyprinidae | *Capoeta heratensis* | Sheraliev, BM | Sheraliev, BM | Surxondaryo | Amu Darya | Surkhan Darya | 37.255833 | 67.338611 | 19/07/2019 |
| 78. | Capoeta_heratensis_259 | SWU19072019259 | FFU078-20 | MW649230 | Cyprinidae | *Capoeta heratensis* | Sheraliev, BM | Sheraliev, BM | Surxondaryo | Amu Darya | Surkhan Darya | 37.255833 | 67.338611 | 19/07/2019 |
| 79. | Capoeta_heratensis_260 | SWU19072019260 | FFU079-20 | MW649231 | Cyprinidae | *Capoeta heratensis* | Sheraliev, BM | Sheraliev, BM | Surxondaryo | Amu Darya | Surkhan Darya | 37.255833 | 67.338611 | 19/07/2019 |
| 80. | Capoeta_heratensis_306 | SWU21072019307 | FFU080-20 | MW649232 | Cyprinidae | *Capoeta heratensis* | Sheraliev, BM | Sheraliev, BM | Surxondaryo | Amu Darya | Surkhan Darya | 37.238888 | 67.329722 | 21/07/2019 |
| 81. | Capoeta_heratensis_320 | SWU21072019320 | FFU081-20 | MW649233 | Cyprinidae | *Capoeta heratensis* | Sheraliev, BM | Sheraliev, BM | Surxondaryo | Amu Darya | Surkhan Darya | 37.238888 | 67.329722 | 21/07/2019 |
| 82. | Capoeta_heratensis_322 | SWU21072019322 | FFU082-20 | MW649234 | Cyprinidae | *Capoeta heratensis* | Sheraliev, BM | Sheraliev, BM | Surxondaryo | Amu Darya | Surkhan Darya | 37.238888 | 67.329722 | 21/07/2019 |
| 83. | Capoeta_heratensis_404 | SWU25072019404 | FFU083-20 | MW649235 | Cyprinidae | *Capoeta heratensis* | Sheraliev, BM | Sheraliev, BM | Surxondaryo | Amu Darya | Sherobod river | 37.747222 | 66.996666 | 25/07/2019 |
| 84. | Capoeta_heratensis_412 | SWU25072019412 | FFU084-20 | MW649236 | Cyprinidae | *Capoeta heratensis* | Sheraliev, BM | Sheraliev, BM | Surxondaryo | Amu Darya | Sherobod river | 37.747222 | 66.996666 | 25/07/2019 |
| 85. | Capoeta_heratensis_413 | SWU25072019413 | FFU085-20 | MW649237 | Cyprinidae | *Capoeta heratensis* | Sheraliev, BM | Sheraliev, BM | Surxondaryo | Amu Darya | Sherobod river | 37.747222 | 66.996666 | 25/07/2019 |
| 86. | Capoeta_heratensis_414 | SWU25072019414 | FFU086-20 | MW649238 | Cyprinidae | *Capoeta heratensis* | Sheraliev, BM | Sheraliev, BM | Surxondaryo | Amu Darya | Sherobod river | 37.747222 | 66.996666 | 25/07/2019 |
| 87. | Capoeta_heratensis_432 | SWU27072019432 | FFU087-20 | MW649239 | Cyprinidae | *Capoeta heratensis* | Sheraliev, BM | Sheraliev, BM | Bukhara | Zeravshan | Echkiliksoy | 40.267551 | 64.743333 | 27/07/2019 |
| 88. | Capoeta_heratensis_433 | SWU27072019433 | FFU088-20 | MW649240 | Cyprinidae | *Capoeta heratensis* | Sheraliev, BM | Sheraliev, BM | Bukhara | Zeravshan | Echkiliksoy | 40.267551 | 64.743333 | 27/07/2019 |
| 89. | Capoeta_heratensis_434 | SWU27072019434 | FFU089-20 | MW649241 | Cyprinidae | *Capoeta heratensis* | Sheraliev, BM | Sheraliev, BM | Bukhara | Zeravshan | Echkiliksoy | 40.267551 | 64.743333 | 27/07/2019 |
| 90. | Capoeta_heratensis_474 | SWU29072019474 | FFU090-20 | MW649242 | Cyprinidae | *Capoeta heratensis* | Sheraliev, BM | Sheraliev, BM | Bukhara | Zeravshan | Unnamed stream | 40.147222 | 64.905805 | 29/07/2019 |
| 91. | Capoeta_heratensis_693 | SWU27042019693 | FFU091-20 | MW649243 | Cyprinidae | *Capoeta heratensis* | Allayarov, S | Sheraliev, BM | Surxondaryo | Amu Darya | Karatag river | 38.363055 | 68.072059 | 27/04/2019 |
| 92. | Capoeta_heratensis_694 | SWU27042019694 | FFU092-20 | MW649244 | Cyprinidae | *Capoeta heratensis* | Allayarov, S | Sheraliev, BM | Surxondaryo | Amu Darya | Karatag river | 38.363055 | 68.072059 | 27/04/2019 |
| 93. | Capoeta_heratensis_695 | SWU27042019695 | FFU093-20 | MW649245 | Cyprinidae | *Capoeta heratensis* | Allayarov, S | Sheraliev, BM | Surxondaryo | Amu Darya | Karatag river | 38.363055 | 68.072059 | 27/04/2019 |
| 94. | Carassius_auratus_134 | SWU22022018134 | FFU094-20 | MW649246 | Cyprinidae | *Carassius auratus* | Sheraliev, BM | Sheraliev, BM | Bukhara | Zeravshan | Qoraqir Lake | 40.385644 | 63.308388 | 22/02/2018 |
| 95. | Carassius_auratus_135 | SWU22022018135 | FFU095-20 | MW649247 | Cyprinidae | *Carassius auratus* | Sheraliev, BM | Sheraliev, BM | Bukhara | Zeravshan | Qoraqir Lake | 40.385644 | 63.308388 | 22/02/2018 |
| 96. | Carassius_auratus_136 | SWU22022018136 | FFU096-20 | MW649248 | Cyprinidae | *Carassius auratus* | Sheraliev, BM | Sheraliev, BM | Bukhara | Zeravshan | Qoraqir Lake | 40.385644 | 63.308388 | 22/02/2018 |
| 97. | Carassius_auratus_160 | SWU16072018160 | FFU097-20 | MW649249 | Cyprinidae | *Carassius auratus* | Sheraliev, BM | Sheraliev, BM | Bukhara | Zeravshan | Zeravshan | 40.146944 | 64.891666 | 16/07/2018 |
| 98. | Carassius_auratus_161 | SWU16072018161 | FFU098-20 | MW649250 | Cyprinidae | *Carassius auratus* | Sheraliev, BM | Sheraliev, BM | Bukhara | Zeravshan | Zeravshan | 40.146944 | 64.891666 | 16/07/2018 |
| 99. | Carassius_auratus_162 | SWU16072018162 | FFU099-20 | MW649251 | Cyprinidae | *Carassius auratus* | Sheraliev, BM | Sheraliev, BM | Bukhara | Zeravshan | Zeravshan | 40.146944 | 64.891666 | 16/07/2018 |
| 100. | Carassius_auratus_164 | SWU16072018164 | FFU100-20 | MW649252 | Cyprinidae | *Carassius auratus* | Sheraliev, BM | Sheraliev, BM | Bukhara | Zeravshan | Zeravshan | 40.146944 | 64.891666 | 16/07/2018 |
| 101. | Carassius_gibelio_011 | SWU14082016011 | FFU101-20 | MW649253 | Cyprinidae | *Carassius gibelio* | Sheraliev, BM | Sheraliev, BM | Surxondaryo | Amu Darya | South Surkhan R. | 37.854013 | 67.628373 | 14/08/2016 |
| 102. | Carassius_gibelio_012 | SWU14082016012 | FFU102-20 | MW649254 | Cyprinidae | *Carassius gibelio* | Sheraliev, BM | Sheraliev, BM | Surxondaryo | Amu Darya | South Surkhan R. | 37.854013 | 67.628373 | 14/08/2016 |
| 103. | Carassius_gibelio_024 | SWU16082016024 | FFU103-20 | MW649255 | Cyprinidae | *Carassius gibelio* | Sheraliev, BM | Sheraliev, BM | Qashqadaryo | Amu Darya | Qorasuv Stream | 38.570458 | 66.296166 | 16/08/2016 |
| 104. | Carassius_gibelio_025 | SWU16082016025 | FFU104-20 | MW649256 | Cyprinidae | *Carassius gibelio* | Sheraliev, BM | Sheraliev, BM | Qashqadaryo | Amu Darya | Qorasuv Stream | 38.570458 | 66.296166 | 16/08/2016 |
| 105. | Carassius_gibelio_026 | SWU16082016026 | FFU105-20 | MW649257 | Cyprinidae | *Carassius gibelio* | Sheraliev, BM | Sheraliev, BM | Qashqadaryo | Amu Darya | Qorasuv Stream | 38.570458 | 66.296166 | 16/08/2016 |
| 106. | Carassius_gibelio_033 | SWU31082016033 | FFU106-20 | MW649258 | Cyprinidae | *Carassius gibelio* | Sheraliev, BM | Sheraliev, BM | Namangan | Syr Darya | Syr Darya | 40.882777 | 71.442222 | 31/08/2016 |
| 107. | Carassius_gibelio_034 | SWU31082016034 | FFU107-20 | MW649259 | Cyprinidae | *Carassius gibelio* | Sheraliev, BM | Sheraliev, BM | Namangan | Syr Darya | Syr Darya | 40.882777 | 71.442222 | 31/08/2016 |
| 108. | Carassius_gibelio_035 | SWU31082016035 | FFU108-20 | MW649260 | Cyprinidae | *Carassius gibelio* | Sheraliev, BM | Sheraliev, BM | Namangan | Syr Darya | Syr Darya | 40.882777 | 71.442222 | 31/08/2016 |
| 109. | Carassius_gibelio_057 | SWU07012017057 | FFU109-20 | MW649261 | Cyprinidae | *Carassius gibelio* | Sheraliev, BM | Sheraliev, BM | Fergana | Syr Darya | Unnamed stream | 40.366437 | 71.316907 | 07/01/2017 |
| 110. | Carassius_gibelio_080 | SWU10072017080 | FFU110-20 | MW649262 | Cyprinidae | *Carassius gibelio* | Sheraliev, BM | Sheraliev, BM | Fergana | Syr Darya | Unnamed stream | 40.588943 | 71.537333 | 10/07/2017 |
| 111. | Carassius_gibelio_081 | SWU10072017081 | FFU111-20 | MW649263 | Cyprinidae | *Carassius gibelio* | Sheraliev, BM | Sheraliev, BM | Fergana | Syr Darya | Unnamed stream | 40.588943 | 71.537333 | 10/07/2017 |
| 112. | Carassius_gibelio_082 | SWU10072017082 | FFU112-20 | MW649264 | Cyprinidae | *Carassius gibelio* | Sheraliev, BM | Sheraliev, BM | Fergana | Syr Darya | Unnamed stream | 40.588943 | 71.537333 | 10/07/2017 |
| 113. | Carassius_gibelio_117 | SWU01022018117 | FFU113-20 | MW649265 | Cyprinidae | *Carassius gibelio* | Sheraliev, BM | Sheraliev, BM | Fergana | Syr Darya | Unnamed Stream | 40.484532 | 71.434862 | 01/02/2018 |
| 114. | Carassius_gibelio_118 | SWU01022018118 | FFU114-20 | MW649266 | Cyprinidae | *Carassius gibelio* | Sheraliev, BM | Sheraliev, BM | Fergana | Syr Darya | Unnamed Stream | 40.484532 | 71.434862 | 01/02/2018 |
| 115. | Carassius_gibelio_119 | SWU01022018119 | FFU115-20 | MW649267 | Cyprinidae | *Carassius gibelio* | Sheraliev, BM | Sheraliev, BM | Fergana | Syr Darya | Unnamed Stream | 40.484532 | 71.434862 | 01/02/2018 |
| 116. | Carassius_gibelio_137 | SWU22022018137 | FFU116-20 | MW649268 | Cyprinidae | *Carassius gibelio* | Sheraliev, BM | Sheraliev, BM | Bukhara | Zeravshan | Qoraqir Lake | 40.385644 | 63.308388 | 22/02/2018 |
| 117. | Carassius_gibelio_138 | SWU22022018138 | FFU117-20 | MW649269 | Cyprinidae | *Carassius gibelio* | Sheraliev, BM | Sheraliev, BM | Bukhara | Zeravshan | Qoraqir Lake | 40.385644 | 63.308388 | 22/02/2018 |
| 118. | Carassius_gibelio_139 | SWU22022018139 | FFU118-20 | MW649270 | Cyprinidae | *Carassius gibelio* | Sheraliev, BM | Sheraliev, BM | Bukhara | Zeravshan | Qoraqir Lake | 40.385644 | 63.308388 | 22/02/2018 |
| 119. | Carassius_gibelio_163 | SWU16072018163 | FFU119-20 | MW649271 | Cyprinidae | *Carassius gibelio* | Sheraliev, BM | Sheraliev, BM | Bukhara | Zeravshan | Zeravshan | 40.146944 | 64.891666 | 16/07/2018 |
| 120. | Carassius_gibelio_184 | SWU18072018184 | FFU120-20 | MW649272 | Cyprinidae | *Carassius gibelio* | Sheraliev, BM | Sheraliev, BM | Navoiy | Zeravshan | Tudakul Lake | 39.801388 | 64.772222 | 18/07/2018 |
| 121. | Carassius_gibelio_223 | SWU07012017223 | FFU121-20 | MW649273 | Cyprinidae | *Carassius gibelio* | Sheraliev, BM | Sheraliev, BM | Fergana | Syr Darya | Unnamed stream | 40.366437 | 71.316907 | 07/01/2017 |
| 122. | Carassius_gibelio_224 | SWU07012017224 | FFU122-20 | MW649274 | Cyprinidae | *Carassius gibelio* | Sheraliev, BM | Sheraliev, BM | Fergana | Syr Darya | Unnamed stream | 40.366437 | 71.316907 | 07/01/2017 |
| 123. | Carassius_gibelio_225 | SWU07012017225 | FFU123-20 | MW649275 | Cyprinidae | *Carassius gibelio* | Sheraliev, BM | Sheraliev, BM | Fergana | Syr Darya | Unnamed stream | 40.366437 | 71.316907 | 07/01/2017 |
| 124. | Carassius_gibelio_583 | SWU09082019583 | FFU124-20 | MW649276 | Cyprinidae | *Carassius gibelio* | Sheraliev, BM | Sheraliev, BM | Namangan | Syr Darya | Naryn River | 40.940911 | 71.853631 | 09/08/2019 |
| 125. | Carassius_gibelio_679 | SWU27042019679 | FFU125-20 | MW649277 | Cyprinidae | *Carassius gibelio* | Allayarov, S | Sheraliev, BM | Surxondaryo | Amu Darya | Karatag river | 38.363055 | 68.072059 | 27/04/2019 |
| 126. | Carassius_gibelio_680 | SWU27042019680 | FFU126-20 | MW649278 | Cyprinidae | *Carassius gibelio* | Allayarov, S | Sheraliev, BM | Surxondaryo | Amu Darya | Karatag river | 38.363055 | 68.072059 | 27/04/2019 |
| 127. | Carassius_gibelio_681 | SWU27042019681 | FFU127-20 | MW649279 | Cyprinidae | *Carassius gibelio* | Allayarov, S | Sheraliev, BM | Surxondaryo | Amu Darya | Karatag river | 38.363055 | 68.072059 | 27/04/2019 |
| 128. | Carassius_gibelio_682 | SWU27042019681 | FFU128-20 | MW649280 | Cyprinidae | *Carassius gibelio* | Allayarov, S | Sheraliev, BM | Surxondaryo | Amu Darya | Karatag river | 38.363055 | 68.072059 | 27/04/2019 |
| 129. | Cyprinus_carpio_013 | SWU14082016013 | FFU129-20 | MW649281 | Cyprinidae | *Cyprinus carpio* | Sheraliev, BM | Sheraliev, BM | Surxondaryo | Amu Darya | South Surkhan R. | 37.854013 | 67.628373 | 14/08/2016 |
| 130. | Cyprinus_carpio_021 | SWU16082016021 | FFU130-20 | MW649282 | Cyprinidae | *Cyprinus carpio* | Sheraliev, BM | Sheraliev, BM | Qashqadaryo | Amu Darya | Qorasuv Stream | 38.570458 | 66.296166 | 16/08/2016 |
| 131. | Cyprinus_carpio_022 | SWU16082016022 | FFU131-20 | MW649283 | Cyprinidae | *Cyprinus carpio* | Sheraliev, BM | Sheraliev, BM | Qashqadaryo | Amu Darya | Qorasuv Stream | 38.570458 | 66.296166 | 16/08/2016 |
| 132. | Cyprinus_carpio_023 | SWU16082016023 | FFU132-20 | MW649284 | Cyprinidae | *Cyprinus carpio* | Sheraliev, BM | Sheraliev, BM | Qashqadaryo | Amu Darya | Qorasuv Stream | 38.570458 | 66.296166 | 16/08/2016 |
| 133. | Cyprinus_carpio_077 | SWU10072017077 | FFU133-20 | MW649285 | Cyprinidae | *Cyprinus carpio* | Sheraliev, BM | Sheraliev, BM | Fergana | Syr Darya | Unnamed stream | 40.588943 | 71.537333 | 10/07/2017 |
| 134. | Cyprinus_carpio_078 | SWU10072017078 | FFU134-20 | MW649286 | Cyprinidae | *Cyprinus carpio* | Sheraliev, BM | Sheraliev, BM | Fergana | Syr Darya | Unnamed stream | 40.588943 | 71.537333 | 10/07/2017 |
| 135. | Cyprinus_carpio_079 | SWU10072017079 | FFU135-20 | MW649287 | Cyprinidae | *Cyprinus carpio* | Sheraliev, BM | Sheraliev, BM | Fergana | Syr Darya | Unnamed stream | 40.588943 | 71.537333 | 10/07/2017 |
| 136. | Cyprinus_carpio_154 | SWU16072018154 | FFU136-20 | MW649288 | Cyprinidae | *Cyprinus carpio* | Sheraliev, BM | Sheraliev, BM | Bukhara | Zeravshan | Zeravshan | 40.146944 | 64.891666 | 16/07/2018 |
| 137. | Cyprinus_carpio_183 | SWU18072018183 | FFU137-20 | MW649289 | Cyprinidae | *Cyprinus carpio* | Sheraliev, BM | Sheraliev, BM | Navoiy | Zeravshan | Tudakul Lake | 39.801388 | 64.772222 | 18/07/2018 |
| 138. | Luciobarbus_brachycephalus_483 | SWU31072019483 | FFU138-20 | MW649290 | Cyprinidae | *Luciobarbus brachycephalus* | Sheraliev, BM | Sheraliev, BM | Khorazm | Amu Darya | Amu Darya | 41.650833 | 60.704722 | 31/07/2019 |
| 139. | Luciobarbus_brachycephalus_484 | SWU31072019484 | FFU139-20 | MW649291 | Cyprinidae | *Luciobarbus brachycephalus* | Sheraliev, BM | Sheraliev, BM | Khorazm | Amu Darya | Amu Darya | 41.650833 | 60.704722 | 31/07/2019 |
| 140. | Luciobarbus_brachycephalus_485 | SWU31072019485 | FFU140-20 | MW649292 | Cyprinidae | *Luciobarbus brachycephalus* | Sheraliev, BM | Sheraliev, BM | Khorazm | Amu Darya | Amu Darya | 41.650833 | 60.704722 | 31/07/2019 |
| 141. | Luciobarbus_brachycephalus_486 | SWU31072019486 | FFU141-20 | MW649293 | Cyprinidae | *Luciobarbus brachycephalus* | Sheraliev, BM | Sheraliev, BM | Khorazm | Amu Darya | Amu Darya | 41.650833 | 60.704722 | 31/07/2019 |

| | | | | | | | | | | | | | | |
|---|---|---|---|---|---|---|---|---|---|---|---|---|---|---|
| 142. | Luciobarbus_brachycephalus_487 | SWU31072019487 | FFU142-20 | MW649294 | Cyprinidae | *Luciobarbus brachycephalus* | Sheraliev, BM | Sheraliev, BM | Khorazm | Amu Darya | Amu Darya | 41.650833 | 60.704722 | 31/07/2019 |
| 143. | Luciobarbus_brachycephalus_488 | SWU31072019488 | FFU143-20 | MW649295 | Cyprinidae | *Luciobarbus brachycephalus* | Sheraliev, BM | Sheraliev, BM | Khorazm | Amu Darya | Amu Darya | 41.650833 | 60.704722 | 31/07/2019 |
| 144. | Luciobarbus_brachycephalus_670 | SWU21082019670 | FFU144-20 | MW649296 | Cyprinidae | *Luciobarbus brachycephalus* | Rozimov, A | Sheraliev, BM | Khorazm | Amu Darya | Amu Darya | 41.650833 | 60.704722 | 21/08/2019 |
| 145. | Luciobarbus_brachycephalus_671 | SWU21082019671 | FFU145-20 | MW649297 | Cyprinidae | *Luciobarbus brachycephalus* | Rozimov, A | Sheraliev, BM | Khorazm | Amu Darya | Amu Darya | 41.650833 | 60.704722 | 21/08/2019 |
| 146. | Luciobarbus_conocephalus_041 | SWU31082016041 | FFU146-20 | MW649298 | Cyprinidae | *Luciobarbus conocephalus* | Sheraliev, BM | Sheraliev, BM | Namangan | Syr Darya | Syr Darya | 40.882777 | 71.442222 | 31/08/2016 |
| 147. | Luciobarbus_conocephalus_042 | SWU31082016042 | FFU147-20 | MW649299 | Cyprinidae | *Luciobarbus conocephalus* | Sheraliev, BM | Sheraliev, BM | Namangan | Syr Darya | Syr Darya | 40.882777 | 71.442222 | 31/08/2016 |
| 148. | Luciobarbus_conocephalus_044 | SWU31082016044 | FFU148-21 | MW649300 | Cyprinidae | *Luciobarbus conocephalus* | Sheraliev, BM | Sheraliev, BM | Namangan | Syr Darya | Syr Darya | 40.882777 | 71.442222 | 31/08/2016 |
| 149. | Luciobarbus_conocephalus_045 | SWU31082016045 | FFU149-21 | MW649301 | Cyprinidae | *Luciobarbus conocephalus* | Sheraliev, BM | Sheraliev, BM | Namangan | Syr Darya | Syr Darya | 40.882777 | 71.442222 | 31/08/2016 |
| 150. | Luciobarbus_conocephalus_046 | SWU31082016046 | FFU150-21 | MW649302 | Cyprinidae | *Luciobarbus conocephalus* | Sheraliev, BM | Sheraliev, BM | Namangan | Syr Darya | Syr Darya | 40.882777 | 71.442222 | 31/08/2016 |
| 151. | Luciobarbus_conocephalus_125 | SWU08022018125 | FFU151-21 | MW649303 | Cyprinidae | *Luciobarbus conocephalus* | Sheraliev, BM | Sheraliev, BM | Fergana | Syr Darya | Sokh River | 40.340176 | 71.013559 | 08/02/2018 |
| 152. | Luciobarbus_conocephalus_171 | SWU17072018171 | FFU152-21 | MW649304 | Cyprinidae | *Luciobarbus conocephalus* | Sheraliev, BM | Sheraliev, BM | Bukhara | Zeravshan | Dzhalvan canal | 40.149666 | 64.885400 | 17/07/2018 |
| 153. | Luciobarbus_conocephalus_172 | SWU17072018172 | FFU153-21 | MW649305 | Cyprinidae | *Luciobarbus conocephalus* | Sheraliev, BM | Sheraliev, BM | Bukhara | Zeravshan | Dzhalvan canal | 40.149666 | 64.885400 | 17/07/2018 |
| 154. | Luciobarbus_conocephalus_229 | SWU31082016229 | FFU154-21 | MW649306 | Cyprinidae | *Luciobarbus conocephalus* | Sheraliev, BM | Sheraliev, BM | Namangan | Syr Darya | Syr Darya | 40.882777 | 71.442222 | 31/08/2016 |
| 155. | Luciobarbus_conocephalus_310 | SWU21072019310 | FFU155-21 | MW649307 | Cyprinidae | *Luciobarbus conocephalus* | Sheraliev, BM | Sheraliev, BM | Surxondaryo | Amu Darya | Surkhan Darya | 37.238888 | 67.329722 | 21/07/2019 |
| 156. | Luciobarbus_conocephalus_311 | SWU21072019311 | FFU156-21 | MW649308 | Cyprinidae | *Luciobarbus conocephalus* | Sheraliev, BM | Sheraliev, BM | Surxondaryo | Amu Darya | Surkhan Darya | 37.238888 | 67.329722 | 21/07/2019 |
| 157. | Luciobarbus_conocephalus_312 | SWU21072019312 | FFU157-21 | MW649309 | Cyprinidae | *Luciobarbus conocephalus* | Sheraliev, BM | Sheraliev, BM | Surxondaryo | Amu Darya | Surkhan Darya | 37.238888 | 67.329722 | 21/07/2019 |
| 158. | Luciobarbus_conocephalus_331 | SWU22072019331 | FFU158-21 | MW649310 | Cyprinidae | *Luciobarbus conocephalus* | Sheraliev, BM | Sheraliev, BM | Surxondaryo | Amu Darya | Karatag river | 38.363055 | 68.072059 | 22/07/2019 |
| 159. | Luciobarbus_conocephalus_332 | SWU22072019332 | FFU159-21 | MW649311 | Cyprinidae | *Luciobarbus conocephalus* | Sheraliev, BM | Sheraliev, BM | Surxondaryo | Amu Darya | Karatag river | 38.363055 | 68.072059 | 22/07/2019 |
| 160. | Luciobarbus_conocephalus_333 | SWU22072019333 | FFU160-21 | MW649312 | Cyprinidae | *Luciobarbus conocephalus* | Sheraliev, BM | Sheraliev, BM | Surxondaryo | Amu Darya | Karatag river | 38.363055 | 68.072059 | 22/07/2019 |
| 161. | Schizothorax_eurystomus_043 | SWU31082016043 | FFU161-21 | MW649313 | Cyprinidae | *Schizothorax eurystomus* | Sheraliev, BM | Sheraliev, BM | Namangan | Syr Darya | Syr Darya | 40.882777 | 71.442222 | 31/08/2016 |
| 162. | Schizothorax_eurystomus_047 | SWU31082016047 | FFU162-21 | MW649314 | Cyprinidae | *Schizothorax eurystomus* | Sheraliev, BM | Sheraliev, BM | Namangan | Syr Darya | Syr Darya | 40.882777 | 71.442222 | 31/08/2016 |
| 163. | Schizothorax_eurystomus_048 | SWU31082016048 | FFU163-21 | MW649315 | Cyprinidae | *Schizothorax eurystomus* | Sheraliev, BM | Sheraliev, BM | Namangan | Syr Darya | Syr Darya | 40.882777 | 71.442222 | 31/08/2016 |
| 164. | Schizothorax_eurystomus_049 | SWU31082016049 | FFU164-21 | MW649316 | Cyprinidae | *Schizothorax eurystomus* | Sheraliev, BM | Sheraliev, BM | Namangan | Syr Darya | Syr Darya | 40.882777 | 71.442222 | 31/08/2016 |
| 165. | Schizothorax_eurystomus_058 | SWU07012017058 | FFU165-21 | MW649317 | Cyprinidae | *Schizothorax eurystomus* | Sheraliev, BM | Sheraliev, BM | Fergana | Syr Darya | Unnamed stream | 40.366437 | 71.316907 | 07/01/2017 |
| 166. | Schizothorax_eurystomus_120 | SWU08022018120 | FFU166-21 | MW649318 | Cyprinidae | *Schizothorax eurystomus* | Sheraliev, BM | Sheraliev, BM | Fergana | Syr Darya | Sokh River | 40.340176 | 71.013559 | 08/02/2018 |
| 167. | Schizothorax_eurystomus_121 | SWU08022018121 | FFU167-21 | MW649319 | Cyprinidae | *Schizothorax eurystomus* | Sheraliev, BM | Sheraliev, BM | Fergana | Syr Darya | Sokh River | 40.340176 | 71.013559 | 08/02/2018 |
| 168. | Schizothorax_eurystomus_122 | SWU08022018122 | FFU168-21 | MW649320 | Cyprinidae | *Schizothorax eurystomus* | Sheraliev, BM | Sheraliev, BM | Fergana | Syr Darya | Sokh River | 40.340176 | 71.013559 | 08/02/2018 |
| 169. | Schizothorax_eurystomus_123 | SWU08022018123 | FFU169-21 | MW649321 | Cyprinidae | *Schizothorax eurystomus* | Sheraliev, BM | Sheraliev, BM | Fergana | Syr Darya | Sokh River | 40.340176 | 71.013559 | 08/02/2018 |
| 170. | Schizothorax_eurystomus_124 | SWU08022018124 | FFU170-21 | MW649322 | Cyprinidae | *Schizothorax eurystomus* | Sheraliev, BM | Sheraliev, BM | Fergana | Syr Darya | Sokh River | 40.340176 | 71.013559 | 08/02/2018 |
| 171. | Schizothorax_eurystomus_198 | SWU11082018198 | FFU171-21 | MW649323 | Cyprinidae | *Schizothorax eurystomus* | Sheraliev, BM | Sheraliev, BM | Fergana | Syr Darya | Unnamed stream | 40.312769 | 71.811647 | 11/08/2018 |
| 172. | Schizothorax_eurystomus_199 | SWU11082018199 | FFU172-21 | MW649324 | Cyprinidae | *Schizothorax eurystomus* | Sheraliev, BM | Sheraliev, BM | Fergana | Syr Darya | Unnamed stream | 40.312769 | 71.811647 | 11/08/2018 |
| 173. | Schizothorax_eurystomus_200 | SWU11082018200 | FFU173-21 | MW649325 | Cyprinidae | *Schizothorax eurystomus* | Sheraliev, BM | Sheraliev, BM | Fergana | Syr Darya | Unnamed stream | 40.312769 | 71.811647 | 11/08/2018 |
| 174. | Schizothorax_eurystomus_201 | SWU11082018201 | FFU174-21 | MW649326 | Cyprinidae | *Schizothorax eurystomus* | Sheraliev, BM | Sheraliev, BM | Fergana | Syr Darya | Unnamed stream | 40.312769 | 71.811647 | 11/08/2018 |
| 175. | Schizothorax_eurystomus_202 | SWU11082018202 | FFU175-21 | MW649327 | Cyprinidae | *Schizothorax eurystomus* | Sheraliev, BM | Sheraliev, BM | Fergana | Syr Darya | Unnamed stream | 40.312769 | 71.811647 | 11/08/2018 |
| 176. | Schizothorax_eurystomus_203 | SWU11082018203 | FFU176-21 | MW649328 | Cyprinidae | *Schizothorax eurystomus* | Sheraliev, BM | Sheraliev, BM | Fergana | Syr Darya | Unnamed stream | 40.312769 | 71.811647 | 11/08/2018 |
| 177. | Schizothorax_eurystomus_211 | SWU15082018211 | FFU177-21 | MW649329 | Cyprinidae | *Schizothorax eurystomus* | Sheraliev, BM | Sheraliev, BM | Fergana | Syr Darya | Shohimardon River | 39.963088 | 71.759857 | 15/08/2018 |
| 178. | Schizothorax_eurystomus_212 | SWU15082018212 | FFU178-21 | MW649330 | Cyprinidae | *Schizothorax eurystomus* | Sheraliev, BM | Sheraliev, BM | Fergana | Syr Darya | Shohimardon River | 39.963088 | 71.759857 | 15/08/2018 |
| 179. | Schizothorax_eurystomus_213 | SWU15082018213 | FFU179-21 | MW649331 | Cyprinidae | *Schizothorax eurystomus* | Sheraliev, BM | Sheraliev, BM | Fergana | Syr Darya | Shohimardon River | 39.963088 | 71.759857 | 15/08/2018 |
| 180. | Schizothorax_eurystomus_214 | SWU15082018214 | FFU180-21 | MW649332 | Cyprinidae | *Schizothorax eurystomus* | Sheraliev, BM | Sheraliev, BM | Fergana | Syr Darya | Shohimardon River | 39.963088 | 71.759857 | 15/08/2018 |
| 181. | Schizothorax_eurystomus_215 | SWU15082018215 | FFU181-21 | MW649333 | Cyprinidae | *Schizothorax eurystomus* | Sheraliev, BM | Sheraliev, BM | Fergana | Syr Darya | Shohimardon River | 39.963088 | 71.759857 | 15/08/2018 |
| 182. | Schizothorax_eurystomus_226 | SWU15012017226 | FFU182-21 | MW649334 | Cyprinidae | *Schizothorax eurystomus* | Sheraliev, BM | Sheraliev, BM | Fergana | Syr Darya | Unnamed stream | 40.326408 | 71.820508 | 15/01/2017 |
| 183. | Schizothorax_eurystomus_227 | SWU15012017227 | FFU183-21 | MW649335 | Cyprinidae | *Schizothorax eurystomus* | Sheraliev, BM | Sheraliev, BM | Fergana | Syr Darya | Unnamed stream | 40.326408 | 71.820508 | 15/01/2017 |
| 184. | Schizothorax_eurystomus_228 | SWU15012017228 | FFU184-21 | MW649336 | Cyprinidae | *Schizothorax eurystomus* | Sheraliev, BM | Sheraliev, BM | Fergana | Syr Darya | Unnamed stream | 40.326408 | 71.820508 | 15/01/2017 |
| 185. | Schizothorax_eurystomus_542 | SWU05082019542 | FFU185-21 | MW649337 | Cyprinidae | *Schizothorax eurystomus* | Bakhromova, B | Sheraliev, BM | Fergana | Syr Darya | Sokh River | 39.980157 | 71.118103 | 05/08/2019 |
| 186. | Schizothorax_eurystomus_563 | SWU09082019563 | FFU186-21 | MW649338 | Cyprinidae | *Schizothorax eurystomus* | Sheraliev, BM | Sheraliev, BM | Namangan | Syr Darya | Kara Darya | 40.918813 | 71.828896 | 09/08/2019 |
| 187. | Schizothorax_eurystomus_564 | SWU09082019564 | FFU187-21 | MW649339 | Cyprinidae | *Schizothorax eurystomus* | Sheraliev, BM | Sheraliev, BM | Namangan | Syr Darya | Kara Darya | 40.918813 | 71.828896 | 09/08/2019 |
| 188. | Schizothorax_eurystomus_565 | SWU09082019565 | FFU188-21 | MW649340 | Cyprinidae | *Schizothorax eurystomus* | Sheraliev, BM | Sheraliev, BM | Namangan | Syr Darya | Kara Darya | 40.918813 | 71.828896 | 09/08/2019 |
| 189. | Schizothorax_eurystomus_566 | SWU09082019566 | FFU189-21 | MW649341 | Cyprinidae | *Schizothorax eurystomus* | Sheraliev, BM | Sheraliev, BM | Namangan | Syr Darya | Kara Darya | 40.918813 | 71.828896 | 09/08/2019 |
| 190. | Schizothorax_eurystomus_567 | SWU09082019567 | FFU190-21 | MW649342 | Cyprinidae | *Schizothorax eurystomus* | Sheraliev, BM | Sheraliev, BM | Namangan | Syr Darya | Kara Darya | 40.918813 | 71.828896 | 09/08/2019 |
| 191. | Schizothorax_eurystomus_577 | SWU09082019577 | FFU191-21 | MW649343 | Cyprinidae | *Schizothorax eurystomus* | Sheraliev, BM | Sheraliev, BM | Namangan | Syr Darya | Kara Darya | 40.918813 | 71.828896 | 09/08/2019 |
| 192. | Schizothorax_eurystomus_578 | SWU09082019578 | FFU192-21 | MW649344 | Cyprinidae | *Schizothorax eurystomus* | Sheraliev, BM | Sheraliev, BM | Namangan | Syr Darya | Kara Darya | 40.918813 | 71.828896 | 09/08/2019 |
| 193. | Schizothorax_eurystomus_584 | SWU09082019584 | FFU193-21 | MW649345 | Cyprinidae | *Schizothorax eurystomus* | Sheraliev, BM | Sheraliev, BM | Namangan | Syr Darya | Naryn River | 40.940911 | 71.853631 | 09/08/2019 |
| 194. | Schizothorax_eurystomus_595 | SWU18082019595 | FFU194-21 | MW649346 | Cyprinidae | *Schizothorax eurystomus* | Sheraliev, BM | Sheraliev, BM | Tashkent | Syr Darya | Chirchik river | 41.274444 | 69.389166 | 18/08/2019 |
| 195. | Schizothorax_eurystomus_596 | SWU18082019596 | FFU195-21 | MW649347 | Cyprinidae | *Schizothorax eurystomus* | Sheraliev, BM | Sheraliev, BM | Tashkent | Syr Darya | Chirchik river | 41.274444 | 69.389166 | 18/08/2019 |
| 196. | Schizothorax_eurystomus_597 | SWU18082019597 | FFU196-21 | MW649348 | Cyprinidae | *Schizothorax eurystomus* | Sheraliev, BM | Sheraliev, BM | Tashkent | Syr Darya | Chirchik river | 41.274444 | 69.389166 | 18/08/2019 |
| 197. | Schizothorax_eurystomus_644 | SWU20082019644 | FFU197-21 | MW649349 | Cyprinidae | *Schizothorax eurystomus* | Sheraliev, BM | Sheraliev, BM | Fergana | Syr Darya | Unnamed stream | 40.305260 | 71.800688 | 20/08/2019 |
| 198. | Schizothorax_eurystomus_645 | SWU20082019645 | FFU198-21 | MW649350 | Cyprinidae | *Schizothorax eurystomus* | Sheraliev, BM | Sheraliev, BM | Fergana | Syr Darya | Unnamed stream | 40.305260 | 71.800688 | 20/08/2019 |
| 199. | Schizothorax_fedtschenkoi_504 | SWU02082019504 | FFU199-21 | MW649351 | Cyprinidae | *Schizothorax fedtschenkoi* | Sheraliev, BM | Sheraliev, BM | Samarkand | Zeravshan | Zeravshan | 39.676388 | 67.076111 | 02/08/2019 |
| 200. | Schizothorax_fedtschenkoi_505 | SWU02082019505 | FFU200-21 | MW649352 | Cyprinidae | *Schizothorax fedtschenkoi* | Sheraliev, BM | Sheraliev, BM | Samarkand | Zeravshan | Zeravshan | 39.676388 | 67.076111 | 02/08/2019 |
| 201. | Schizothorax_fedtschenkoi_506 | SWU02082019506 | FFU201-21 | MW649353 | Cyprinidae | *Schizothorax fedtschenkoi* | Sheraliev, BM | Sheraliev, BM | Samarkand | Zeravshan | Zeravshan | 39.676388 | 67.076111 | 02/08/2019 |
| 202. | Schizothorax_fedtschenkoi_511 | SWU02082019511 | FFU202-21 | MW649354 | Cyprinidae | *Schizothorax fedtschenkoi* | Sheraliev, BM | Sheraliev, BM | Samarkand | Zeravshan | Zeravshan | 39.676388 | 67.076111 | 02/08/2019 |
| 203. | Schizothorax_fedtschenkoi_512 | SWU02082019512 | FFU203-21 | MW649355 | Cyprinidae | *Schizothorax fedtschenkoi* | Sheraliev, BM | Sheraliev, BM | Samarkand | Zeravshan | Zeravshan | 39.676388 | 67.076111 | 02/08/2019 |
| 204. | Schizothorax_sp_006 | SWU08082016006 | FFU662-21 | MW649356 | Cyprinidae | *Schizothorax* sp. | Sheraliev, BM | Sheraliev, BM | Surxondaryo | Amu Darya | Tupalang river | 38.399802 | 67.944766 | 08/08/2016 |
| 205. | Schizothorax_sp_007 | SWU08082016007 | FFU663-21 | MW649357 | Cyprinidae | *Schizothorax* sp. | Sheraliev, BM | Sheraliev, BM | Surxondaryo | Amu Darya | Tupalang river | 38.399802 | 67.944766 | 08/08/2016 |
| 206. | Schizothorax_sp_235 | SWU12082016235 | FFU664-21 | MW649358 | Cyprinidae | *Schizothorax* sp. | Sheraliev, BM | Sheraliev, BM | Surxondaryo | Amu Darya | Tupalang river | 38.399802 | 67.944766 | 12/08/2016 |
| 207. | Schizothorax_sp_402 | SWU25072019402 | FFU665-21 | MW649359 | Cyprinidae | *Schizothorax* sp. | Sheraliev, BM | Sheraliev, BM | Surxondaryo | Amu Darya | Sherobod river | 37.747222 | 66.996666 | 25/07/2019 |
| 208. | Schizothorax_sp_415 | SWU24072019415 | FFU666-21 | MW649360 | Cyprinidae | *Schizothorax* sp. | Sheraliev, BM | Sheraliev, BM | Surxondaryo | Amu Darya | Sayrob spring | 38.073178 | 66.968351 | 24/07/2019 |
| 209. | Esox_lucius_701 | SWU09112019701 | FFU204-21 | MW649361 | Esocidae | *Esox lucius* | Rozimov, A | Sheraliev, BM | Khorazm | Amu Darya | Unnamed stream | 41.488388 | 60.828055 | 09/11/2019 |
| 210. | Esox_lucius_702 | SWU13112010702 | FFU205-21 | MW649362 | Esocidae | *Esox lucius* | Rozimov, A | Sheraliev, BM | Khorazm | Amu Darya | Unnamed stream | 41.488388 | 60.828055 | 13/11/2019 |
| 211. | Esox_lucius_703 | SWU13112019703 | FFU206-21 | MW649363 | Esocidae | *Esox lucius* | Rozimov, A | Sheraliev, BM | Khorazm | Amu Darya | Unnamed stream | 41.488388 | 60.828055 | 13/11/2019 |
| 212. | Neogobius_melanostomus_716 | SWU18052020716 | FFU207-21 | MW649364 | Gobiidae | *Neogobius melanostonus* | Rozimov, A | Sheraliev, BM | Karakalpakstan | Amu Darya | Central canal | 42.635638 | 59.806944 | 18/05/2020 |
| 213. | Neogobius_pallasi_717 | SWU18052020717 | FFU208-21 | MW649365 | Gobiidae | *Neogobius pallasi* | Rozimov, A | Sheraliev, BM | Karakalpakstan | Amu Darya | Central canal | 42.635638 | 59.806944 | 18/05/2020 |
| 214. | Rhinogobius_sp_264 | SWU19072019264 | FFU209-21 | MW649366 | Gobiidae | *Rhinogobius* sp. | Sheraliev, BM | Sheraliev, BM | Surxondaryo | Amu Darya | Surkhan Darya | 37.255833 | 67.338611 | 19/07/2019 |
| 215. | Rhinogobius_sp_265 | SWU19072019265 | FFU210-21 | MW649367 | Gobiidae | *Rhinogobius* sp. | Sheraliev, BM | Sheraliev, BM | Surxondaryo | Amu Darya | Surkhan Darya | 37.255833 | 67.338611 | 19/07/2019 |
| 216. | Rhinogobius_sp_339 | SWU22072019339 | FFU211-21 | MW649368 | Gobiidae | *Rhinogobius* sp. | Sheraliev, BM | Sheraliev, BM | Surxondaryo | Amu Darya | Karatag river | 38.363055 | 68.072059 | 22/07/2019 |
| 217. | Rhinogobius_sp_340 | SWU22072019340 | FFU212-21 | MW649369 | Gobiidae | *Rhinogobius* sp. | Sheraliev, BM | Sheraliev, BM | Surxondaryo | Amu Darya | Karatag river | 38.363055 | 68.072059 | 22/07/2019 |
| 218. | Rhinogobius_sp_341 | SWU22072019341 | FFU213-21 | MW649370 | Gobiidae | *Rhinogobius* sp. | Sheraliev, BM | Sheraliev, BM | Surxondaryo | Amu Darya | Karatag river | 38.363055 | 68.072059 | 22/07/2019 |

| | | | | | | | | | | | | | | |
|---|---|---|---|---|---|---|---|---|---|---|---|---|---|---|
| 219. | Rhinogobius_sp_342 | SWU22072019342 | FFU214-21 | MW649371 | Gobiidae | *Rhinogobius* sp. | Sheraliev, BM | Sheraliev, BM | Surxondaryo | Amu Darya | Karatag river | 38.363055 | 68.072059 | 22/07/2019 |
| 220. | Rhinogobius_sp_343 | SWU22072019343 | FFU215-21 | MW649372 | Gobiidae | *Rhinogobius* sp. | Sheraliev, BM | Sheraliev, BM | Surxondaryo | Amu Darya | Karatag river | 38.363055 | 68.072059 | 22/07/2019 |
| 221. | Rhinogobius_sp_397 | SWU24072019397 | FFU216-21 | MW649373 | Gobiidae | *Rhinogobius* sp. | Sheraliev, BM | Sheraliev, BM | Surxondaryo | Amu Darya | Surkhan Darya | 37.823333 | 67.618611 | 24/07/2019 |
| 222. | Rhinogobius_sp_398 | SWU24072019398 | FFU217-21 | MW649374 | Gobiidae | *Rhinogobius* sp. | Sheraliev, BM | Sheraliev, BM | Surxondaryo | Amu Darya | Surkhan Darya | 37.823333 | 67.618611 | 24/07/2019 |
| 223. | Rhinogobius_sp_399 | SWU24072019399 | FFU218-21 | MW649375 | Gobiidae | *Rhinogobius* sp. | Sheraliev, BM | Sheraliev, BM | Surxondaryo | Amu Darya | Surkhan Darya | 37.823333 | 67.618611 | 24/07/2019 |
| 224. | Rhinogobius_sp_400 | SWU24072019400 | FFU219-21 | MW649376 | Gobiidae | *Rhinogobius* sp. | Sheraliev, BM | Sheraliev, BM | Surxondaryo | Amu Darya | Surkhan Darya | 37.823333 | 67.618611 | 24/07/2019 |
| 225. | Rhinogobius_sp_417 | SWU27072019417 | FFU220-21 | MW649377 | Gobiidae | *Rhinogobius* sp. | Sheraliev, BM | Sheraliev, BM | Bukhara | Zeravshan | Echkiliksoy | 40.267551 | 64.743333 | 27/07/2019 |
| 226. | Rhinogobius_sp_418 | SWU27072019418 | FFU221-21 | MW649378 | Gobiidae | *Rhinogobius* sp. | Sheraliev, BM | Sheraliev, BM | Bukhara | Zeravshan | Echkiliksoy | 40.267551 | 64.743333 | 27/07/2019 |
| 227. | Rhinogobius_sp_419 | SWU27072019419 | FFU222-21 | MW649379 | Gobiidae | *Rhinogobius* sp. | Sheraliev, BM | Sheraliev, BM | Bukhara | Zeravshan | Echkiliksoy | 40.267551 | 64.743333 | 27/07/2019 |
| 228. | Rhinogobius_sp_420 | SWU27072019420 | FFU223-21 | MW649380 | Gobiidae | *Rhinogobius* sp. | Sheraliev, BM | Sheraliev, BM | Bukhara | Zeravshan | Echkiliksoy | 40.267551 | 64.743333 | 27/07/2019 |
| 229. | Rhinogobius_sp_437 | SWU27072019437 | FFU224-21 | MW649381 | Gobiidae | *Rhinogobius* sp. | Sheraliev, BM | Sheraliev, BM | Bukhara | Zeravshan | Echkiliksoy | 40.267551 | 64.743333 | 27/07/2019 |
| 230. | Rhinogobius_sp_438 | SWU27072019438 | FFU225-21 | MW649382 | Gobiidae | *Rhinogobius* sp. | Sheraliev, BM | Sheraliev, BM | Bukhara | Zeravshan | Echkiliksoy | 40.267551 | 64.743333 | 27/07/2019 |
| 231. | Rhinogobius_sp_459 | SWU28072019459 | FFU226-21 | MW649383 | Gobiidae | *Rhinogobius* sp. | Sheraliev, BM | Sheraliev, BM | Bukhara | Zeravshan | Zeravshan | 40.071012 | 64.778429 | 28/07/2019 |
| 232. | Rhinogobius_sp_467 | SWU28072019467 | FFU227-21 | MW649384 | Gobiidae | *Rhinogobius* sp. | Sheraliev, BM | Sheraliev, BM | Bukhara | Zeravshan | Zeravshan | 40.071012 | 64.778429 | 28/07/2019 |
| 233. | Rhinogobius_sp_468 | SWU28072019468 | FFU228-21 | MW649385 | Gobiidae | *Rhinogobius* sp. | Sheraliev, BM | Sheraliev, BM | Bukhara | Zeravshan | Zeravshan | 40.071012 | 64.778429 | 28/07/2019 |
| 234. | Rhinogobius_sp_493 | SWU31072019493 | FFU229-21 | MW649386 | Gobiidae | *Rhinogobius* sp. | Sheraliev, BM | Sheraliev, BM | Khorazm | Amu Darya | Amu Darya | 41.650833 | 60.704722 | 31/07/2019 |
| 235. | Rhinogobius_sp_494 | SWU31072019494 | FFU230-21 | MW649387 | Gobiidae | *Rhinogobius* sp. | Sheraliev, BM | Sheraliev, BM | Khorazm | Amu Darya | Amu Darya | 41.650833 | 60.704722 | 31/07/2019 |
| 236. | Rhinogobius_sp_495 | SWU31072019495 | FFU231-21 | MW649388 | Gobiidae | *Rhinogobius* sp. | Sheraliev, BM | Sheraliev, BM | Khorazm | Amu Darya | Amu Darya | 41.650833 | 60.704722 | 31/07/2019 |
| 237. | Rhinogobius_sp_496 | SWU31072019496 | FFU232-21 | MW649389 | Gobiidae | *Rhinogobius* sp. | Sheraliev, BM | Sheraliev, BM | Khorazm | Amu Darya | Amu Darya | 41.650833 | 60.704722 | 31/07/2019 |
| 238. | Rhinogobius_sp_497 | SWU31072019497 | FFU233-21 | MW649390 | Gobiidae | *Rhinogobius* sp. | Sheraliev, BM | Sheraliev, BM | Khorazm | Amu Darya | Amu Darya | 41.650833 | 60.704722 | 31/07/2019 |
| 239. | Rhinogobius_sp_543 | SWU09082019543 | FFU234-21 | MW649391 | Gobiidae | *Rhinogobius* sp. | Sheraliev, BM | Sheraliev, BM | Namangan | Syr Darya | Kara Darya | 40.918813 | 71.828896 | 09/08/2019 |
| 240. | Rhinogobius_sp_544 | SWU09082019544 | FFU235-21 | MW649392 | Gobiidae | *Rhinogobius* sp. | Sheraliev, BM | Sheraliev, BM | Namangan | Syr Darya | Kara Darya | 40.918813 | 71.828896 | 09/08/2019 |
| 241. | Rhinogobius_sp_545 | SWU09082019545 | FFU236-21 | MW649393 | Gobiidae | *Rhinogobius* sp. | Sheraliev, BM | Sheraliev, BM | Namangan | Syr Darya | Kara Darya | 40.918813 | 71.828896 | 09/08/2019 |
| 242. | Rhinogobius_sp_546 | SWU09082019546 | FFU237-21 | MW649394 | Gobiidae | *Rhinogobius* sp. | Sheraliev, BM | Sheraliev, BM | Namangan | Syr Darya | Kara Darya | 40.918813 | 71.828896 | 09/08/2019 |
| 243. | Rhinogobius_sp_547 | SWU09082019547 | FFU238-21 | MW649395 | Gobiidae | *Rhinogobius* sp. | Sheraliev, BM | Sheraliev, BM | Namangan | Syr Darya | Kara Darya | 40.918813 | 71.828896 | 09/08/2019 |
| 244. | Rhinogobius_sp_549 | SWU09082019549 | FFU239-21 | MW649396 | Gobiidae | *Rhinogobius* sp. | Sheraliev, BM | Sheraliev, BM | Namangan | Syr Darya | Kara Darya | 40.918813 | 71.828896 | 09/08/2019 |
| 245. | Rhinogobius_sp_586 | SWU10082019586 | FFU240-21 | MW649397 | Gobiidae | *Rhinogobius* sp. | Sheraliev, BM | Sheraliev, BM | Namangan | Syr Darya | Syr Darya | 40.881008 | 71.686687 | 10/08/2019 |
| 246. | Rhinogobius_sp_587 | SWU10082019587 | FFU241-21 | MW649398 | Gobiidae | *Rhinogobius* sp. | Sheraliev, BM | Sheraliev, BM | Namangan | Syr Darya | Syr Darya | 40.881008 | 71.686687 | 10/08/2019 |
| 247. | Rhinogobius_sp_588 | SWU10082019588 | FFU242-21 | MW649399 | Gobiidae | *Rhinogobius* sp. | Sheraliev, BM | Sheraliev, BM | Namangan | Syr Darya | Syr Darya | 40.881008 | 71.686687 | 10/08/2019 |
| 248. | Rhinogobius_sp_589 | SWU10082019589 | FFU243-21 | MW649400 | Gobiidae | *Rhinogobius* sp. | Sheraliev, BM | Sheraliev, BM | Namangan | Syr Darya | Syr Darya | 40.881008 | 71.686687 | 10/08/2019 |
| 249. | Rhinogobius_sp_590 | SWU10082019590 | FFU244-21 | MW649401 | Gobiidae | *Rhinogobius* sp. | Sheraliev, BM | Sheraliev, BM | Namangan | Syr Darya | Syr Darya | 40.881008 | 71.686687 | 10/08/2019 |
| 250. | Rhinogobius_sp_613 | SWU18082019613 | FFU245-21 | MW649402 | Gobiidae | *Rhinogobius* sp. | Sheraliev, BM | Sheraliev, BM | Tashkent | Syr Darya | Chirchik River | 41.274444 | 69.389166 | 18/08/2019 |
| 251. | Abbottina_rivularis_059 | SWU07012017059 | FFU246-21 | MW649403 | Gobionidae | *Abbottina rivularis* | Sheraliev, BM | Sheraliev, BM | Fergana | Syr Darya | Unnamed stream | 40.366437 | 71.316907 | 07/01/2017 |
| 252. | Abbottina_rivularis_060 | SWU07012017060 | FFU247-21 | MW649404 | Gobionidae | *Abbottina rivularis* | Sheraliev, BM | Sheraliev, BM | Fergana | Syr Darya | Unnamed stream | 40.366437 | 71.316907 | 07/01/2017 |
| 253. | Abbottina_rivularis_153 | SWU16072018153 | FFU248-21 | MW649405 | Gobionidae | *Abbottina rivularis* | Sheraliev, BM | Sheraliev, BM | Bukhara | Zeravshan | Zeravshan | 40.146944 | 64.891666 | 16/07/2018 |
| 254. | Abbottina_rivularis_285 | SWU20072019285 | FFU249-21 | MW649406 | Gobionidae | *Abbottina rivularis* | Sheraliev, BM | Sheraliev, BM | Surxondaryo | Amu Darya | Amu Darya | 37.235555 | 67.660555 | 20/07/2019 |
| 255. | Abbottina_rivularis_338 | SWU22072019338 | FFU250-21 | MW649407 | Gobionidae | *Abbottina rivularis* | Sheraliev, BM | Sheraliev, BM | Surxondaryo | Amu Darya | Karatag river | 38.363055 | 68.072059 | 22/07/2019 |
| 256. | Abbottina_rivularis_428 | SWU27072019428 | FFU251-21 | MW649408 | Gobionidae | *Abbottina rivularis* | Sheraliev, BM | Sheraliev, BM | Bukhara | Zeravshan | Echkiliksoy | 40.267551 | 64.743333 | 27/07/2019 |
| 257. | Abbottina_rivularis_465 | SWU28072019465 | FFU252-21 | MW649409 | Gobionidae | *Abbottina rivularis* | Sheraliev, BM | Sheraliev, BM | Bukhara | Zeravshan | Zeravshan | 40.071012 | 64.778429 | 28/07/2019 |
| 258. | Abbottina_rivularis_498 | SWU31072019498 | FFU253-21 | MW649410 | Gobionidae | *Abbottina rivularis* | Sheraliev, BM | Sheraliev, BM | Khorazm | Amu Darya | Amu Darya | 41.650833 | 60.704722 | 31/07/2019 |
| 259. | Abbottina_rivularis_602 | SWU18082019602 | FFU254-21 | MW649411 | Gobionidae | *Abbottina rivularis* | Sheraliev, BM | Sheraliev, BM | Tashkent | Syr Darya | Chirchik river | 41.274444 | 69.389166 | 18/08/2019 |
| 260. | Abbottina_rivularis_690 | SWU27042019690 | FFU255-21 | MW649412 | Gobionidae | *Abbottina rivularis* | Allayarov, S | Sheraliev, BM | Surxondaryo | Amu Darya | Karatag river | 38.363055 | 68.072059 | 27/04/2019 |
| 261. | Abbottina_rivularis_691 | SWU27042019691 | FFU256-21 | MW649413 | Gobionidae | *Abbottina rivularis* | Allayarov, S | Sheraliev, BM | Surxondaryo | Amu Darya | Karatag river | 38.363055 | 68.072059 | 27/04/2019 |
| 262. | Gobio_lepidolaemus_031 | SWU31082016031 | FFU257-21 | MW649414 | Gobionidae | *Gobio lepidolaemus* | Sheraliev, BM | Sheraliev, BM | Namangan | Syr Darya | Syr Darya | 40.882777 | 71.442222 | 31/08/2016 |
| 263. | Gobio_lepidolaemus_061 | SWU07012017061 | FFU258-21 | MW649415 | Gobionidae | *Gobio lepidolaemus* | Sheraliev, BM | Sheraliev, BM | Fergana | Syr Darya | Unnamed stream | 40.366437 | 71.316907 | 07/01/2017 |
| 264. | Gobio_lepidolaemus_062 | SWU07012017062 | FFU259-21 | MW649416 | Gobionidae | *Gobio lepidolaemus* | Sheraliev, BM | Sheraliev, BM | Fergana | Syr Darya | Unnamed stream | 40.366437 | 71.316907 | 07/01/2017 |
| 265. | Gobio_lepidolaemus_063 | SWU07012017063 | FFU260-21 | MW649417 | Gobionidae | *Gobio lepidolaemus* | Sheraliev, BM | Sheraliev, BM | Fergana | Syr Darya | Unnamed stream | 40.366437 | 71.316907 | 07/01/2017 |
| 266. | Gobio_lepidolaemus_098 | SWU25012018098 | FFU261-21 | MW649418 | Gobionidae | *Gobio lepidolaemus* | Sheraliev, BM | Sheraliev, BM | Fergana | Syr Darya | Fergana Canal | 40.388497 | 71.327960 | 25/01/2018 |
| 267. | Gobio_lepidolaemus_099 | SWU25012018099 | FFU262-21 | MW649419 | Gobionidae | *Gobio lepidolaemus* | Sheraliev, BM | Sheraliev, BM | Fergana | Syr Darya | Fergana Canal | 40.388497 | 71.327960 | 25/01/2018 |
| 268. | Gobio_lepidolaemus_100 | SWU25012018100 | FFU263-21 | MW649420 | Gobionidae | *Gobio lepidolaemus* | Sheraliev, BM | Sheraliev, BM | Fergana | Syr Darya | Fergana Canal | 40.388497 | 71.327960 | 25/01/2018 |
| 269. | Gobio_lepidolaemus_192 | SWU11082018192 | FFU264-21 | MW649421 | Gobionidae | *Gobio lepidolaemus* | Sheraliev, BM | Sheraliev, BM | Fergana | Syr Darya | Unnamed stream | 40.312769 | 71.811647 | 11/08/2018 |
| 270. | Gobio_lepidolaemus_193 | SWU11082018193 | FFU265-21 | MW649422 | Gobionidae | *Gobio lepidolaemus* | Sheraliev, BM | Sheraliev, BM | Fergana | Syr Darya | Unnamed stream | 40.312769 | 71.811647 | 11/08/2018 |
| 271. | Gobio_lepidolaemus_194 | SWU11082018194 | FFU266-21 | MW649423 | Gobionidae | *Gobio lepidolaemus* | Sheraliev, BM | Sheraliev, BM | Fergana | Syr Darya | Unnamed stream | 40.312769 | 71.811647 | 11/08/2018 |
| 272. | Gobio_lepidolaemus_195 | SWU11082018195 | FFU267-21 | MW649424 | Gobionidae | *Gobio lepidolaemus* | Sheraliev, BM | Sheraliev, BM | Fergana | Syr Darya | Unnamed stream | 40.312769 | 71.811647 | 11/08/2018 |
| 273. | Gobio_lepidolaemus_196 | SWU11082018196 | FFU268-21 | MW649425 | Gobionidae | *Gobio lepidolaemus* | Sheraliev, BM | Sheraliev, BM | Fergana | Syr Darya | Unnamed stream | 40.312769 | 71.811647 | 11/08/2018 |
| 274. | Gobio_lepidolaemus_197 | SWU11082018197 | FFU269-21 | MW649426 | Gobionidae | *Gobio lepidolaemus* | Sheraliev, BM | Sheraliev, BM | Fergana | Syr Darya | Unnamed stream | 40.312769 | 71.811647 | 11/08/2018 |
| 275. | Gobio_lepidolaemus_257 | SWU19072019257 | FFU270-21 | MN810113 | Gobionidae | *Gobio lepidolaemus* | Sheraliev, BM | Sheraliev, BM | Surxondaryo | Amu Darya | Surkhan Darya | 37.255833 | 67.338611 | 19/07/2019 |
| 276. | Gobio_lepidolaemus_568 | SWU09082019568 | FFU271-21 | MN810114 | Gobionidae | *Gobio lepidolaemus* | Sheraliev, BM | Sheraliev, BM | Namangan | Syr Darya | Kara Darya | 40.918813 | 71.828896 | 09/08/2019 |
| 277. | Gobio_lepidolaemus_569 | SWU09082019569 | FFU272-21 | MW649427 | Gobionidae | *Gobio lepidolaemus* | Sheraliev, BM | Sheraliev, BM | Namangan | Syr Darya | Kara Darya | 40.918813 | 71.828896 | 09/08/2019 |
| 278. | Gobio_lepidolaemus_579 | SWU09082019579 | FFU273-21 | MW649428 | Gobionidae | *Gobio lepidolaemus* | Sheraliev, BM | Sheraliev, BM | Namangan | Syr Darya | Kara Darya | 40.918813 | 71.828896 | 09/08/2019 |
| 279. | Gobio_lepidolaemus_603 | SWU18082019603 | FFU274-21 | MW649429 | Gobionidae | *Gobio lepidolaemus* | Sheraliev, BM | Sheraliev, BM | Tashkent | Syr Darya | Chirchik river | 41.274444 | 69.389166 | 18/08/2019 |
| 280. | Gobio_nigrescens_241 | SWU16082016241 | FFU275-21 | MN810111 | Gobionidae | *Gobio nigrescens* | Sheraliev, BM | Sheraliev, BM | Qashqadaryo | Amu Darya | Qorasuv Stream | 38.570458 | 66.296166 | 16/08/2016 |
| 281. | Gobio_nigrescens_247 | SWU16072018247 | FFU276-21 | MW649430 | Gobionidae | *Gobio nigrescens* | Sheraliev, BM | Sheraliev, BM | Bukhara | Zeravshan | Zeravshan | 40.146944 | 64.891666 | 16/07/2018 |
| 282. | Gobio_nigrescens_429 | SWU27072019429 | FFU277-21 | MW649431 | Gobionidae | *Gobio nigrescens* | Sheraliev, BM | Sheraliev, BM | Bukhara | Zeravshan | Echkiliksoy | 40.267551 | 64.743333 | 27/07/2019 |
| 283. | Gobio_nigrescens_435 | SWU27072019435 | FFU278-21 | MW649432 | Gobionidae | *Gobio nigrescens* | Sheraliev, BM | Sheraliev, BM | Bukhara | Zeravshan | Echkiliksoy | 40.267551 | 64.743333 | 27/07/2019 |
| 284. | Gobio_nigrescens_436 | SWU27072019436 | FFU279-21 | MW649433 | Gobionidae | *Gobio nigrescens* | Sheraliev, BM | Sheraliev, BM | Bukhara | Zeravshan | Echkiliksoy | 40.267551 | 64.743333 | 27/07/2019 |
| 285. | Gobio_nigrescens_531 | SWU02082019531 | FFU280-21 | MN810112 | Gobionidae | *Gobio nigrescens* | Sheraliev, BM | Sheraliev, BM | Samarkand | Zeravshan | Zeravshan | 39.676388 | 67.076111 | 02/08/2019 |
| 286. | Gobio_nigrescens_532 | SWU02082019532 | FFU281-21 | MW649434 | Gobionidae | *Gobio nigrescens* | Sheraliev, BM | Sheraliev, BM | Samarkand | Zeravshan | Zeravshan | 39.676388 | 67.076111 | 02/08/2019 |
| 287. | Gobio_nigrescens_533 | SWU02082019533 | FFU282-21 | MW649435 | Gobionidae | *Gobio nigrescens* | Sheraliev, BM | Sheraliev, BM | Samarkand | Zeravshan | Zeravshan | 39.676388 | 67.076111 | 02/08/2019 |
| 288. | Gobio_sibiricus_373 | SWU23072019373 | FFU283-21 | MN810110 | Gobionidae | *Gobio sibiricus* | Sheraliev, BM | Sheraliev, BM | Surxondaryo | Amu Darya | Tupalang River | 38.318966 | 68.006169 | 23/07/2019 |
| 289. | Gobio_sibiricus_383 | SWU23072019383 | FFU284-21 | MW649436 | Gobionidae | *Gobio sibiricus* | Sheraliev, BM | Sheraliev, BM | Surxondaryo | Amu Darya | Tupalang River | 38.318966 | 68.006169 | 23/07/2019 |
| 290. | Pseudorasbora_parva_064 | SWU07012017064 | FFU285-21 | MW649437 | Gobionidae | *Pseudorasbora parva* | Sheraliev, BM | Sheraliev, BM | Fergana | Syr Darya | Unnamed stream | 40.366437 | 71.316907 | 07/01/2017 |
| 291. | Pseudorasbora_parva_065 | SWU07012017065 | FFU286-21 | MW649438 | Gobionidae | *Pseudorasbora parva* | Sheraliev, BM | Sheraliev, BM | Fergana | Syr Darya | Unnamed stream | 40.366437 | 71.316907 | 07/01/2017 |
| 292. | Pseudorasbora_parva_101 | SWU25012018101 | FFU287-21 | MW649439 | Gobionidae | *Pseudorasbora parva* | Sheraliev, BM | Sheraliev, BM | Fergana | Syr Darya | Fergana Canal | 40.388497 | 71.327960 | 25/01/2018 |
| 293. | Pseudorasbora_parva_102 | SWU25012018102 | FFU288-21 | MW649440 | Gobionidae | *Pseudorasbora parva* | Sheraliev, BM | Sheraliev, BM | Fergana | Syr Darya | Fergana Canal | 40.388497 | 71.327960 | 25/01/2018 |
| 294. | Pseudorasbora_parva_103 | SWU25012018103 | FFU289-21 | MW649441 | Gobionidae | *Pseudorasbora parva* | Sheraliev, BM | Sheraliev, BM | Fergana | Syr Darya | Fergana Canal | 40.388497 | 71.327960 | 25/01/2018 |
| 295. | Pseudorasbora_parva_144 | SWU21022018144 | FFU290-21 | MW649442 | Gobionidae | *Pseudorasbora parva* | Sheraliev, BM | Sheraliev, BM | Tashkent | Syr Darya | Chirchik River | 41.370266 | 69.512026 | 21/02/2018 |

| | | | | | | | | | | | | | | |
|---|---|---|---|---|---|---|---|---|---|---|---|---|---|---|
| 296. | Pseudorasbora_parva_266 | SWU19072019266 | FFU291-21 | MW649443 | Gobionidae | *Pseudorasbora parva* | Sheraliev, BM | Sheraliev, BM | Surxondaryo | Amu Darya | Surkhan Darya | 37.255833 | 67.338611 | 19/07/2019 |
| 297. | Pseudorasbora_parva_309 | SWU21072019309 | FFU292-21 | MW649444 | Gobionidae | *Pseudorasbora parva* | Sheraliev, BM | Sheraliev, BM | Surxondaryo | Amu Darya | Surkhan Darya | 37.238888 | 67.329722 | 21/07/2019 |
| 298. | Pseudorasbora_parva_442 | SWU27072019442 | FFU293-21 | MW649445 | Gobionidae | *Pseudorasbora parva* | Sheraliev, BM | Sheraliev, BM | Bukhara | Zeravshan | Echkiliksoy | 40.267551 | 64.743333 | 27/07/2019 |
| 299. | Pseudorasbora_parva_580 | SWU09082019580 | FFU294-21 | MW649446 | Gobionidae | *Pseudorasbora parva* | Sheraliev, BM | Sheraliev, BM | Namangan | Syr Darya | Kara Darya | 40.918813 | 71.828896 | 09/08/2019 |
| 300. | Pseudorasbora_parva_692 | SWU27042019692 | FFU295-21 | MW649447 | Gobionidae | *Pseudorasbora parva* | Allayarov, S | Sheraliev, BM | Surxondaryo | Amu Darya | Karatag river | 38.363055 | 68.072059 | 27/04/2019 |
| 301. | Abramis_brama_180 | SWU18072018180 | FFU296-21 | MW649448 | Leuciscidae | *Abramis brama* | Sheraliev, BM | Sheraliev, BM | Navoiy | Zeravshan | Tudakul Lake | 39.801388 | 64.772222 | 18/07/2018 |
| 302. | Abramis_brama_181 | SWU18072018181 | FFU297-21 | MW649449 | Leuciscidae | *Abramis brama* | Sheraliev, BM | Sheraliev, BM | Navoiy | Zeravshan | Tudakul Lake | 39.801388 | 64.772222 | 18/07/2018 |
| 303. | Alburnoides_holciki_001 | SWU08082016001 | FFU298-21 | MN872408 | Leuciscidae | *Alburnoides holciki* | Sheraliev, BM | Sheraliev, BM | Surxondaryo | Amu Darya | Tupalang river | 38.399802 | 67.944766 | 08/08/2016 |
| 304. | Alburnoides_holciki_002 | SWU08082016002 | FFU299-21 | MW649450 | Leuciscidae | *Alburnoides holciki* | Sheraliev, BM | Sheraliev, BM | Surxondaryo | Amu Darya | Tupalang river | 38.399802 | 67.944766 | 08/08/2016 |
| 305. | Alburnoides_holciki_003 | SWU08082016003 | FFU300-21 | MW649451 | Leuciscidae | *Alburnoides holciki* | Sheraliev, BM | Sheraliev, BM | Surxondaryo | Amu Darya | Tupalang river | 38.399802 | 67.944766 | 08/08/2016 |
| 306. | Alburnoides_holciki_155 | SWU16072018155 | FFU301-21 | MW649452 | Leuciscidae | *Alburnoides holciki* | Sheraliev, BM | Sheraliev, BM | Bukhara | Zeravshan | Zeravshan | 40.146944 | 64.891666 | 16/07/2018 |
| 307. | Alburnoides_holciki_156 | SWU16072018156 | FFU302-21 | MW649453 | Leuciscidae | *Alburnoides holciki* | Sheraliev, BM | Sheraliev, BM | Bukhara | Zeravshan | Zeravshan | 40.146944 | 64.891666 | 16/07/2018 |
| 308. | Alburnoides_holciki_157 | SWU16072018157 | FFU303-21 | MW649454 | Leuciscidae | *Alburnoides holciki* | Sheraliev, BM | Sheraliev, BM | Bukhara | Zeravshan | Zeravshan | 40.146944 | 64.891666 | 16/07/2018 |
| 309. | Alburnoides_holciki_158 | SWU16072018158 | FFU304-21 | MW649455 | Leuciscidae | *Alburnoides holciki* | Sheraliev, BM | Sheraliev, BM | Bukhara | Zeravshan | Zeravshan | 40.146944 | 64.891666 | 16/07/2018 |
| 310. | Alburnoides_holciki_159 | SWU16072018159 | FFU305-21 | MW649456 | Leuciscidae | *Alburnoides holciki* | Sheraliev, BM | Sheraliev, BM | Bukhara | Zeravshan | Zeravshan | 40.146944 | 64.891666 | 16/07/2018 |
| 311. | Alburnoides_holciki_236 | SWU12082016236 | FFU306-21 | MW649457 | Leuciscidae | *Alburnoides holciki* | Sheraliev, BM | Sheraliev, BM | Surxondaryo | Amu Darya | Tupalang | 38.376388 | 67.963166 | 12/08/2016 |
| 312. | Alburnoides_holciki_237 | SWU12082016237 | FFU307-21 | MW649458 | Leuciscidae | *Alburnoides holciki* | Sheraliev, BM | Sheraliev, BM | Surxondaryo | Amu Darya | Tupalang | 38.376388 | 67.963166 | 12/08/2016 |
| 313. | Alburnoides_holciki_238 | SWU12082016238 | FFU308-21 | MW649459 | Leuciscidae | *Alburnoides holciki* | Sheraliev, BM | Sheraliev, BM | Surxondaryo | Amu Darya | Tupalang | 38.376388 | 67.963166 | 12/08/2016 |
| 314. | Alburnoides_holciki_250 | SWU16072018250 | FFU309-21 | MW649460 | Leuciscidae | *Alburnoides holciki* | Sheraliev, BM | Sheraliev, BM | Bukhara | Zeravshan | Zeravshan | 40.146944 | 64.891666 | 16/07/2018 |
| 315. | Alburnoides_holciki_251 | SWU16072018251 | FFU310-21 | MW649461 | Leuciscidae | *Alburnoides holciki* | Sheraliev, BM | Sheraliev, BM | Bukhara | Zeravshan | Zeravshan | 40.146944 | 64.891666 | 16/07/2018 |
| 316. | Alburnoides_holciki_267 | SWU19072019267 | FFU311-21 | MW649462 | Leuciscidae | *Alburnoides holciki* | Sheraliev, BM | Sheraliev, BM | Surxondaryo | Amu Darya | Surkhan Darya | 37.255833 | 67.338611 | 19/07/2019 |
| 317. | Alburnoides_holciki_268 | SWU19072019268 | FFU312-21 | MW649463 | Leuciscidae | *Alburnoides holciki* | Sheraliev, BM | Sheraliev, BM | Surxondaryo | Amu Darya | Surkhan Darya | 37.255833 | 67.338611 | 19/07/2019 |
| 318. | Alburnoides_holciki_269 | SWU19072019269 | FFU313-21 | MW649464 | Leuciscidae | *Alburnoides holciki* | Sheraliev, BM | Sheraliev, BM | Surxondaryo | Amu Darya | Surkhan Darya | 37.255833 | 67.338611 | 19/07/2019 |
| 319. | Alburnoides_holciki_270 | SWU19072019270 | FFU314-21 | MW649465 | Leuciscidae | *Alburnoides holciki* | Sheraliev, BM | Sheraliev, BM | Surxondaryo | Amu Darya | Surkhan Darya | 37.255833 | 67.338611 | 19/07/2019 |
| 320. | Alburnoides_holciki_275 | SWU19072019275 | FFU315-21 | MW649466 | Leuciscidae | *Alburnoides holciki* | Sheraliev, BM | Sheraliev, BM | Surxondaryo | Amu Darya | Surkhan Darya | 37.255833 | 67.338611 | 19/07/2019 |
| 321. | Alburnoides_holciki_276 | SWU19072019276 | FFU316-21 | MW649467 | Leuciscidae | *Alburnoides holciki* | Sheraliev, BM | Sheraliev, BM | Surxondaryo | Amu Darya | Surkhan Darya | 37.255833 | 67.338611 | 19/07/2019 |
| 322. | Alburnoides_holciki_277 | SWU19072019277 | FFU317-21 | MW649468 | Leuciscidae | *Alburnoides holciki* | Sheraliev, BM | Sheraliev, BM | Surxondaryo | Amu Darya | Surkhan Darya | 37.255833 | 67.338611 | 19/07/2019 |
| 323. | Alburnoides_holciki_278 | SWU19072019278 | FFU318-21 | MW649469 | Leuciscidae | *Alburnoides holciki* | Sheraliev, BM | Sheraliev, BM | Surxondaryo | Amu Darya | Surkhan Darya | 37.255833 | 67.338611 | 19/07/2019 |
| 324. | Alburnoides_holciki_279 | SWU19072019279 | FFU319-21 | MW649470 | Leuciscidae | *Alburnoides holciki* | Sheraliev, BM | Sheraliev, BM | Surxondaryo | Amu Darya | Surkhan Darya | 37.255833 | 67.338611 | 19/07/2019 |
| 325. | Alburnoides_holciki_321 | SWU21072019321 | FFU320-21 | MW649471 | Leuciscidae | *Alburnoides holciki* | Sheraliev, BM | Sheraliev, BM | Surxondaryo | Amu Darya | Surkhan Darya | 37.238888 | 67.329722 | 21/07/2019 |
| 326. | Alburnoides_holciki_323 | SWU21072019323 | FFU321-21 | MW649472 | Leuciscidae | *Alburnoides holciki* | Sheraliev, BM | Sheraliev, BM | Surxondaryo | Amu Darya | Surkhan Darya | 37.238888 | 67.329722 | 21/07/2019 |
| 327. | Alburnoides_holciki_369 | SWU19072019369 | FFU322-21 | MW649473 | Leuciscidae | *Alburnoides holciki* | Sheraliev, BM | Sheraliev, BM | Surxondaryo | Amu Darya | Surkhan Darya | 37.255833 | 67.338611 | 19/07/2019 |
| 328. | Alburnoides_holciki_370 | SWU19072019370 | FFU323-21 | MW649474 | Leuciscidae | *Alburnoides holciki* | Sheraliev, BM | Sheraliev, BM | Surxondaryo | Amu Darya | Surkhan Darya | 37.255833 | 67.338611 | 19/07/2019 |
| 329. | Alburnoides_holciki_409 | SWU25072019409 | FFU324-21 | MW649475 | Leuciscidae | *Alburnoides holciki* | Sheraliev, BM | Sheraliev, BM | Surxondaryo | Amu Darya | Sherobod river | 37.747222 | 66.996666 | 25/07/2019 |
| 330. | Alburnoides_holciki_410 | SWU25072019410 | FFU325-21 | MW649476 | Leuciscidae | *Alburnoides holciki* | Sheraliev, BM | Sheraliev, BM | Surxondaryo | Amu Darya | Sherobod river | 37.747222 | 66.996666 | 25/07/2019 |
| 331. | Alburnoides_holciki_411 | SWU25072019411 | FFU326-21 | MW649477 | Leuciscidae | *Alburnoides holciki* | Sheraliev, BM | Sheraliev, BM | Surxondaryo | Amu Darya | Sherobod river | 37.747222 | 66.996666 | 25/07/2019 |
| 332. | Alburnoides_holciki_426 | SWU27072019426 | FFU327-21 | MW649478 | Leuciscidae | *Alburnoides holciki* | Sheraliev, BM | Sheraliev, BM | Bukhara | Zeravshan | Echkiliksoy | 40.267551 | 64.743333 | 27/07/2019 |
| 333. | Alburnoides_holciki_427 | SWU27072019427 | FFU328-21 | MW649479 | Leuciscidae | *Alburnoides holciki* | Sheraliev, BM | Sheraliev, BM | Bukhara | Zeravshan | Echkiliksoy | 40.267551 | 64.743333 | 27/07/2019 |
| 334. | Alburnoides_holciki_443 | SWU27072019443 | FFU329-21 | MW649480 | Leuciscidae | *Alburnoides holciki* | Sheraliev, BM | Sheraliev, BM | Bukhara | Zeravshan | Echkiliksoy | 40.267551 | 64.743333 | 27/07/2019 |
| 335. | Alburnoides_holciki_444 | SWU27072019444 | FFU330-21 | MW649481 | Leuciscidae | *Alburnoides holciki* | Sheraliev, BM | Sheraliev, BM | Bukhara | Zeravshan | Echkiliksoy | 40.267551 | 64.743333 | 27/07/2019 |
| 336. | Alburnoides_holciki_445 | SWU27072019445 | FFU331-21 | MW649482 | Leuciscidae | *Alburnoides holciki* | Sheraliev, BM | Sheraliev, BM | Bukhara | Zeravshan | Echkiliksoy | 40.267551 | 64.743333 | 27/07/2019 |
| 337. | Alburnoides_holciki_489 | SWU31072019489 | FFU332-21 | MW649483 | Leuciscidae | *Alburnoides holciki* | Sheraliev, BM | Sheraliev, BM | Khorazm | Amu Darya | Amu Darya | 41.650833 | 60.704722 | 31/07/2019 |
| 338. | Alburnoides_holciki_536 | SWU02082019536 | FFU333-21 | MW649484 | Leuciscidae | *Alburnoides holciki* | Sheraliev, BM | Sheraliev, BM | Samarkand | Zeravshan | Zeravshan | 39.676388 | 67.076111 | 02/08/2019 |
| 339. | Alburnoides_holciki_537 | SWU02082019537 | FFU334-21 | MW649485 | Leuciscidae | *Alburnoides holciki* | Sheraliev, BM | Sheraliev, BM | Samarkand | Zeravshan | Zeravshan | 39.676388 | 67.076111 | 02/08/2019 |
| 340. | Alburnoides_holciki_538 | SWU02082019538 | FFU335-21 | MW649486 | Leuciscidae | *Alburnoides holciki* | Sheraliev, BM | Sheraliev, BM | Samarkand | Zeravshan | Zeravshan | 39.676388 | 67.076111 | 02/08/2019 |
| 341. | Alburnoides_holciki_697 | SWU27042019697 | FFU336-21 | MW649487 | Leuciscidae | *Alburnoides holciki* | Allayarov, S | Sheraliev, BM | Surxondaryo | Amu Darya | Karatag river | 38.363055 | 68.072059 | 27/04/2019 |
| 342. | Alburnoides_holciki_698 | SWU27042019698 | FFU337-21 | MW649488 | Leuciscidae | *Alburnoides holciki* | Allayarov, S | Sheraliev, BM | Surxondaryo | Amu Darya | Karatag river | 38.363055 | 68.072059 | 27/04/2019 |
| 343. | Alburnoides_holciki_699 | SWU27042019699 | FFU338-21 | MW649489 | Leuciscidae | *Alburnoides holciki* | Allayarov, S | Sheraliev, BM | Surxondaryo | Amu Darya | Karatag river | 38.363055 | 68.072059 | 27/04/2019 |
| 344. | Alburnoides_taeniatus_430 | SWU27072019430 | FFU339-21 | MW649490 | Leuciscidae | *Alburnus taeniatus* | Sheraliev, BM | Sheraliev, BM | Bukhara | Zeravshan | Echkiliksoy | 40.267551 | 64.743333 | 27/07/2019 |
| 345. | Alburnoides_taeniatus_534 | SWU02082019534 | FFU340-21 | MW649491 | Leuciscidae | *Alburnus taeniatus* | Sheraliev, BM | Sheraliev, BM | Samarkand | Zeravshan | Zeravshan | 39.676388 | 67.076111 | 02/08/2019 |
| 346. | Alburnoides_taeniatus_535 | SWU02082019535 | FFU341-21 | MW649492 | Leuciscidae | *Alburnus taeniatus* | Sheraliev, BM | Sheraliev, BM | Samarkand | Zeravshan | Zeravshan | 39.676388 | 67.076111 | 02/08/2019 |
| 347. | Alburnus_chalcoides_016 | SWU15082016016 | FFU342-21 | MW649493 | Leuciscidae | *Alburnus chalcoides* | Sheraliev, BM | Sheraliev, BM | Qashqadaryo | Amu Darya | Qorasuv Stream | 38.570458 | 66.296166 | 15/08/2016 |
| 348. | Alburnus_chalcoides_017 | SWU15082016017 | FFU343-21 | MW649494 | Leuciscidae | *Alburnus chalcoides* | Sheraliev, BM | Sheraliev, BM | Qashqadaryo | Amu Darya | Qorasuv Stream | 38.570458 | 66.296166 | 15/08/2016 |
| 349. | Alburnus_chalcoides_018 | SWU15082016018 | FFU344-21 | MW649495 | Leuciscidae | *Alburnus chalcoides* | Sheraliev, BM | Sheraliev, BM | Qashqadaryo | Amu Darya | Qorasuv Stream | 38.570458 | 66.296166 | 15/08/2016 |
| 350. | Alburnus_chalcoides_177 | SWU18072018177 | FFU345-21 | MW649496 | Leuciscidae | *Alburnus chalcoides* | Sheraliev, BM | Sheraliev, BM | Navoiy | Zeravshan | Tudakul Lake | 39.801388 | 64.772222 | 18/07/2018 |
| 351. | Alburnus_chalcoides_274 | SWU19072019274 | FFU346-21 | MW649497 | Leuciscidae | *Alburnus chalcoides* | Sheraliev, BM | Sheraliev, BM | Surxondaryo | Amu Darya | Surkhan Darya | 37.255833 | 67.338611 | 19/07/2019 |
| 352. | Alburnus_chalcoides_280 | SWU19072019280 | FFU347-21 | MW649498 | Leuciscidae | *Alburnus chalcoides* | Sheraliev, BM | Sheraliev, BM | Surxondaryo | Amu Darya | Surkhan Darya | 37.255833 | 67.338611 | 19/07/2019 |
| 353. | Alburnus_chalcoides_431 | SWU27072019431 | FFU348-21 | MW649499 | Leuciscidae | *Alburnus chalcoides* | Sheraliev, BM | Sheraliev, BM | Bukhara | Zeravshan | Echkiliksoy | 40.267551 | 64.743333 | 27/07/2019 |
| 354. | Alburnus_chalcoides_440 | SWU27072019440 | FFU349-21 | MW649500 | Leuciscidae | *Alburnus chalcoides* | Sheraliev, BM | Sheraliev, BM | Bukhara | Zeravshan | Echkiliksoy | 40.267551 | 64.743333 | 27/07/2019 |
| 355. | Alburnus_chalcoides_441 | SWU27072019441 | FFU350-21 | MW649501 | Leuciscidae | *Alburnus chalcoides* | Sheraliev, BM | Sheraliev, BM | Bukhara | Zeravshan | Echkiliksoy | 40.267551 | 64.743333 | 27/07/2019 |
| 356. | Alburnus_chalcoides_471 | SWU29072019471 | FFU351-21 | MW649502 | Leuciscidae | *Alburnus chalcoides* | Sheraliev, BM | Sheraliev, BM | Bukhara | Zeravshan | Unnamed stream | 40.147222 | 64.905805 | 29/07/2019 |
| 357. | Alburnus_chalcoides_472 | SWU29072019472 | FFU352-21 | MW649503 | Leuciscidae | *Alburnus chalcoides* | Sheraliev, BM | Sheraliev, BM | Bukhara | Zeravshan | Unnamed stream | 40.147222 | 64.905805 | 29/07/2019 |
| 358. | Alburnus_chalcoides_473 | SWU29072019473 | FFU353-21 | MW649504 | Leuciscidae | *Alburnus chalcoides* | Sheraliev, BM | Sheraliev, BM | Bukhara | Zeravshan | Unnamed stream | 40.147222 | 64.905805 | 29/07/2019 |
| 359. | Alburnus_oblongus_591 | SWU18082019591 | FFU354-21 | MW649505 | Leuciscidae | *Alburnus oblongus* | Sheraliev, BM | Sheraliev, BM | Tashkent | Syr Darya | Chirchik river | 41.274444 | 69.389166 | 18/08/2019 |
| 360. | Alburnus_oblongus_592 | SWU18082019592 | FFU355-21 | MW649506 | Leuciscidae | *Alburnus oblongus* | Sheraliev, BM | Sheraliev, BM | Tashkent | Syr Darya | Chirchik river | 41.274444 | 69.389166 | 18/08/2019 |
| 361. | Alburnus_oblongus_593 | SWU18082019593 | FFU356-21 | MW649507 | Leuciscidae | *Alburnus oblongus* | Sheraliev, BM | Sheraliev, BM | Tashkent | Syr Darya | Chirchik river | 41.274444 | 69.389166 | 18/08/2019 |
| 362. | Alburnus_oblongus_594 | SWU18082019594 | FFU357-21 | MW649508 | Leuciscidae | *Alburnus oblongus* | Sheraliev, BM | Sheraliev, BM | Tashkent | Syr Darya | Chirchik river | 41.274444 | 69.389166 | 18/08/2019 |
| 363. | Alburnus_oblongus_604 | SWU18082019604 | FFU358-21 | MW649509 | Leuciscidae | *Alburnus oblongus* | Sheraliev, BM | Sheraliev, BM | Tashkent | Syr Darya | Chirchik river | 41.450555 | 69.598611 | 18/08/2019 |
| 364. | Alburnus_oblongus_605 | SWU18082019605 | FFU359-21 | MW649510 | Leuciscidae | *Alburnus oblongus* | Sheraliev, BM | Sheraliev, BM | Tashkent | Syr Darya | Chirchik river | 41.450555 | 69.598611 | 18/08/2019 |
| 365. | Alburnus_oblongus_606 | SWU18082019606 | FFU360-21 | MW649511 | Leuciscidae | *Alburnus oblongus* | Sheraliev, BM | Sheraliev, BM | Tashkent | Syr Darya | Chirchik river | 41.450555 | 69.598611 | 18/08/2019 |
| 366. | Alburnus_oblongus_607 | SWU18082019607 | FFU361-21 | MW649512 | Leuciscidae | *Alburnus oblongus* | Sheraliev, BM | Sheraliev, BM | Tashkent | Syr Darya | Chirchik river | 41.450555 | 69.598611 | 18/08/2019 |
| 367. | Capoetobrama_kuschakewitschi_314 | SWU21072019313 | FFU362-21 | MW649513 | Leuciscidae | *Capoetobrama kuschakewitschi* | Allayarov, S | Sheraliev, BM | Surxondaryo | Amu Darya | Amu Darya | 37.235555 | 67.660555 | 21/07/2019 |
| 368. | Capoetobrama_kuschakewitschi_315 | SWU21072019313 | FFU363-21 | MW649514 | Leuciscidae | *Capoetobrama kuschakewitschi* | Allayarov, S | Sheraliev, BM | Surxondaryo | Amu Darya | Amu Darya | 37.235555 | 67.660555 | 21/07/2019 |
| 369. | Capoetobrama_kuschakewitschi_316 | SWU21072019313 | FFU364-21 | MW649515 | Leuciscidae | *Capoetobrama kuschakewitschi* | Allayarov, S | Sheraliev, BM | Surxondaryo | Amu Darya | Amu Darya | 37.235555 | 67.660555 | 21/07/2019 |

| | | | | | | | | | | | | | | |
|---|---|---|---|---|---|---|---|---|---|---|---|---|---|---|
| 370. | Capoetobrama_kuschakewitschi_317 | SWU21072019313 | FFU365-21 | MW649516 | Leuciscidae | *Capoetobrama kuschakewitschi* | Allayarov, S | Sheraliev, BM | Surxondaryo | Amu Darya | Amu Darya | 37.235555 | 67.660555 | 21/07/2019 |
| 371. | Capoetobrama_kuschakewitschi_491 | SWU31072019491 | FFU366-21 | MW649517 | Leuciscidae | *Capoetobrama kuschakewitschi* | Sheraliev, BM | Sheraliev, BM | Khorazm | Amu Darya | Amu Darya | 41.650833 | 60.704722 | 31/07/2019 |
| 372. | Capoetobrama_kuschakewitschi_492 | SWU31072019492 | FFU367-21 | MW649518 | Leuciscidae | *Capoetobrama kuschakewitschi* | Sheraliev, BM | Sheraliev, BM | Khorazm | Amu Darya | Amu Darya | 41.650833 | 60.704722 | 31/07/2019 |
| 373. | Capoetobrama_kuschakewitschi_499 | SWU31072019499 | FFU368-21 | MW649519 | Leuciscidae | *Capoetobrama kuschakewitschi* | Sheraliev, BM | Sheraliev, BM | Khorazm | Amu Darya | Amu Darya | 41.650833 | 60.704722 | 31/07/2019 |
| 374. | Capoetobrama_kuschakewitschi_501 | SWU31072019501 | FFU369-21 | MW649520 | Leuciscidae | *Capoetobrama kuschakewitschi* | Sheraliev, BM | Sheraliev, BM | Khorazm | Amu Darya | Amu Darya | 41.650833 | 60.704722 | 31/07/2019 |
| 375. | Leuciscus_aspius_673 | SWU21082019673 | FFU370-21 | MW649521 | Leuciscidae | *Leuciscus aspius* | Rozimov, A | Sheraliev, BM | Khorazm | Amu Darya | Amu Darya | 41.948638 | 60.444194 | 21/08/2019 |
| 376. | Leuciscus_lehmanni_015 | SWU15082016015 | FFU371-21 | MN872388 | Leuciscidae | *Leuciscus lehmanni* | Sheraliev, BM | Sheraliev, BM | Qashqadaryo | Amu Darya | Qorasuv Stream | 38.570458 | 66.296166 | 15/08/2016 |
| 377. | Leuciscus_lehmanni_133 | SWU22022018133 | FFU372-21 | MN872389 | Leuciscidae | *Leuciscus lehmanni* | Sheraliev, BM | Sheraliev, BM | Bukhara | Zeravshan | Qoraqir Lake | 40.385644 | 63.308388 | 22/02/2018 |
| 378. | Leuciscus_lehmanni_271 | SWU19072019271 | FFU373-21 | MN872390 | Leuciscidae | *Leuciscus lehmanni* | Sheraliev, BM | Sheraliev, BM | Surxondaryo | Amu Darya | Surkhan Darya | 37.255833 | 67.338611 | 19/07/2019 |
| 379. | Leuciscus_lehmanni_272 | SWU19072019271 | FFU374-21 | MN872391 | Leuciscidae | *Leuciscus lehmanni* | Sheraliev, BM | Sheraliev, BM | Surxondaryo | Amu Darya | Surkhan Darya | 37.255833 | 67.338611 | 19/07/2019 |
| 380. | Leuciscus_lehmanni_273 | SWU19072019271 | FFU375-21 | MN872392 | Leuciscidae | *Leuciscus lehmanni* | Sheraliev, BM | Sheraliev, BM | Surxondaryo | Amu Darya | Surkhan Darya | 37.255833 | 67.338611 | 19/07/2019 |
| 381. | Leuciscus_lehmanni_295 | SWU21072019295 | FFU376-21 | MN872393 | Leuciscidae | *Leuciscus lehmanni* | Sheraliev, BM | Sheraliev, BM | Surxondaryo | Amu Darya | Surkhan Darya | 37.238888 | 67.329722 | 21/07/2019 |
| 382. | Leuciscus_lehmanni_304 | SWU21072019304 | FFU377-21 | MN872394 | Leuciscidae | *Leuciscus lehmanni* | Sheraliev, BM | Sheraliev, BM | Surxondaryo | Amu Darya | Surkhan Darya | 37.238888 | 67.329722 | 21/07/2019 |
| 383. | Leuciscus_lehmanni_305 | SWU21072019305 | FFU378-21 | MN872395 | Leuciscidae | *Leuciscus lehmanni* | Sheraliev, BM | Sheraliev, BM | Surxondaryo | Amu Darya | Surkhan Darya | 37.238888 | 67.329722 | 21/07/2019 |
| 384. | Leuciscus_lehmanni_475 | SWU29072019475 | FFU379-21 | MN872396 | Leuciscidae | *Leuciscus lehmanni* | Sheraliev, BM | Sheraliev, BM | Bukhara | Zeravshan | Unnamed stream | 40.147222 | 64.905805 | 29/07/2019 |
| 385. | Leuciscus_lehmanni_476 | SWU29072019475 | FFU380-21 | MN872397 | Leuciscidae | *Leuciscus lehmanni* | Sheraliev, BM | Sheraliev, BM | Bukhara | Zeravshan | Unnamed stream | 40.147222 | 64.905805 | 29/07/2019 |
| 386. | Pelecus_cultratus_036 | SWU31082016036 | FFU381-21 | MW649522 | Leuciscidae | *Pelecus cultratus* | Sheraliev, BM | Sheraliev, BM | Namangan | Syr Darya | Syr Darya | 40.882777 | 71.442222 | 31/08/2016 |
| 387. | Pelecus_cultratus_037 | SWU31082016037 | FFU382-21 | MW649523 | Leuciscidae | *Pelecus cultratus* | Sheraliev, BM | Sheraliev, BM | Namangan | Syr Darya | Syr Darya | 40.882777 | 71.442222 | 31/08/2016 |
| 388. | Pelecus_cultratus_038 | SWU31082016038 | FFU383-21 | MW649524 | Leuciscidae | *Pelecus cultratus* | Sheraliev, BM | Sheraliev, BM | Namangan | Syr Darya | Syr Darya | 40.882777 | 71.442222 | 31/08/2016 |
| 389. | Pelecus_cultratus_039 | SWU31082016039 | FFU384-21 | MW649525 | Leuciscidae | *Pelecus cultratus* | Sheraliev, BM | Sheraliev, BM | Namangan | Syr Darya | Syr Darya | 40.882777 | 71.442222 | 31/08/2016 |
| 390. | Petroleuciscus_squaliusculus_068 | SWU15012017068 | FFU385-21 | MN872398 | Leuciscidae | *Leuciscus squaliusculus* | Sheraliev, BM | Sheraliev, BM | Fergana | Syr Darya | Unnamed stream | 40.326408 | 71.820508 | 15/01/2017 |
| 391. | Petroleuciscus_squaliusculus_069 | SWU15012017069 | FFU386-21 | MN872399 | Leuciscidae | *Leuciscus squaliusculus* | Sheraliev, BM | Sheraliev, BM | Fergana | Syr Darya | Unnamed stream | 40.326408 | 71.820508 | 15/01/2017 |
| 392. | Petroleuciscus_squaliusculus_070 | SWU15012017070 | FFU387-21 | MN872400 | Leuciscidae | *Leuciscus squaliusculus* | Sheraliev, BM | Sheraliev, BM | Fergana | Syr Darya | Unnamed stream | 40.326408 | 71.820508 | 15/01/2017 |
| 393. | Petroleuciscus_squaliusculus_128 | SWU08022018128 | FFU388-21 | MN872401 | Leuciscidae | *Leuciscus squaliusculus* | Sheraliev, BM | Sheraliev, BM | Fergana | Syr Darya | Sokh River | 40.340176 | 71.013559 | 08/02/2018 |
| 394. | Petroleuciscus_squaliusculus_191 | SWU11082018191 | FFU389-21 | MN872402 | Leuciscidae | *Leuciscus squaliusculus* | Sheraliev, BM | Sheraliev, BM | Fergana | Syr Darya | Unnamed stream | 40.312769 | 71.811647 | 11/08/2018 |
| 395. | Petroleuciscus_squaliusculus_581 | SWU09082019581 | FFU390-21 | MN872403 | Leuciscidae | *Leuciscus squaliusculus* | Sheraliev, BM | Sheraliev, BM | Namangan | Syr Darya | Naryn River | 40.940911 | 71.853631 | 09/08/2019 |
| 396. | Petroleuciscus_squaliusculus_582 | SWU09082019582 | FFU391-21 | MN872404 | Leuciscidae | *Leuciscus squaliusculus* | Sheraliev, BM | Sheraliev, BM | Namangan | Syr Darya | Naryn River | 40.940911 | 71.853631 | 09/08/2019 |
| 397. | Petroleuciscus_squaliusculus_639 | SWU20082019639 | FFU392-21 | MN872405 | Leuciscidae | *Leuciscus squaliusculus* | Sheraliev, BM | Sheraliev, BM | Fergana | Syr Darya | Unnamed stream | 40.305260 | 71.800688 | 20/08/2019 |
| 398. | Petroleuciscus_squaliusculus_640 | SWU20082019640 | FFU393-21 | MN872406 | Leuciscidae | *Leuciscus squaliusculus* | Sheraliev, BM | Sheraliev, BM | Fergana | Syr Darya | Unnamed stream | 40.305260 | 71.800688 | 20/08/2019 |
| 399. | Petroleuciscus_squaliusculus_641 | SWU20082019641 | FFU394-21 | MN872407 | Leuciscidae | *Leuciscus squaliusculus* | Sheraliev, BM | Sheraliev, BM | Fergana | Syr Darya | Unnamed stream | 40.305260 | 71.800688 | 20/08/2019 |
| 400. | Rutilus_rutilus_aralensis_071 | SWU10072017071 | FFU395-21 | MW649526 | Leuciscidae | *Rutilus lacustris* | Sheraliev, BM | Sheraliev, BM | Fergana | Syr Darya | Unnamed stream | 40.588943 | 71.537333 | 10/07/2017 |
| 401. | Rutilus_rutilus_aralensis_072 | SWU10072017072 | FFU396-21 | MW649527 | Leuciscidae | *Rutilus lacustris* | Sheraliev, BM | Sheraliev, BM | Fergana | Syr Darya | Unnamed stream | 40.588943 | 71.537333 | 10/07/2017 |
| 402. | Rutilus_rutilus_aralensis_073 | SWU10072017073 | FFU397-21 | MW649528 | Leuciscidae | *Rutilus lacustris* | Sheraliev, BM | Sheraliev, BM | Fergana | Syr Darya | Unnamed stream | 40.588943 | 71.537333 | 10/07/2017 |
| 403. | Rutilus_rutilus_aralensis_074 | SWU10072017074 | FFU398-21 | MW649529 | Leuciscidae | *Rutilus lacustris* | Sheraliev, BM | Sheraliev, BM | Fergana | Syr Darya | Unnamed stream | 40.588943 | 71.537333 | 10/07/2017 |
| 404. | Rutilus_rutilus_aralensis_075 | SWU10072017075 | FFU399-21 | MW649530 | Leuciscidae | *Rutilus lacustris* | Sheraliev, BM | Sheraliev, BM | Fergana | Syr Darya | Unnamed stream | 40.588943 | 71.537333 | 10/07/2017 |
| 405. | Rutilus_rutilus_aralensis_076 | SWU10072017076 | FFU400-21 | MW649531 | Leuciscidae | *Rutilus lacustris* | Sheraliev, BM | Sheraliev, BM | Fergana | Syr Darya | Unnamed stream | 40.588943 | 71.537333 | 10/07/2017 |
| 406. | Rutilus_rutilus_aralensis_127 | SWU08022018127 | FFU401-21 | MW649532 | Leuciscidae | *Rutilus lacustris* | Sheraliev, BM | Sheraliev, BM | Fergana | Syr Darya | Sokh River | 40.340176 | 71.013559 | 08/02/2018 |
| 407. | Rutilus_rutilus_aralensis_447 | SWU28072019447 | FFU402-21 | MW649533 | Leuciscidae | *Rutilus lacustris* | Sheraliev, BM | Sheraliev, BM | Bukhara | Zeravshan | Zeravshan | 40.071012 | 64.778429 | 28/07/2019 |
| 408. | Rutilus_rutilus_aralensis_448 | SWU28072019448 | FFU403-21 | MW649534 | Leuciscidae | *Rutilus lacustris* | Sheraliev, BM | Sheraliev, BM | Bukhara | Zeravshan | Zeravshan | 40.071012 | 64.778429 | 28/07/2019 |
| 409. | Rutilus_rutilus_aralensis_449 | SWU28072019449 | FFU404-21 | MW649535 | Leuciscidae | *Rutilus lacustris* | Sheraliev, BM | Sheraliev, BM | Bukhara | Zeravshan | Zeravshan | 40.071012 | 64.778429 | 28/07/2019 |
| 410. | Rutilus_rutilus_aralensis_674 | SWU21082019674 | FFU405-21 | MW649536 | Leuciscidae | *Rutilus lacustris* | Rozimov, A | Sheraliev, BM | Khorazm | Amu Darya | Amu Darya | 41.948638 | 60.444194 | 21/08/2019 |
| 411. | Dzihunia_amudarjensis_286 | SWU20072019286 | FFU406-21 | MW649537 | Nemacheilidae | *Dzihunia amudarjensis* | Sheraliev, BM | Sheraliev, BM | Surxondaryo | Amu Darya | Amu Darya | 37.235555 | 67.660555 | 20/07/2019 |
| 412. | Dzihunia_amudarjensis_287 | SWU20072019287 | FFU407-21 | MW649538 | Nemacheilidae | *Dzihunia amudarjensis* | Sheraliev, BM | Sheraliev, BM | Surxondaryo | Amu Darya | Amu Darya | 37.235555 | 67.660555 | 20/07/2019 |
| 413. | Dzihunia_amudarjensis_288 | SWU20072019288 | FFU408-21 | MW649539 | Nemacheilidae | *Dzihunia amudarjensis* | Sheraliev, BM | Sheraliev, BM | Surxondaryo | Amu Darya | Amu Darya | 37.235555 | 67.660555 | 20/07/2019 |
| 414. | Dzihunia_amudarjensis_289 | SWU20072019289 | FFU409-21 | MW649540 | Nemacheilidae | *Dzihunia amudarjensis* | Sheraliev, BM | Sheraliev, BM | Surxondaryo | Amu Darya | Amu Darya | 37.235555 | 67.660555 | 20/07/2019 |
| 415. | Dzihunia_amudarjensis_290 | SWU20072019290 | FFU410-21 | MW649541 | Nemacheilidae | *Dzihunia amudarjensis* | Sheraliev, BM | Sheraliev, BM | Surxondaryo | Amu Darya | Amu Darya | 37.235555 | 67.660555 | 20/07/2019 |
| 416. | Dzihunia_amudarjensis_291 | SWU20072019291 | FFU411-21 | MW649542 | Nemacheilidae | *Dzihunia amudarjensis* | Sheraliev, BM | Sheraliev, BM | Surxondaryo | Amu Darya | Amu Darya | 37.235555 | 67.660555 | 20/07/2019 |
| 417. | Dzihunia_amudarjensis_292 | SWU20072019292 | FFU412-21 | MW649543 | Nemacheilidae | *Dzihunia amudarjensis* | Sheraliev, BM | Sheraliev, BM | Surxondaryo | Amu Darya | Amu Darya | 37.235555 | 67.660555 | 20/07/2019 |
| 418. | Dzihunia_amudarjensis_293 | SWU20072019293 | FFU413-21 | MW649544 | Nemacheilidae | *Dzihunia amudarjensis* | Sheraliev, BM | Sheraliev, BM | Surxondaryo | Amu Darya | Amu Darya | 37.235555 | 67.660555 | 20/07/2019 |
| 419. | Dzihunia_amudarjensis_294 | SWU20072019294 | FFU414-21 | MW649545 | Nemacheilidae | *Dzihunia amudarjensis* | Sheraliev, BM | Sheraliev, BM | Surxondaryo | Amu Darya | Amu Darya | 37.235555 | 67.660555 | 20/07/2019 |
| 420. | Dzihunia_amudarjensis_669 | SWU21082019669 | FFU415-21 | MW649546 | Nemacheilidae | *Dzihunia amudarjensis* | Rozimov, A | Sheraliev, BM | Khorazm | Amu Darya | Turangasaka Ch. | 41.734111 | 60.547944 | 21/08/2019 |
| 421. | Dzihunia_amudarjensis_700 | SWU27042019700 | FFU416-21 | MW649547 | Nemacheilidae | *Dzihunia amudarjensis* | Allayarov, S | Sheraliev, BM | Surxondaryo | Amu Darya | Karatag river | 38.363055 | 68.072059 | 27/04/2019 |
| 422. | Dzihunia_sp1_Karatag_359 | SWU22072019359 | FFU603-21 | MW649548 | Nemacheilidae | *Dzihunia* sp1 | Sheraliev, BM | Sheraliev, BM | Surxondaryo | Amu Darya | Karatag river | 38.363055 | 68.072059 | 22/07/2019 |
| 423. | Dzihunia_sp1_Karatag_360 | SWU22072019360 | FFU632-21 | MW649549 | Nemacheilidae | *Dzihunia* sp1 | Sheraliev, BM | Sheraliev, BM | Surxondaryo | Amu Darya | Karatag river | 38.363055 | 68.072059 | 22/07/2019 |
| 424. | Dzihunia_sp1_Karatag_361 | SWU22072019361 | FFU633-21 | MW649550 | Nemacheilidae | *Dzihunia* sp1 | Sheraliev, BM | Sheraliev, BM | Surxondaryo | Amu Darya | Karatag river | 38.363055 | 68.072059 | 22/07/2019 |
| 425. | Dzihunia_sp1_Karatag_363 | SWU22072019363 | FFU634-21 | MW649551 | Nemacheilidae | *Dzihunia* sp1 | Sheraliev, BM | Sheraliev, BM | Surxondaryo | Amu Darya | Karatag river | 38.363055 | 68.072059 | 22/07/2019 |
| 426. | Dzihunia_sp1_Karatag_364 | SWU22072019364 | FFU635-21 | MW649552 | Nemacheilidae | *Dzihunia* sp1 | Sheraliev, BM | Sheraliev, BM | Surxondaryo | Amu Darya | Karatag river | 38.363055 | 68.072059 | 22/07/2019 |
| 427. | Dzihunia_sp1_Karatag_365 | SWU22072019365 | FFU636-21 | MW649553 | Nemacheilidae | *Dzihunia* sp1 | Sheraliev, BM | Sheraliev, BM | Surxondaryo | Amu Darya | Karatag river | 38.363055 | 68.072059 | 22/07/2019 |
| 428. | Dzihunia_sp1_Karatag_366 | SWU22072019366 | FFU637-21 | MW649554 | Nemacheilidae | *Dzihunia* sp1 | Sheraliev, BM | Sheraliev, BM | Surxondaryo | Amu Darya | Karatag river | 38.363055 | 68.072059 | 22/07/2019 |
| 429. | Dzihunia_sp1_Karatag_367 | SWU22072019367 | FFU638-21 | MW649555 | Nemacheilidae | *Dzihunia* sp1 | Sheraliev, BM | Sheraliev, BM | Surxondaryo | Amu Darya | Karatag river | 38.363055 | 68.072059 | 22/07/2019 |
| 430. | Dzihunia_sp1_Karatag_368 | SWU22072019368 | FFU639-21 | MW649556 | Nemacheilidae | *Dzihunia* sp1 | Sheraliev, BM | Sheraliev, BM | Surxondaryo | Amu Darya | Karatag river | 38.363055 | 68.072059 | 22/07/2019 |
| 431. | Dzihunia_sp1_Tupalang_377 | SWU23072019377 | FFU640-21 | MW649557 | Nemacheilidae | *Dzihunia* sp1 | Sheraliev, BM | Sheraliev, BM | Surxondaryo | Amu Darya | Tupalang River | 38.318966 | 68.006169 | 23/07/2019 |
| 432. | Dzihunia_sp1_Tupalang_378 | SWU23072019378 | FFU641-21 | MW649558 | Nemacheilidae | *Dzihunia* sp1 | Sheraliev, BM | Sheraliev, BM | Surxondaryo | Amu Darya | Tupalang River | 38.318966 | 68.006169 | 23/07/2019 |
| 433. | Dzihunia_sp2_Karatag_362 | SWU22072019672 | FFU642-21 | MW649559 | Nemacheilidae | *Dzihunia* sp2 | Sheraliev, BM | Sheraliev, BM | Surxondaryo | Amu Darya | Karatag river | 38.363055 | 68.072059 | 22/07/2019 |
| 434. | Dzihunia_sp2_Sherobod_405 | SWU25072019405 | FFU643-21 | MW649560 | Nemacheilidae | *Dzihunia* sp2 | Sheraliev, BM | Sheraliev, BM | Surxondaryo | Amu Darya | Sherobod river | 37.747222 | 66.996666 | 25/07/2019 |
| 435. | Dzihunia_sp2_Sherobod_406 | SWU25072019406 | FFU644-21 | MW649561 | Nemacheilidae | *Dzihunia* sp2 | Sheraliev, BM | Sheraliev, BM | Surxondaryo | Amu Darya | Sherobod river | 37.747222 | 66.996666 | 25/07/2019 |
| 436. | Dzihunia_sp3_Chirchik_626 | SWU18082019626 | FFU645-21 | MW649562 | Nemacheilidae | *Dzihunia* sp3 | Sheraliev, BM | Sheraliev, BM | Tashkent | Syr Darya | Chirchik river | 41.450555 | 69.598611 | 18/08/2019 |
| 437. | Dzihunia_sp3_Chirchik_627 | SWU18082019627 | FFU646-21 | MW649563 | Nemacheilidae | *Dzihunia* sp3 | Sheraliev, BM | Sheraliev, BM | Tashkent | Syr Darya | Chirchik river | 41.450555 | 69.598611 | 18/08/2019 |
| 438. | Dzihunia_sp3_Chirchik_628 | SWU18082019628 | FFU647-21 | MW649564 | Nemacheilidae | *Dzihunia* sp3 | Sheraliev, BM | Sheraliev, BM | Tashkent | Syr Darya | Chirchik river | 41.450555 | 69.598611 | 18/08/2019 |
| 439. | Dzihunia_sp3_Chirchik_629 | SWU18082019629 | FFU648-21 | MW649565 | Nemacheilidae | *Dzihunia* sp3 | Sheraliev, BM | Sheraliev, BM | Tashkent | Syr Darya | Chirchik river | 41.450555 | 69.598611 | 18/08/2019 |
| 440. | Dzihunia_sp3_Chirchik_635 | SWU18082019635 | FFU649-21 | MW649566 | Nemacheilidae | *Dzihunia* sp3 | Sheraliev, BM | Sheraliev, BM | Tashkent | Syr Darya | Chirchik river | 41.450555 | 69.598611 | 18/08/2019 |
| 441. | Dzihunia_sp3_Chirchik_636 | SWU18082019636 | FFU650-21 | MW649567 | Nemacheilidae | *Dzihunia* sp3 | Sheraliev, BM | Sheraliev, BM | Tashkent | Syr Darya | Chirchik river | 41.450555 | 69.598611 | 18/08/2019 |

| | | | | | | | | | | | | | | |
|---|---|---|---|---|---|---|---|---|---|---|---|---|---|---|
| 442. | Dzihunia_sp3_Chirchik_637 | SWU18082019637 | FFU651-21 | MW649568 | Nemacheilidae | *Dzihunia* sp3 | Sheraliev, BM | Sheraliev, BM | Tashkent | Syr Darya | Chirchik river | 41.450555 | 69.598611 | 18/08/2019 |
| 443. | Dzihunia_sp3_Chirchik_638 | SWU18082019638 | FFU652-21 | MW649569 | Nemacheilidae | *Dzihunia* sp3 | Sheraliev, BM | Sheraliev, BM | Tashkent | Syr Darya | Chirchik river | 41.450555 | 69.598611 | 18/08/2019 |
| 444. | Paracobitis_longicauda_004 | SWU08082016004 | FFU417-21 | MW649570 | Nemacheilidae | *Paracobitis longicauda* | Sheraliev, BM | Sheraliev, BM | Surxondaryo | Amu Darya | Tupalang river | 38.399802 | 67.944766 | 08/08/2016 |
| 445. | Paracobitis_longicauda_005 | SWU08082016005 | FFU418-21 | MW649571 | Nemacheilidae | *Paracobitis longicauda* | Sheraliev, BM | Sheraliev, BM | Surxondaryo | Amu Darya | Tupalang river | 38.399802 | 67.944766 | 08/08/2016 |
| 446. | Paracobitis_longicauda_083 | SWU08082016083 | FFU419-21 | MW649572 | Nemacheilidae | *Paracobitis longicauda* | Sheraliev, BM | Sheraliev, BM | Surxondaryo | Amu Darya | Tupalang river | 38.399802 | 67.944766 | 08/08/2016 |
| 447. | Paracobitis_longicauda_242 | SWU08082016242 | FFU420-21 | MW649573 | Nemacheilidae | *Paracobitis longicauda* | Sheraliev, BM | Sheraliev, BM | Surxondaryo | Amu Darya | Tupalang river | 38.399802 | 67.944766 | 08/08/2016 |
| 448. | Paracobitis_longicauda_325 | SWU21072019325 | FFU421-21 | MW649574 | Nemacheilidae | *Paracobitis longicauda* | Sheraliev, BM | Sheraliev, BM | Surxondaryo | Amu Darya | Surkhan Darya | 37.238888 | 67.329722 | 21/07/2019 |
| 449. | Paracobitis_longicauda_344 | SWU22072019344 | FFU422-21 | MW649575 | Nemacheilidae | *Paracobitis longicauda* | Sheraliev, BM | Sheraliev, BM | Surxondaryo | Amu Darya | Karatag river | 38.363055 | 68.072059 | 22/07/2019 |
| 450. | Paracobitis_longicauda_345 | SWU22072019345 | FFU423-21 | MW649576 | Nemacheilidae | *Paracobitis longicauda* | Sheraliev, BM | Sheraliev, BM | Surxondaryo | Amu Darya | Karatag river | 38.363055 | 68.072059 | 22/07/2019 |
| 451. | Paracobitis_longicauda_346 | SWU22072019346 | FFU424-21 | MW649577 | Nemacheilidae | *Paracobitis longicauda* | Sheraliev, BM | Sheraliev, BM | Surxondaryo | Amu Darya | Karatag river | 38.363055 | 68.072059 | 22/07/2019 |
| 452. | Paracobitis_longicauda_347 | SWU22072019347 | FFU425-21 | MW649578 | Nemacheilidae | *Paracobitis longicauda* | Sheraliev, BM | Sheraliev, BM | Surxondaryo | Amu Darya | Karatag river | 38.363055 | 68.072059 | 22/07/2019 |
| 453. | Paracobitis_longicauda_348 | SWU22072019348 | FFU426-21 | MW649579 | Nemacheilidae | *Paracobitis longicauda* | Sheraliev, BM | Sheraliev, BM | Surxondaryo | Amu Darya | Karatag river | 38.363055 | 68.072059 | 22/07/2019 |
| 454. | Paracobitis_longicauda_349 | SWU22072019349 | FFU427-21 | MW649580 | Nemacheilidae | *Paracobitis longicauda* | Sheraliev, BM | Sheraliev, BM | Surxondaryo | Amu Darya | Karatag river | 38.363055 | 68.072059 | 22/07/2019 |
| 455. | Paracobitis_longicauda_350 | SWU22072019350 | FFU428-21 | MW649581 | Nemacheilidae | *Paracobitis longicauda* | Sheraliev, BM | Sheraliev, BM | Surxondaryo | Amu Darya | Karatag river | 38.363055 | 68.072059 | 22/07/2019 |
| 456. | Paracobitis_longicauda_351 | SWU22072019351 | FFU429-21 | MW649582 | Nemacheilidae | *Paracobitis longicauda* | Sheraliev, BM | Sheraliev, BM | Surxondaryo | Amu Darya | Karatag river | 38.363055 | 68.072059 | 22/07/2019 |
| 457. | Paracobitis_longicauda_352 | SWU22072019352 | FFU430-21 | MW649583 | Nemacheilidae | *Paracobitis longicauda* | Sheraliev, BM | Sheraliev, BM | Surxondaryo | Amu Darya | Karatag river | 38.363055 | 68.072059 | 22/07/2019 |
| 458. | Paracobitis_longicauda_375 | SWU23072019375 | FFU431-21 | MW649584 | Nemacheilidae | *Paracobitis longicauda* | Sheraliev, BM | Sheraliev, BM | Surxondaryo | Amu Darya | Tupalang River | 38.318966 | 68.006169 | 23/07/2019 |
| 459. | Paracobitis_longicauda_407 | SWU25072019407 | FFU432-21 | MW649585 | Nemacheilidae | *Paracobitis longicauda* | Sheraliev, BM | Sheraliev, BM | Surxondaryo | Amu Darya | Sherobod river | 37.747222 | 66.996666 | 25/07/2019 |
| 460. | Paracobitis_longicauda_408 | SWU25072019408 | FFU433-21 | MW649586 | Nemacheilidae | *Paracobitis longicauda* | Sheraliev, BM | Sheraliev, BM | Surxondaryo | Amu Darya | Sherobod river | 37.747222 | 66.996666 | 25/07/2019 |
| 461. | Paracobitis_longicauda_502 | SWU02082019502 | FFU434-21 | MW649587 | Nemacheilidae | *Paracobitis longicauda* | Sheraliev, BM | Sheraliev, BM | Samarkand | Zeravshan | Zeravshan | 39.676388 | 67.076111 | 02/08/2019 |
| 462. | Paracobitis_longicauda_683 | SWU27042019683 | FFU435-21 | MW649588 | Nemacheilidae | *Paracobitis longicauda* | Allayarov, S | Sheraliev, BM | Surxondaryo | Amu Darya | Karatag river | 38.363055 | 68.072059 | 27/04/2019 |
| 463. | Paracobitis_longicauda_684 | SWU27042019684 | FFU436-21 | MW649589 | Nemacheilidae | *Paracobitis longicauda* | Allayarov, S | Sheraliev, BM | Surxondaryo | Amu Darya | Karatag river | 38.363055 | 68.072059 | 27/04/2019 |
| 464. | Paracobitis_longicauda_685 | SWU27042019685 | FFU437-21 | MW649590 | Nemacheilidae | *Paracobitis longicauda* | Allayarov, S | Sheraliev, BM | Surxondaryo | Amu Darya | Karatag river | 38.363055 | 68.072059 | 27/04/2019 |
| 465. | Paracobitis_longicauda_686 | SWU27042019686 | FFU438-21 | MW649591 | Nemacheilidae | *Paracobitis longicauda* | Allayarov, S | Sheraliev, BM | Surxondaryo | Amu Darya | Karatag river | 38.363055 | 68.072059 | 27/04/2019 |
| 466. | Paracobitis_longicauda_687 | SWU27042019687 | FFU439-21 | MW649592 | Nemacheilidae | *Paracobitis longicauda* | Allayarov, S | Sheraliev, BM | Surxondaryo | Amu Darya | Karatag river | 38.363055 | 68.072059 | 27/04/2019 |
| 467. | Paracobitis_longicauda_688 | SWU27042019688 | FFU440-21 | MW649593 | Nemacheilidae | *Paracobitis longicauda* | Allayarov, S | Sheraliev, BM | Surxondaryo | Amu Darya | Karatag river | 38.363055 | 68.072059 | 27/04/2019 |
| 468. | Paracobitis_longicauda_689 | SWU27042019689 | FFU441-21 | MW649594 | Nemacheilidae | *Paracobitis longicauda* | Allayarov, S | Sheraliev, BM | Surxondaryo | Amu Darya | Karatag river | 38.363055 | 68.072059 | 27/04/2019 |
| 469. | Triplophysa_sp1_Zeravshan_503 | SWU02082019503 | FFU604-21 | MW649595 | Nemacheilidae | *Triplophysa* sp.1 | Sheraliev, BM | Sheraliev, BM | Samarkand | Zeravshan | Zeravshan | 39.676388 | 67.076111 | 02/08/2019 |
| 470. | Triplophysa_sp1_Zeravshan_513 | SWU02082019513 | FFU605-21 | MW649596 | Nemacheilidae | *Triplophysa* sp.1 | Sheraliev, BM | Sheraliev, BM | Samarkand | Zeravshan | Zeravshan | 39.676388 | 67.076111 | 02/08/2019 |
| 471. | Triplophysa_sp1_Zeravshan_514 | SWU02082019514 | FFU606-21 | MW649597 | Nemacheilidae | *Triplophysa* sp.1 | Sheraliev, BM | Sheraliev, BM | Samarkand | Zeravshan | Zeravshan | 39.676388 | 67.076111 | 02/08/2019 |
| 472. | Triplophysa_sp1_Zeravshan_515 | SWU02082019515 | FFU607-21 | MW649598 | Nemacheilidae | *Triplophysa* sp.1 | Sheraliev, BM | Sheraliev, BM | Samarkand | Zeravshan | Zeravshan | 39.676388 | 67.076111 | 02/08/2019 |
| 473. | Triplophysa_sp2_Chirchik_630 | SWU18082019630 | FFU608-21 | MW649599 | Nemacheilidae | *Triplophysa* sp.2 | Sheraliev, BM | Sheraliev, BM | Tashkent | Syr Darya | Chirchik river | 41.450555 | 69.598611 | 18/08/2019 |
| 474. | Triplophysa_sp2_Chirchik_631 | SWU18082019631 | FFU609-21 | MW649600 | Nemacheilidae | *Triplophysa* sp.2 | Sheraliev, BM | Sheraliev, BM | Tashkent | Syr Darya | Chirchik river | 41.450555 | 69.598611 | 18/08/2019 |
| 475. | Triplophysa_sp2_Chirchik_632 | SWU18082019632 | FFU610-21 | MW649601 | Nemacheilidae | *Triplophysa* sp.2 | Sheraliev, BM | Sheraliev, BM | Tashkent | Syr Darya | Chirchik river | 41.450555 | 69.598611 | 18/08/2019 |
| 476. | Triplophysa_sp2_Chirchik_633 | SWU18082019633 | FFU611-21 | MW649602 | Nemacheilidae | *Triplophysa* sp.2 | Sheraliev, BM | Sheraliev, BM | Tashkent | Syr Darya | Chirchik river | 41.450555 | 69.598611 | 18/08/2019 |
| 477. | Triplophysa_sp3_Fergana_647 | SWU13082019647 | FFU612-21 | MW649603 | Nemacheilidae | *Triplophysa ferganaensis* | Sheraliev, BM | Sheraliev, BM | Fergana | Syr Darya | Shohimardon River | 39.963088 | 71.759857 | 13/08/2019 |
| 478. | Triplophysa_sp3_Fergana_648 | SWU13082019648 | FFU613-21 | MW649604 | Nemacheilidae | *Triplophysa ferganaensis* | Sheraliev, BM | Sheraliev, BM | Fergana | Syr Darya | Shohimardon River | 39.963088 | 71.759857 | 13/08/2019 |
| 479. | Triplophysa_sp3_Fergana_649 | SWU13082019649 | FFU614-21 | MW649605 | Nemacheilidae | *Triplophysa ferganaensis* | Sheraliev, BM | Sheraliev, BM | Fergana | Syr Darya | Shohimardon River | 39.963088 | 71.759857 | 13/08/2019 |
| 480. | Triplophysa_sp3_Fergana_650 | SWU13082019650 | FFU615-21 | MW649606 | Nemacheilidae | *Triplophysa ferganaensis* | Sheraliev, BM | Sheraliev, BM | Fergana | Syr Darya | Shohimardon River | 39.963088 | 71.759857 | 13/08/2019 |
| 481. | Triplophysa_sp3_Fergana_651 | SWU13082019651 | FFU616-21 | MW649607 | Nemacheilidae | *Triplophysa ferganaensis* | Sheraliev, BM | Sheraliev, BM | Fergana | Syr Darya | Shohimardon River | 39.963088 | 71.759857 | 13/08/2019 |
| 482. | Triplophysa_sp3_Fergana_652 | SWU13082019652 | FFU617-21 | MW649608 | Nemacheilidae | *Triplophysa ferganaensis* | Sheraliev, BM | Sheraliev, BM | Fergana | Syr Darya | Shohimardon River | 39.963088 | 71.759857 | 13/08/2019 |
| 483. | Triplophysa_sp3_Fergana_653 | SWU13082019653 | FFU618-21 | MW649609 | Nemacheilidae | *Triplophysa ferganaensis* | Sheraliev, BM | Sheraliev, BM | Fergana | Syr Darya | Shohimardon River | 39.963088 | 71.759857 | 13/08/2019 |
| 484. | Triplophysa_sp3_Fergana_654 | SWU13082019654 | FFU619-21 | MW649610 | Nemacheilidae | *Triplophysa ferganaensis* | Sheraliev, BM | Sheraliev, BM | Fergana | Syr Darya | Shohimardon River | 39.963088 | 71.759857 | 13/08/2019 |
| 485. | Triplophysa_sp3_Fergana_655 | SWU13082019655 | FFU620-21 | MW649611 | Nemacheilidae | *Triplophysa ferganaensis* | Sheraliev, BM | Sheraliev, BM | Fergana | Syr Darya | Shohimardon River | 39.963088 | 71.759857 | 13/08/2019 |
| 486. | Triplophysa_sp3_Fergana_656 | SWU13082019656 | FFU621-21 | MW649612 | Nemacheilidae | *Triplophysa ferganaensis* | Sheraliev, BM | Sheraliev, BM | Fergana | Syr Darya | Shohimardon River | 39.963088 | 71.759857 | 13/08/2019 |
| 487. | Triplophysa_sp3_Fergana_657 | SWU13082019657 | FFU622-21 | MW649613 | Nemacheilidae | *Triplophysa ferganaensis* | Sheraliev, BM | Sheraliev, BM | Fergana | Syr Darya | Shohimardon River | 39.963088 | 71.759857 | 13/08/2019 |
| 488. | Triplophysa_sp3_Fergana_658 | SWU13082019658 | FFU623-21 | MW649614 | Nemacheilidae | *Triplophysa ferganaensis* | Sheraliev, BM | Sheraliev, BM | Fergana | Syr Darya | Shohimardon River | 39.963088 | 71.759857 | 13/08/2019 |
| 489. | Triplophysa_sp3_Fergana_659 | SWU13082019659 | FFU624-21 | MW649615 | Nemacheilidae | *Triplophysa ferganaensis* | Sheraliev, BM | Sheraliev, BM | Fergana | Syr Darya | Shohimardon River | 39.963088 | 71.759857 | 13/08/2019 |
| 490. | Triplophysa_sp3_Fergana_660 | SWU13082019660 | FFU625-21 | MW649616 | Nemacheilidae | *Triplophysa ferganaensis* | Sheraliev, BM | Sheraliev, BM | Fergana | Syr Darya | Shohimardon River | 39.963088 | 71.759857 | 13/08/2019 |
| 491. | Triplophysa_sp3_Fergana_661 | SWU13082019661 | FFU626-21 | MW649617 | Nemacheilidae | *Triplophysa ferganaensis* | Sheraliev, BM | Sheraliev, BM | Fergana | Syr Darya | Shohimardon River | 39.963088 | 71.759857 | 13/08/2019 |
| 492. | Triplophysa_sp3_Fergana_662 | SWU13082019662 | FFU627-21 | MW649618 | Nemacheilidae | *Triplophysa ferganaensis* | Sheraliev, BM | Sheraliev, BM | Fergana | Syr Darya | Shohimardon River | 39.963088 | 71.759857 | 13/08/2019 |
| 493. | Triplophysa_sp3_Fergana_663 | SWU13082019663 | FFU628-21 | MW649619 | Nemacheilidae | *Triplophysa ferganaensis* | Sheraliev, BM | Sheraliev, BM | Fergana | Syr Darya | Shohimardon River | 39.963088 | 71.759857 | 13/08/2019 |
| 494. | Triplophysa_sp3_Fergana_664 | SWU13082019664 | FFU629-21 | MW649620 | Nemacheilidae | *Triplophysa ferganaensis* | Sheraliev, BM | Sheraliev, BM | Fergana | Syr Darya | Shohimardon River | 39.963088 | 71.759857 | 13/08/2019 |
| 495. | Triplophysa_sp3_Fergana_665 | SWU13082019665 | FFU630-21 | MW649621 | Nemacheilidae | *Triplophysa ferganaensis* | Sheraliev, BM | Sheraliev, BM | Fergana | Syr Darya | Shohimardon River | 39.963088 | 71.759857 | 13/08/2019 |
| 496. | Triplophysa_sp3_Fergana_666 | SWU13082019666 | FFU631-21 | MW649622 | Nemacheilidae | *Triplophysa ferganaensis* | Sheraliev, BM | Sheraliev, BM | Fergana | Syr Darya | Shohimardon River | 39.963088 | 71.759857 | 13/08/2019 |
| 497. | Triplophysa_strauchii_052 | SWU07012017052 | FFU442-21 | MW649623 | Nemacheilidae | *Triplophysa strauchii* | Sheraliev, BM | Sheraliev, BM | Fergana | Syr Darya | Unnamed stream | 40.366437 | 71.316907 | 07/01/2017 |
| 498. | Triplophysa_strauchii_053 | SWU07012017053 | FFU443-21 | MW649624 | Nemacheilidae | *Triplophysa strauchii* | Sheraliev, BM | Sheraliev, BM | Fergana | Syr Darya | Unnamed stream | 40.366437 | 71.316907 | 07/01/2017 |
| 499. | Triplophysa_strauchii_056 | SWU07012017056 | FFU444-21 | MW649625 | Nemacheilidae | *Triplophysa strauchii* | Sheraliev, BM | Sheraliev, BM | Fergana | Syr Darya | Unnamed stream | 40.366437 | 71.316907 | 07/01/2017 |
| 500. | Triplophysa_strauchii_104 | SWU25012018104 | FFU445-21 | MW649626 | Nemacheilidae | *Triplophysa strauchii* | Sheraliev, BM | Sheraliev, BM | Fergana | Syr Darya | Fergana Canal | 40.388497 | 71.327960 | 25/01/2018 |
| 501. | Triplophysa_strauchii_105 | SWU25012018105 | FFU446-21 | MW649627 | Nemacheilidae | *Triplophysa strauchii* | Sheraliev, BM | Sheraliev, BM | Fergana | Syr Darya | Fergana Canal | 40.388497 | 71.327960 | 25/01/2018 |
| 502. | Triplophysa_strauchii_106 | SWU25012018106 | FFU447-21 | MW649628 | Nemacheilidae | *Triplophysa strauchii* | Sheraliev, BM | Sheraliev, BM | Fergana | Syr Darya | Fergana Canal | 40.388497 | 71.327960 | 25/01/2018 |
| 503. | Triplophysa_strauchii_129 | SWU21022018129 | FFU448-21 | MW649629 | Nemacheilidae | *Triplophysa strauchii* | Sheraliev, BM | Sheraliev, BM | Tashkent | Syr Darya | Chirchik River | 41.370266 | 69.512026 | 21/02/2018 |
| 504. | Triplophysa_strauchii_130 | SWU21022018130 | FFU449-21 | MW649630 | Nemacheilidae | *Triplophysa strauchii* | Sheraliev, BM | Sheraliev, BM | Tashkent | Syr Darya | Chirchik River | 41.370266 | 69.512026 | 21/02/2018 |
| 505. | Triplophysa_strauchii_204 | SWU11082018204 | FFU450-21 | MW649631 | Nemacheilidae | *Triplophysa strauchii* | Sheraliev, BM | Sheraliev, BM | Fergana | Syr Darya | Unnamed stream | 40.312769 | 71.811647 | 11/08/2018 |
| 506. | Triplophysa_strauchii_205 | SWU11082018205 | FFU451-21 | MW649632 | Nemacheilidae | *Triplophysa strauchii* | Sheraliev, BM | Sheraliev, BM | Fergana | Syr Darya | Unnamed stream | 40.312769 | 71.811647 | 11/08/2018 |
| 507. | Triplophysa_strauchii_206 | SWU11082018206 | FFU452-21 | MW649633 | Nemacheilidae | *Triplophysa strauchii* | Sheraliev, BM | Sheraliev, BM | Fergana | Syr Darya | Unnamed stream | 40.312769 | 71.811647 | 11/08/2018 |
| 508. | Triplophysa_strauchii_207 | SWU11082018207 | FFU453-21 | MW649634 | Nemacheilidae | *Triplophysa strauchii* | Sheraliev, BM | Sheraliev, BM | Fergana | Syr Darya | Unnamed stream | 40.312769 | 71.811647 | 11/08/2018 |
| 509. | Triplophysa_strauchii_208 | SWU11082018208 | FFU454-21 | MW649635 | Nemacheilidae | *Triplophysa strauchii* | Sheraliev, BM | Sheraliev, BM | Fergana | Syr Darya | Unnamed stream | 40.312769 | 71.811647 | 11/08/2018 |
| 510. | Triplophysa_strauchii_210 | SWU11082018210 | FFU455-21 | MW649636 | Nemacheilidae | *Triplophysa strauchii* | Sheraliev, BM | Sheraliev, BM | Fergana | Syr Darya | Unnamed stream | 40.312769 | 71.811647 | 11/08/2018 |
| 511. | Triplophysa_strauchii_230 | SWU15012017230 | FFU456-21 | MW649637 | Nemacheilidae | *Triplophysa strauchii* | Sheraliev, BM | Sheraliev, BM | Fergana | Syr Darya | Unnamed stream | 40.326408 | 71.820508 | 15/01/2017 |
| 512. | Triplophysa_strauchii_231 | SWU15012017231 | FFU457-21 | MW649638 | Nemacheilidae | *Triplophysa strauchii* | Sheraliev, BM | Sheraliev, BM | Fergana | Syr Darya | Unnamed stream | 40.326408 | 71.820508 | 15/01/2017 |
| 513. | Triplophysa_strauchii_248 | SWU11082018248 | FFU458-21 | MW649639 | Nemacheilidae | *Triplophysa strauchii* | Sheraliev, BM | Sheraliev, BM | Fergana | Syr Darya | Unnamed stream | 40.312769 | 71.811647 | 11/08/2018 |
| 514. | Triplophysa_strauchii_249 | SWU11082018249 | FFU459-21 | MW649640 | Nemacheilidae | *Triplophysa strauchii* | Sheraliev, BM | Sheraliev, BM | Fergana | Syr Darya | Unnamed stream | 40.312769 | 71.811647 | 11/08/2018 |
| 515. | Triplophysa_strauchii_551 | SWU09082019551 | FFU460-21 | MW649641 | Nemacheilidae | *Triplophysa strauchii* | Sheraliev, BM | Sheraliev, BM | Namangan | Syr Darya | Kara Darya | 40.918813 | 71.828896 | 09/08/2019 |
| 516. | Triplophysa_strauchii_617 | SWU18082019617 | FFU461-21 | MW649642 | Nemacheilidae | *Triplophysa strauchii* | Sheraliev, BM | Sheraliev, BM | Tashkent | Syr Darya | Chirchik River | 41.274444 | 69.389166 | 18/08/2019 |
| 517. | Triplophysa_strauchii_618 | SWU18082019618 | FFU462-21 | MW649643 | Nemacheilidae | *Triplophysa strauchii* | Sheraliev, BM | Sheraliev, BM | Tashkent | Syr Darya | Chirchik River | 41.274444 | 69.389166 | 18/08/2019 |
| 518. | Triplophysa_strauchii_619 | SWU18082019619 | FFU463-21 | MW649644 | Nemacheilidae | *Triplophysa strauchii* | Sheraliev, BM | Sheraliev, BM | Tashkent | Syr Darya | Chirchik River | 41.274444 | 69.389166 | 18/08/2019 |

| | | | | | | | | | | | | | | |
|---|---|---|---|---|---|---|---|---|---|---|---|---|---|---|
| 519. | Triplophysa_strauchii_621 | SWU18082019621 | FFU464-21 | MW649645 | Nemacheilidae | *Triplophysa strauchii* | Sheraliev, BM | Sheraliev, BM | Tashkent | Syr Darya | Chirchik River | 41.274444 | 69.389166 | 18/08/2019 |
| 520. | Triplophysa_strauchii_622 | SWU18082019622 | FFU465-21 | MW649646 | Nemacheilidae | *Triplophysa strauchii* | Sheraliev, BM | Sheraliev, BM | Tashkent | Syr Darya | Chirchik River | 41.274444 | 69.389166 | 18/08/2019 |
| 521. | Triplophysa_strauchii_623 | SWU18082019623 | FFU466-21 | MW649647 | Nemacheilidae | *Triplophysa strauchii* | Sheraliev, BM | Sheraliev, BM | Tashkent | Syr Darya | Chirchik River | 41.274444 | 69.389166 | 18/08/2019 |
| 522. | Triplophysa_strauchii_624 | SWU18082019624 | FFU467-21 | MW649648 | Nemacheilidae | *Triplophysa strauchii* | Sheraliev, BM | Sheraliev, BM | Tashkent | Syr Darya | Chirchik River | 41.274444 | 69.389166 | 18/08/2019 |
| 523. | Triplophysa_strauchii_625 | SWU18082019625 | FFU468-21 | MW649649 | Nemacheilidae | *Triplophysa strauchii* | Sheraliev, BM | Sheraliev, BM | Tashkent | Syr Darya | Chirchik River | 41.274444 | 69.389166 | 18/08/2019 |
| 524. | Triplophysa_strauchii_642 | SWU20082019642 | FFU469-21 | MW649650 | Nemacheilidae | *Triplophysa strauchii* | Sheraliev, BM | Sheraliev, BM | Fergana | Syr Darya | Unnamed stream | 40.305260 | 71.800688 | 20/08/2019 |
| 525. | Triplophysa_strauchii_643 | SWU20082019643 | FFU470-21 | MW649651 | Nemacheilidae | *Triplophysa strauchii* | Sheraliev, BM | Sheraliev, BM | Fergana | Syr Darya | Unnamed stream | 40.305260 | 71.800688 | 20/08/2019 |
| 526. | Sander_lucioperca_009 | SWU14082016009 | FFU471-21 | MW649652 | Percidae | *Sander lucioperca* | Sheraliev, BM | Sheraliev, BM | Surxondaryo | Amu Darya | South Surkhan R. | 37.854013 | 67.628373 | 14/08/2016 |
| 527. | Sander_lucioperca_010 | SWU14082016010 | FFU472-21 | MW649653 | Percidae | *Sander lucioperca* | Sheraliev, BM | Sheraliev, BM | Surxondaryo | Amu Darya | South Surkhan R. | 37.854013 | 67.628373 | 14/08/2016 |
| 528. | Sander_lucioperca_185 | SWU18072018185 | FFU473-21 | MW649654 | Percidae | *Sander lucioperca* | Sheraliev, BM | Sheraliev, BM | Navoiy | Zeravshan | Tudakul Lake | 39.801388 | 64.772222 | 18/07/2018 |
| 529. | Sander_lucioperca_186 | SWU18072018186 | FFU474-21 | MW649655 | Percidae | *Sander lucioperca* | Sheraliev, BM | Sheraliev, BM | Navoiy | Zeravshan | Tudakul Lake | 39.801388 | 64.772222 | 18/07/2018 |
| 530. | Sander_lucioperca_187 | SWU18072018187 | FFU475-21 | MW649656 | Percidae | *Sander lucioperca* | Sheraliev, BM | Sheraliev, BM | Navoiy | Zeravshan | Tudakul Lake | 39.801388 | 64.772222 | 18/07/2018 |
| 531. | Sander_lucioperca_188 | SWU18072018188 | FFU476-21 | MW649657 | Percidae | *Sander lucioperca* | Sheraliev, BM | Sheraliev, BM | Navoiy | Zeravshan | Tudakul Lake | 39.801388 | 64.772222 | 18/07/2018 |
| 532. | Sander_lucioperca_189 | SWU18072018189 | FFU477-21 | MW649658 | Percidae | *Sander lucioperca* | Sheraliev, BM | Sheraliev, BM | Navoiy | Zeravshan | Tudakul Lake | 39.801388 | 64.772222 | 18/07/2018 |
| 533. | Sander_lucioperca_439 | SWU27072019439 | FFU478-21 | MW649659 | Percidae | *Sander lucioperca* | Sheraliev, BM | Sheraliev, BM | Bukhara | Zeravshan | Echkiliksoy | 40.267551 | 64.743333 | 27/07/2019 |
| 534. | Sander_lucioperca_446 | SWU28072019446 | FFU479-21 | MW649660 | Percidae | *Sander lucioperca* | Sheraliev, BM | Sheraliev, BM | Bukhara | Zeravshan | Zeravshan | 40.071012 | 64.778429 | 28/07/2019 |
| 535. | Sander_lucioperca_477 | SWU30072019477 | FFU480-21 | MW649661 | Percidae | *Sander lucioperca* | Sheraliev, BM | Sheraliev, BM | Khorazm | Amu Darya | Unnamed canal | 41.282666 | 60.803916 | 30/07/2019 |
| 536. | Gambusia_holbrooki_140 | SWU25012018140 | FFU481-21 | MW649662 | Poeciliidae | *Gambusia holbrooki* | Sheraliev, BM | Sheraliev, BM | Fergana | Syr Darya | Fergana Canal | 40.396753 | 71.407328 | 25/01/2018 |
| 537. | Gambusia_holbrooki_141 | SWU25012018141 | FFU482-21 | MW649663 | Poeciliidae | *Gambusia holbrooki* | Sheraliev, BM | Sheraliev, BM | Fergana | Syr Darya | Fergana Canal | 40.396753 | 71.407328 | 25/01/2018 |
| 538. | Gambusia_holbrooki_142 | SWU25012018142 | FFU483-21 | MW649664 | Poeciliidae | *Gambusia holbrooki* | Sheraliev, BM | Sheraliev, BM | Fergana | Syr Darya | Fergana Canal | 40.396753 | 71.407328 | 25/01/2018 |
| 539. | Gambusia_holbrooki_143 | SWU25012018143 | FFU484-21 | MW649665 | Poeciliidae | *Gambusia holbrooki* | Sheraliev, BM | Sheraliev, BM | Fergana | Syr Darya | Fergana Canal | 40.396753 | 71.407328 | 25/01/2018 |
| 540. | Gambusia_holbrooki_216 | SWU16072018216 | FFU485-21 | MW649666 | Poeciliidae | *Gambusia holbrooki* | Sheraliev, BM | Sheraliev, BM | Bukhara | Zeravshan | Zeravshan | 40.146944 | 64.891666 | 16/07/2018 |
| 541. | Gambusia_holbrooki_217 | SWU16072018217 | FFU486-21 | MW649667 | Poeciliidae | *Gambusia holbrooki* | Sheraliev, BM | Sheraliev, BM | Bukhara | Zeravshan | Zeravshan | 40.146944 | 64.891666 | 16/07/2018 |
| 542. | Gambusia_holbrooki_262 | SWU19072019262 | FFU487-21 | MW649668 | Poeciliidae | *Gambusia holbrooki* | Sheraliev, BM | Sheraliev, BM | Surxondaryo | Amu Darya | Surkhan Darya | 37.255833 | 67.338611 | 19/07/2019 |
| 543. | Gambusia_holbrooki_263 | SWU19072019263 | FFU488-21 | MW649669 | Poeciliidae | *Gambusia holbrooki* | Sheraliev, BM | Sheraliev, BM | Surxondaryo | Amu Darya | Surkhan Darya | 37.255833 | 67.338611 | 19/07/2019 |
| 544. | Gambusia_holbrooki_296 | SWU21072019296 | FFU489-21 | MW649670 | Poeciliidae | *Gambusia holbrooki* | Sheraliev, BM | Sheraliev, BM | Surxondaryo | Amu Darya | Surkhan Darya | 37.238888 | 67.329722 | 21/07/2019 |
| 545. | Gambusia_holbrooki_297 | SWU21072019297 | FFU490-21 | MW649671 | Poeciliidae | *Gambusia holbrooki* | Sheraliev, BM | Sheraliev, BM | Surxondaryo | Amu Darya | Surkhan Darya | 37.238888 | 67.329722 | 21/07/2019 |
| 546. | Gambusia_holbrooki_298 | SWU21072019298 | FFU491-21 | MW649672 | Poeciliidae | *Gambusia holbrooki* | Sheraliev, BM | Sheraliev, BM | Surxondaryo | Amu Darya | Surkhan Darya | 37.238888 | 67.329722 | 21/07/2019 |
| 547. | Gambusia_holbrooki_299 | SWU21072019299 | FFU492-21 | MW649673 | Poeciliidae | *Gambusia holbrooki* | Sheraliev, BM | Sheraliev, BM | Surxondaryo | Amu Darya | Surkhan Darya | 37.238888 | 67.329722 | 21/07/2019 |
| 548. | Gambusia_holbrooki_334 | SWU22072019334 | FFU493-21 | MW649674 | Poeciliidae | *Gambusia holbrooki* | Sheraliev, BM | Sheraliev, BM | Surxondaryo | Amu Darya | Karatag river | 38.363055 | 68.072059 | 22/07/2019 |
| 549. | Gambusia_holbrooki_335 | SWU22072019335 | FFU494-21 | MW649675 | Poeciliidae | *Gambusia holbrooki* | Sheraliev, BM | Sheraliev, BM | Surxondaryo | Amu Darya | Karatag river | 38.363055 | 68.072059 | 22/07/2019 |
| 550. | Gambusia_holbrooki_336 | SWU22072019336 | FFU495-21 | MW649676 | Poeciliidae | *Gambusia holbrooki* | Sheraliev, BM | Sheraliev, BM | Surxondaryo | Amu Darya | Karatag river | 38.363055 | 68.072059 | 22/07/2019 |
| 551. | Gambusia_holbrooki_337 | SWU22072019337 | FFU496-21 | MW649677 | Poeciliidae | *Gambusia holbrooki* | Sheraliev, BM | Sheraliev, BM | Surxondaryo | Amu Darya | Karatag river | 38.363055 | 68.072059 | 22/07/2019 |
| 552. | Gambusia_holbrooki_393 | SWU24072019393 | FFU497-21 | MW649678 | Poeciliidae | *Gambusia holbrooki* | Sheraliev, BM | Sheraliev, BM | Surxondaryo | Amu Darya | Surkhan Darya | 37.823333 | 67.618611 | 24/07/2019 |
| 553. | Gambusia_holbrooki_394 | SWU24072019394 | FFU498-21 | MW649679 | Poeciliidae | *Gambusia holbrooki* | Sheraliev, BM | Sheraliev, BM | Surxondaryo | Amu Darya | Surkhan Darya | 37.823333 | 67.618611 | 24/07/2019 |
| 554. | Gambusia_holbrooki_395 | SWU24072019395 | FFU499-21 | MW649680 | Poeciliidae | *Gambusia holbrooki* | Sheraliev, BM | Sheraliev, BM | Surxondaryo | Amu Darya | Surkhan Darya | 37.823333 | 67.618611 | 24/07/2019 |
| 555. | Gambusia_holbrooki_396 | SWU24072019396 | FFU500-21 | MW649681 | Poeciliidae | *Gambusia holbrooki* | Sheraliev, BM | Sheraliev, BM | Surxondaryo | Amu Darya | Surkhan Darya | 37.823333 | 67.618611 | 24/07/2019 |
| 556. | Gambusia_holbrooki_421 | SWU27072019421 | FFU501-21 | MW649682 | Poeciliidae | *Gambusia holbrooki* | Sheraliev, BM | Sheraliev, BM | Bukhara | Zeravshan | Echkiliksoy | 40.267551 | 64.743333 | 27/07/2019 |
| 557. | Gambusia_holbrooki_422 | SWU27072019422 | FFU502-21 | MW649683 | Poeciliidae | *Gambusia holbrooki* | Sheraliev, BM | Sheraliev, BM | Bukhara | Zeravshan | Echkiliksoy | 40.267551 | 64.743333 | 27/07/2019 |
| 558. | Gambusia_holbrooki_423 | SWU27072019423 | FFU503-21 | MW649684 | Poeciliidae | *Gambusia holbrooki* | Sheraliev, BM | Sheraliev, BM | Bukhara | Zeravshan | Echkiliksoy | 40.267551 | 64.743333 | 27/07/2019 |
| 559. | Gambusia_holbrooki_424 | SWU27072019424 | FFU504-21 | MW649685 | Poeciliidae | *Gambusia holbrooki* | Sheraliev, BM | Sheraliev, BM | Bukhara | Zeravshan | Echkiliksoy | 40.267551 | 64.743333 | 27/07/2019 |
| 560. | Gambusia_holbrooki_425 | SWU27072019425 | FFU505-21 | MW649686 | Poeciliidae | *Gambusia holbrooki* | Sheraliev, BM | Sheraliev, BM | Bukhara | Zeravshan | Echkiliksoy | 40.267551 | 64.743333 | 27/07/2019 |
| 561. | Gambusia_holbrooki_463 | SWU28072019463 | FFU506-21 | MW649687 | Poeciliidae | *Gambusia holbrooki* | Sheraliev, BM | Sheraliev, BM | Bukhara | Zeravshan | Zeravshan | 40.071012 | 64.778429 | 28/07/2019 |
| 562. | Gambusia_holbrooki_464 | SWU28072019464 | FFU507-21 | MW649688 | Poeciliidae | *Gambusia holbrooki* | Sheraliev, BM | Sheraliev, BM | Bukhara | Zeravshan | Zeravshan | 40.071012 | 64.778429 | 28/07/2019 |
| 563. | Gambusia_holbrooki_525 | SWU02082019525 | FFU508-21 | MW649689 | Poeciliidae | *Gambusia holbrooki* | Sheraliev, BM | Sheraliev, BM | Samarkand | Zeravshan | Zeravshan | 39.676388 | 67.076111 | 02/08/2019 |
| 564. | Gambusia_holbrooki_526 | SWU02082019526 | FFU509-21 | MW649690 | Poeciliidae | *Gambusia holbrooki* | Sheraliev, BM | Sheraliev, BM | Samarkand | Zeravshan | Zeravshan | 39.676388 | 67.076111 | 02/08/2019 |
| 565. | Gambusia_holbrooki_527 | SWU02082019527 | FFU510-21 | MW649691 | Poeciliidae | *Gambusia holbrooki* | Sheraliev, BM | Sheraliev, BM | Samarkand | Zeravshan | Zeravshan | 39.676388 | 67.076111 | 02/08/2019 |
| 566. | Gambusia_holbrooki_528 | SWU02082019528 | FFU511-21 | MW649692 | Poeciliidae | *Gambusia holbrooki* | Sheraliev, BM | Sheraliev, BM | Samarkand | Zeravshan | Zeravshan | 39.676388 | 67.076111 | 02/08/2019 |
| 567. | Gambusia_holbrooki_529 | SWU02082019529 | FFU512-21 | MW649693 | Poeciliidae | *Gambusia holbrooki* | Sheraliev, BM | Sheraliev, BM | Samarkand | Zeravshan | Zeravshan | 39.676388 | 67.076111 | 02/08/2019 |
| 568. | Gambusia_holbrooki_530 | SWU02082019530 | FFU513-21 | MW649694 | Poeciliidae | *Gambusia holbrooki* | Sheraliev, BM | Sheraliev, BM | Samarkand | Zeravshan | Zeravshan | 39.676388 | 67.076111 | 02/08/2019 |
| 569. | Gambusia_holbrooki_559 | SWU09082019559 | FFU514-21 | MW649695 | Poeciliidae | *Gambusia holbrooki* | Sheraliev, BM | Sheraliev, BM | Namangan | Syr Darya | Kara Darya | 40.918813 | 71.828896 | 09/08/2019 |
| 570. | Gambusia_holbrooki_560 | SWU09082019560 | FFU515-21 | MW649696 | Poeciliidae | *Gambusia holbrooki* | Sheraliev, BM | Sheraliev, BM | Namangan | Syr Darya | Kara Darya | 40.918813 | 71.828896 | 09/08/2019 |
| 571. | Gambusia_holbrooki_561 | SWU09082019561 | FFU516-21 | MW649697 | Poeciliidae | *Gambusia holbrooki* | Sheraliev, BM | Sheraliev, BM | Namangan | Syr Darya | Kara Darya | 40.918813 | 71.828896 | 09/08/2019 |
| 572. | Gambusia_holbrooki_562 | SWU09082019562 | FFU517-21 | MW649698 | Poeciliidae | *Gambusia holbrooki* | Sheraliev, BM | Sheraliev, BM | Namangan | Syr Darya | Kara Darya | 40.918813 | 71.828896 | 09/08/2019 |
| 573. | Gambusia_holbrooki_598 | SWU18082019598 | FFU518-21 | MW649699 | Poeciliidae | *Gambusia holbrooki* | Sheraliev, BM | Sheraliev, BM | Tashkent | Syr Darya | Chirchik river | 41.274444 | 69.389166 | 18/08/2019 |
| 574. | Gambusia_holbrooki_599 | SWU18082019599 | FFU519-21 | MW649700 | Poeciliidae | *Gambusia holbrooki* | Sheraliev, BM | Sheraliev, BM | Tashkent | Syr Darya | Chirchik river | 41.274444 | 69.389166 | 18/08/2019 |
| 575. | Gambusia_holbrooki_600 | SWU18082019600 | FFU520-21 | MW649701 | Poeciliidae | *Gambusia holbrooki* | Sheraliev, BM | Sheraliev, BM | Tashkent | Syr Darya | Chirchik river | 41.274444 | 69.389166 | 18/08/2019 |
| 576. | Gambusia_holbrooki_601 | SWU18082019601 | FFU521-21 | MW649702 | Poeciliidae | *Gambusia holbrooki* | Sheraliev, BM | Sheraliev, BM | Tashkent | Syr Darya | Chirchik river | 41.274444 | 69.389166 | 18/08/2019 |
| 577. | Gambusia_holbrooki_675 | SWU21082019675 | FFU522-21 | MW649703 | Poeciliidae | *Gambusia holbrooki* | Rozimov, A | Sheraliev, BM | Khorazm | Amu Darya | Unnamed canal | 41.521638 | 60.566111 | 21/08/2019 |
| 578. | Gambusia_holbrooki_676 | SWU21082019676 | FFU523-21 | MW649704 | Poeciliidae | *Gambusia holbrooki* | Rozimov, A | Sheraliev, BM | Khorazm | Amu Darya | Unnamed canal | 41.521638 | 60.566111 | 21/08/2019 |
| 579. | Gambusia_holbrooki_677 | SWU21082019677 | FFU524-21 | MW649705 | Poeciliidae | *Gambusia holbrooki* | Rozimov, A | Sheraliev, BM | Khorazm | Amu Darya | Unnamed canal | 41.521638 | 60.566111 | 21/08/2019 |
| 580. | Oncorhynchus_mykiss_704 | SWU03012020704 | FFU525-21 | MW649706 | Salmonidae | *Oncorhynchus mykiss* | Xushboqov, Q | Sheraliev, BM | Surxondaryo | Amu Darya | Tupalang | 38.570627 | 67.805326 | 03/01/2020 |
| 581. | Oncorhynchus_mykiss_705 | SWU03012020705 | FFU526-21 | MW649707 | Salmonidae | *Oncorhynchus mykiss* | Xushboqov, Q | Sheraliev, BM | Surxondaryo | Amu Darya | Tupalang | 38.570627 | 67.805326 | 03/01/2020 |
| 582. | Silurus_glanis_050 | SWU31082016050 | FFU527-21 | MW649708 | Siluridae | *Silurus glanis* | Sheraliev, BM | Sheraliev, BM | Namangan | Syr Darya | Syr Darya | 40.882777 | 71.442222 | 31/08/2016 |
| 583. | Silurus_glanis_051 | SWU31082016051 | FFU528-21 | MW649709 | Siluridae | *Silurus glanis* | Sheraliev, BM | Sheraliev, BM | Namangan | Syr Darya | Syr Darya | 40.882777 | 71.442222 | 31/08/2016 |
| 584. | Silurus_glanis_176 | SWU18072018176 | FFU529-21 | MW649710 | Siluridae | *Silurus glanis* | Sheraliev, BM | Sheraliev, BM | Navoiy | Zeravshan | Tudakul Lake | 39.801388 | 64.772222 | 18/07/2018 |
| 585. | Silurus_glanis_281 | SWU20072019281 | FFU530-21 | MW649711 | Siluridae | *Silurus glanis* | Sheraliev, BM | Sheraliev, BM | Surxondaryo | Amu Darya | Amu Darya | 37.235555 | 67.660555 | 20/07/2019 |
| 586. | Silurus_glanis_282 | SWU20072019282 | FFU531-21 | MW649712 | Siluridae | *Silurus glanis* | Sheraliev, BM | Sheraliev, BM | Surxondaryo | Amu Darya | Amu Darya | 37.235555 | 67.660555 | 20/07/2019 |
| 587. | Silurus_glanis_283 | SWU20072019283 | FFU532-21 | MW649713 | Siluridae | *Silurus glanis* | Sheraliev, BM | Sheraliev, BM | Surxondaryo | Amu Darya | Amu Darya | 37.235555 | 67.660555 | 20/07/2019 |
| 588. | Silurus_glanis_667 | SWU21082019667 | FFU533-21 | MW649714 | Siluridae | *Silurus glanis* | Rozimov, A | Sheraliev, BM | Khorazm | Amu Darya | Amu Darya | 41.880305 | 60.450888 | 21/08/2019 |
| 589. | Silurus_glanis_668 | SWU21082019668 | FFU534-21 | MW649715 | Siluridae | *Silurus glanis* | Rozimov, A | Sheraliev, BM | Khorazm | Amu Darya | Amu Darya | 41.880305 | 60.450888 | 21/08/2019 |
| 590. | Glyptosternon_oschanini_576 | SWU09082019576 | FFU653-21 | MW649716 | Sisoridae | *Glyptosternon oschanini* | Sheraliev, BM | Sheraliev, BM | Namangan | Syr Darya | Kara Darya | 40.918813 | 71.828896 | 09/08/2019 |
| 591. | Glyptosternon_sp_084 | SWU08082016084 | FFU654-21 | MW649717 | Sisoridae | *Glyptosternon* sp. | Sheraliev, BM | Sheraliev, BM | Surxondaryo | Amu Darya | Tupalang river | 38.541702 | 67.823001 | 08/08/2016 |
| 592. | Glyptosternon_sp_085 | SWU08082016085 | FFU655-21 | MW649718 | Sisoridae | *Glyptosternon* sp. | Sheraliev, BM | Sheraliev, BM | Surxondaryo | Amu Darya | Tupalang river | 38.541702 | 67.823001 | 08/08/2016 |
| 593. | Glyptosternon_sp_087 | SWU08082016087 | FFU656-21 | MW649719 | Sisoridae | *Glyptosternon* sp. | Sheraliev, BM | Sheraliev, BM | Surxondaryo | Amu Darya | Tupalang river | 38.541702 | 67.823001 | 08/08/2016 |
| 594. | Glyptosternon_sp_088 | SWU08082016088 | FFU657-21 | MW649720 | Sisoridae | *Glyptosternon* sp. | Sheraliev, BM | Sheraliev, BM | Surxondaryo | Amu Darya | Tupalang river | 38.541702 | 67.823001 | 08/08/2016 |
| 595. | Glyptosternon_sp_218 | SWU08082016218 | FFU658-21 | MW649721 | Sisoridae | *Glyptosternon* sp. | Sheraliev, BM | Sheraliev, BM | Surxondaryo | Amu Darya | Tupalang river | 38.541702 | 67.823001 | 08/08/2016 |

| | | | | | | | | | | | | | | |
|---|---|---|---|---|---|---|---|---|---|---|---|---|---|---|
| 596. | Glyptosternon_sp_220 | SWU08082016220 | FFU659-21 | MW649722 | Sisoridae | *Glyptosternon* sp. | Sheraliev, BM | Sheraliev, BM | Surxondaryo | Amu Darya | Tupalang river | 38.541702 | 67.823001 | 08/08/2016 |
| 597. | Glyptosternon_sp_221 | SWU08082016221 | FFU660-21 | MW649723 | Sisoridae | *Glyptosternon* sp. | Sheraliev, BM | Sheraliev, BM | Surxondaryo | Amu Darya | Tupalang river | 38.541702 | 67.823001 | 08/08/2016 |
| 598. | Glyptosternon_sp_222 | SWU08082016222 | FFU661-21 | MW649724 | Sisoridae | *Glyptosternon* sp. | Sheraliev, BM | Sheraliev, BM | Surxondaryo | Amu Darya | Tupalang river | 38.541702 | 67.823001 | 08/08/2016 |
| 599. | Ctenopharyngodon_idella_107 | SWU28012018107 | FFU535-21 | MW649725 | Xenocyprididae | *Ctenopharyngodon idella* | Sheraliev, BM | Sheraliev, BM | Fergana | Syr Darya | Local fish market | | | 28/01/2018 |
| 600. | Ctenopharyngodon_idella_108 | SWU28012018108 | FFU536-21 | MW649726 | Xenocyprididae | *Ctenopharyngodon idella* | Sheraliev, BM | Sheraliev, BM | Fergana | Syr Darya | Local fish market | | | 28/01/2018 |
| 601. | Ctenopharyngodon_idella_109 | SWU28012018109 | FFU537-21 | MW649727 | Xenocyprididae | *Ctenopharyngodon idella* | Sheraliev, BM | Sheraliev, BM | Fergana | Syr Darya | Local fish market | | | 28/01/2018 |
| 602. | Ctenopharyngodon_idella_110 | SWU28012018110 | FFU538-21 | MW649728 | Xenocyprididae | *Ctenopharyngodon idella* | Sheraliev, BM | Sheraliev, BM | Fergana | Syr Darya | Local fish market | | | 28/01/2018 |
| 603. | Ctenopharyngodon_idella_111 | SWU28012018111 | FFU539-21 | MW649729 | Xenocyprididae | *Ctenopharyngodon idella* | Sheraliev, BM | Sheraliev, BM | Fergana | Syr Darya | Local fish market | | | 28/01/2018 |
| 604. | Ctenopharyngodon_idella_168 | SWU17072018168 | FFU540-21 | MW649730 | Xenocyprididae | *Ctenopharyngodon idella* | Sheraliev, BM | Sheraliev, BM | Bukhara | Zeravshan | Dzhalvan canal | 40.149666 | 64.885400 | 17/07/2018 |
| 605. | Ctenopharyngodon_idella_466 | SWU28072019466 | FFU541-21 | MW649731 | Xenocyprididae | *Ctenopharyngodon idella* | Sheraliev, BM | Sheraliev, BM | Bukhara | Zeravshan | Zeravshan | 40.071012 | 64.778429 | 28/07/2019 |
| 606. | Ctenopharyngodon_idella_672 | SWU21082019672 | FFU542-21 | MW649732 | Xenocyprididae | *Ctenopharyngodon idella* | Rozimov, A | Sheraliev, BM | Khorazm | Amu Darya | Amu Darya | 41.880305 | 60.450888 | 21/08/2019 |
| 607. | Hemiculter_leucisculus_019 | SWU16082016019 | FFU543-21 | MW649733 | Xenocyprididae | *Hemiculter leucisculus* | Sheraliev, BM | Sheraliev, BM | Qashqadaryo | Amu Darya | Qorasuv Stream | 38.570458 | 66.296166 | 16/08/2016 |
| 608. | Hemiculter_leucisculus_020 | SWU16082016020 | FFU544-21 | MW649734 | Xenocyprididae | *Hemiculter leucisculus* | Sheraliev, BM | Sheraliev, BM | Qashqadaryo | Amu Darya | Qorasuv Stream | 38.570458 | 66.296166 | 16/08/2016 |
| 609. | Hemiculter_leucisculus_126 | SWU08022018126 | FFU545-21 | MW649735 | Xenocyprididae | *Hemiculter leucisculus* | Sheraliev, BM | Sheraliev, BM | Fergana | Syr Darya | Sokh River | 40.340176 | 71.013559 | 08/02/2018 |
| 610. | Hemiculter_leucisculus_151 | SWU16072018151 | FFU546-21 | MW649736 | Xenocyprididae | *Hemiculter leucisculus* | Sheraliev, BM | Sheraliev, BM | Bukhara | Zeravshan | Zeravshan | 40.146944 | 64.891666 | 16/07/2018 |
| 611. | Hemiculter_leucisculus_152 | SWU16072018152 | FFU547-21 | MW649737 | Xenocyprididae | *Hemiculter leucisculus* | Sheraliev, BM | Sheraliev, BM | Bukhara | Zeravshan | Zeravshan | 40.146944 | 64.891666 | 16/07/2018 |
| 612. | Hemiculter_leucisculus_166 | SWU17072018166 | FFU548-21 | MW649738 | Xenocyprididae | *Hemiculter leucisculus* | Sheraliev, BM | Sheraliev, BM | Bukhara | Zeravshan | Dzhalvan canal | 40.149666 | 64.885400 | 17/07/2018 |
| 613. | Hemiculter_leucisculus_167 | SWU17072018167 | FFU549-21 | MW649739 | Xenocyprididae | *Hemiculter leucisculus* | Sheraliev, BM | Sheraliev, BM | Bukhara | Zeravshan | Dzhalvan canal | 40.149666 | 64.885400 | 17/07/2018 |
| 614. | Hemiculter_leucisculus_233 | SWU16082016233 | FFU550-21 | MW649740 | Xenocyprididae | *Hemiculter leucisculus* | Sheraliev, BM | Sheraliev, BM | Qashqadaryo | Amu Darya | Qorasuv Stream | 38.570458 | 66.296166 | 16/08/2016 |
| 615. | Hemiculter_leucisculus_318 | SWU21072019318 | FFU551-21 | MW649741 | Xenocyprididae | *Hemiculter leucisculus* | Sheraliev, BM | Sheraliev, BM | Surxondaryo | Amu Darya | Surkhan Darya | 37.238888 | 67.329722 | 21/07/2019 |
| 616. | Hemiculter_leucisculus_319 | SWU21072019319 | FFU552-21 | MW649742 | Xenocyprididae | *Hemiculter leucisculus* | Sheraliev, BM | Sheraliev, BM | Surxondaryo | Amu Darya | Surkhan Darya | 37.238888 | 67.329722 | 21/07/2019 |
| 617. | Hemiculter_leucisculus_324 | SWU21072019324 | FFU553-21 | MW649743 | Xenocyprididae | *Hemiculter leucisculus* | Sheraliev, BM | Sheraliev, BM | Surxondaryo | Amu Darya | Surkhan Darya | 37.238888 | 67.329722 | 21/07/2019 |
| 618. | Hemiculter_leucisculus_391 | SWU24072019391 | FFU554-21 | MW649744 | Xenocyprididae | *Hemiculter leucisculus* | Sheraliev, BM | Sheraliev, BM | Surxondaryo | Amu Darya | Surkhan Darya | 37.823333 | 67.618611 | 24/07/2019 |
| 619. | Hemiculter_leucisculus_392 | SWU24072019392 | FFU555-21 | MW649745 | Xenocyprididae | *Hemiculter leucisculus* | Sheraliev, BM | Sheraliev, BM | Surxondaryo | Amu Darya | Surkhan Darya | 37.823333 | 67.618611 | 24/07/2019 |
| 620. | Hemiculter_leucisculus_401 | SWU24072019401 | FFU556-21 | MW649746 | Xenocyprididae | *Hemiculter leucisculus* | Sheraliev, BM | Sheraliev, BM | Surxondaryo | Amu Darya | Surkhan Darya | 37.823333 | 67.618611 | 24/07/2019 |
| 621. | Hemiculter_leucisculus_453 | SWU28072019453 | FFU557-21 | MW649747 | Xenocyprididae | *Hemiculter leucisculus* | Sheraliev, BM | Sheraliev, BM | Bukhara | Zeravshan | Zeravshan | 40.071012 | 64.778429 | 28/07/2019 |
| 622. | Hemiculter_leucisculus_454 | SWU28072019454 | FFU558-21 | MW649748 | Xenocyprididae | *Hemiculter leucisculus* | Sheraliev, BM | Sheraliev, BM | Bukhara | Zeravshan | Zeravshan | 40.071012 | 64.778429 | 28/07/2019 |
| 623. | Hemiculter_leucisculus_455 | SWU28072019455 | FFU559-21 | MW649749 | Xenocyprididae | *Hemiculter leucisculus* | Sheraliev, BM | Sheraliev, BM | Bukhara | Zeravshan | Zeravshan | 40.071012 | 64.778429 | 28/07/2019 |
| 624. | Hemiculter_leucisculus_456 | SWU28072019456 | FFU560-21 | MW649750 | Xenocyprididae | *Hemiculter leucisculus* | Sheraliev, BM | Sheraliev, BM | Bukhara | Zeravshan | Zeravshan | 40.071012 | 64.778429 | 28/07/2019 |
| 625. | Hemiculter_leucisculus_469 | SWU29072019469 | FFU561-21 | MW649751 | Xenocyprididae | *Hemiculter leucisculus* | Sheraliev, BM | Sheraliev, BM | Bukhara | Zeravshan | Unnamed stream | 40.147222 | 64.905805 | 29/07/2019 |
| 626. | Hemiculter_leucisculus_470 | SWU29072019470 | FFU562-21 | MW649752 | Xenocyprididae | *Hemiculter leucisculus* | Sheraliev, BM | Sheraliev, BM | Bukhara | Zeravshan | Unnamed stream | 40.147222 | 64.905805 | 29/07/2019 |
| 627. | Hemiculter_leucisculus_478 | SWU30072019478 | FFU563-21 | MW649753 | Xenocyprididae | *Hemiculter leucisculus* | Sheraliev, BM | Sheraliev, BM | Khorazm | Amu Darya | Unnamed canal | 41.282666 | 60.803916 | 30/07/2019 |
| 628. | Hemiculter_leucisculus_479 | SWU30072019479 | FFU564-21 | MW649754 | Xenocyprididae | *Hemiculter leucisculus* | Sheraliev, BM | Sheraliev, BM | Khorazm | Amu Darya | Unnamed canal | 41.282666 | 60.803916 | 30/07/2019 |
| 629. | Hemiculter_leucisculus_480 | SWU30072019480 | FFU565-21 | MW649755 | Xenocyprididae | *Hemiculter leucisculus* | Sheraliev, BM | Sheraliev, BM | Khorazm | Amu Darya | Unnamed canal | 41.282666 | 60.803916 | 30/07/2019 |
| 630. | Hemiculter_leucisculus_696 | SWU27042019696 | FFU566-21 | MW649756 | Xenocyprididae | *Hemiculter leucisculus* | Allayarov, S | Sheraliev, BM | Surxondaryo | Amu Darya | Karatag river | 38.363055 | 68.072059 | 27/04/2019 |
| 631. | Hemiculter_leucisculus_711 | SWU19042020711 | FFU567-21 | MW649757 | Xenocyprididae | *Hemiculter leucisculus* | Rozimov, A | Sheraliev, BM | Khorazm | Amu Darya | Unnamed stream | 41.540222 | 60.166666 | 19/04/2020 |
| 632. | Hypophthalmichthys_molitrix_089 | SWU23012018089 | FFU568-21 | MW649758 | Xenocyprididae | *Hypophthalmichthys molitrix* | Sheraliev, BM | Sheraliev, BM | Fergana | Syr Darya | Local fish market | | | 23/01/2018 |
| 633. | Hypophthalmichthys_molitrix_090 | SWU23012018090 | FFU569-21 | MW649759 | Xenocyprididae | *Hypophthalmichthys molitrix* | Sheraliev, BM | Sheraliev, BM | Fergana | Syr Darya | Local fish market | | | 23/01/2018 |
| 634. | Hypophthalmichthys_molitrix_091 | SWU23012018091 | FFU570-21 | MW649760 | Xenocyprididae | *Hypophthalmichthys molitrix* | Sheraliev, BM | Sheraliev, BM | Fergana | Syr Darya | Local fish market | | | 23/01/2018 |
| 635. | Hypophthalmichthys_molitrix_097 | SWU25012018097 | FFU571-21 | MW649761 | Xenocyprididae | *Hypophthalmichthys molitrix* | Sheraliev, BM | Sheraliev, BM | Fergana | Syr Darya | Fergana Canal | 40.388497 | 71.327960 | 25/01/2018 |
| 636. | Hypophthalmichthys_molitrix_182 | SWU18072018182 | FFU572-21 | MW649762 | Xenocyprididae | *Hypophthalmichthys molitrix* | Sheraliev, BM | Sheraliev, BM | Navoiy | Zeravshan | Tudakul Lake | 39.801388 | 64.772222 | 18/07/2018 |
| 637. | Hypophthalmichthys_molitrix_450 | SWU28072019450 | FFU573-21 | MW649763 | Xenocyprididae | *Hypophthalmichthys molitrix* | Sheraliev, BM | Sheraliev, BM | Bukhara | Zeravshan | Zeravshan | 40.071012 | 64.778429 | 28/07/2019 |
| 638. | Hypophthalmichthys_molitrix_451 | SWU28072019451 | FFU574-21 | MW649764 | Xenocyprididae | *Hypophthalmichthys molitrix* | Sheraliev, BM | Sheraliev, BM | Bukhara | Zeravshan | Zeravshan | 40.071012 | 64.778429 | 28/07/2019 |
| 639. | Hypophthalmichthys_molitrix_452 | SWU28072019452 | FFU575-21 | MW649765 | Xenocyprididae | *Hypophthalmichthys molitrix* | Sheraliev, BM | Sheraliev, BM | Bukhara | Zeravshan | Zeravshan | 40.071012 | 64.778429 | 28/07/2019 |
| 640. | Hypophthalmichthys_nobilis_112 | SWU29012018112 | FFU576-21 | MW649766 | Xenocyprididae | *Hypophthalmichthys nobilis* | Sheraliev, BM | Sheraliev, BM | Fergana | Syr Darya | Local fish market | | | 29/01/2018 |
| 641. | Hypophthalmichthys_nobilis_113 | SWU29012018113 | FFU577-21 | MW649767 | Xenocyprididae | *Hypophthalmichthys nobilis* | Sheraliev, BM | Sheraliev, BM | Fergana | Syr Darya | Local fish market | | | 29/01/2018 |
| 642. | Hypophthalmichthys_nobilis_114 | SWU29012018114 | FFU578-21 | MW649768 | Xenocyprididae | *Hypophthalmichthys nobilis* | Sheraliev, BM | Sheraliev, BM | Fergana | Syr Darya | Local fish market | | | 29/01/2018 |
| 643. | Hypophthalmichthys_nobilis_115 | SWU29012018115 | FFU579-21 | MW649769 | Xenocyprididae | *Hypophthalmichthys nobilis* | Sheraliev, BM | Sheraliev, BM | Fergana | Syr Darya | Local fish market | | | 29/01/2018 |
| 644. | Hypophthalmichthys_nobilis_116 | SWU29012018116 | FFU580-21 | MW649770 | Xenocyprididae | *Hypophthalmichthys nobilis* | Sheraliev, BM | Sheraliev, BM | Fergana | Syr Darya | Local fish market | | | 29/01/2018 |
| 645. | Mylopharyngodon_piceus_714 | SWU29052020714 | FFU581-21 | MW649771 | Xenocyprididae | *Mylopharyngodon piceus* | Rozimov, A | Sheraliev, BM | Khorazm | Amu Darya | Amu Darya | 41.550111 | 60.868916 | 29/05/2020 |
| 646. | Mylopharyngodon_piceus_715 | SWU06062020715 | FFU582-21 | MW649772 | Xenocyprididae | *Mylopharyngodon piceus* | Rozimov, A | Sheraliev, BM | Khorazm | Amu Darya | Amu Darya | 41.550111 | 60.868916 | 06/06/2020 |
| 647. | Opsariichthys_cf_bidens_027 | SWU31082016027 | FFU583-21 | MW649773 | Xenocyprididae | *Opsariichthys bidens* | Sheraliev, BM | Sheraliev, BM | Namangan | Syr Darya | Syr Darya | 40.882777 | 71.442222 | 31/08/2016 |
| 648. | Opsariichthys_cf_bidens_028 | SWU31082016028 | FFU584-21 | MW649774 | Xenocyprididae | *Opsariichthys bidens* | Sheraliev, BM | Sheraliev, BM | Namangan | Syr Darya | Syr Darya | 40.882777 | 71.442222 | 31/08/2016 |
| 649. | Opsariichthys_cf_bidens_029 | SWU31082016029 | FFU585-21 | MW649775 | Xenocyprididae | *Opsariichthys bidens* | Sheraliev, BM | Sheraliev, BM | Namangan | Syr Darya | Syr Darya | 40.882777 | 71.442222 | 31/08/2016 |
| 650. | Opsariichthys_cf_bidens_030 | SWU31082016030 | FFU586-21 | MW649776 | Xenocyprididae | *Opsariichthys bidens* | Sheraliev, BM | Sheraliev, BM | Namangan | Syr Darya | Syr Darya | 40.882777 | 71.442222 | 31/08/2016 |
| 651. | Opsariichthys_cf_bidens_032 | SWU31082016032 | FFU587-21 | MW649777 | Xenocyprididae | *Opsariichthys bidens* | Sheraliev, BM | Sheraliev, BM | Namangan | Syr Darya | Syr Darya | 40.882777 | 71.442222 | 31/08/2016 |
| 652. | Opsariichthys_cf_bidens_389 | SWU24072019389 | FFU588-21 | MW649778 | Xenocyprididae | *Opsariichthys bidens* | Sheraliev, BM | Sheraliev, BM | Surxondaryo | Amu Darya | Surkhan Darya | 37.823333 | 67.618611 | 24/07/2019 |
| 653. | Opsariichthys_cf_bidens_608 | SWU18082019608 | FFU589-21 | MW649779 | Xenocyprididae | *Opsariichthys bidens* | Sheraliev, BM | Sheraliev, BM | Tashkent | Syr Darya | Chirchik River | 41.274444 | 69.389166 | 18/08/2019 |
| 654. | Parabramis_pekinensis_131 | SWU20022018131 | FFU590-21 | MW649780 | Xenocyprididae | *Parabramis pekinensis* | Sheraliev, BM | Sheraliev, BM | Navoiy | Zeravshan | Tudakul Lake | 39.907075 | 64.860898 | 20/02/2018 |
| 655. | Parabramis_pekinensis_132 | SWU20022018132 | FFU591-21 | MW649781 | Xenocyprididae | *Parabramis pekinensis* | Sheraliev, BM | Sheraliev, BM | Navoiy | Zeravshan | Tudakul Lake | 39.907075 | 64.860898 | 20/02/2018 |
| 656. | Parabramis_pekinensis_150 | SWU16072018150 | FFU592-21 | MW649782 | Xenocyprididae | *Parabramis pekinensis* | Sheraliev, BM | Sheraliev, BM | Bukhara | Zeravshan | Zeravshan | 40.146944 | 64.891666 | 16/07/2018 |
| 657. | Parabramis_pekinensis_165 | SWU17072018165 | FFU593-21 | MW649783 | Xenocyprididae | *Parabramis pekinensis* | Sheraliev, BM | Sheraliev, BM | Bukhara | Zeravshan | Dzhalvan canal | 40.149666 | 64.885400 | 17/07/2018 |
| 658. | Parabramis_pekinensis_173 | SWU18082018173 | FFU594-21 | MW649784 | Xenocyprididae | *Parabramis pekinensis* | Sheraliev, BM | Sheraliev, BM | Navoiy | Zeravshan | Tudakul Lake | 39.801388 | 64.772222 | 18/07/2018 |
| 659. | Parabramis_pekinensis_174 | SWU18082018174 | FFU595-21 | MW649785 | Xenocyprididae | *Parabramis pekinensis* | Sheraliev, BM | Sheraliev, BM | Navoiy | Zeravshan | Tudakul Lake | 39.801388 | 64.772222 | 18/07/2018 |
| 660. | Parabramis_pekinensis_175 | SWU18082018175 | FFU596-21 | MW649786 | Xenocyprididae | *Parabramis pekinensis* | Sheraliev, BM | Sheraliev, BM | Navoiy | Zeravshan | Tudakul Lake | 39.801388 | 64.772222 | 18/07/2018 |
| 661. | Parabramis_pekinensis_178 | SWU18082018178 | FFU597-21 | MW649787 | Xenocyprididae | *Parabramis pekinensis* | Sheraliev, BM | Sheraliev, BM | Navoiy | Zeravshan | Tudakul Lake | 39.801388 | 64.772222 | 18/07/2018 |
| 662. | Parabramis_pekinensis_179 | SWU18082018179 | FFU598-21 | MW649788 | Xenocyprididae | *Parabramis pekinensis* | Sheraliev, BM | Sheraliev, BM | Navoiy | Zeravshan | Tudakul Lake | 39.801388 | 64.772222 | 18/07/2018 |
| 663. | Parabramis_pekinensis_457 | SWU28072019457 | FFU599-21 | MW649789 | Xenocyprididae | *Parabramis pekinensis* | Sheraliev, BM | Sheraliev, BM | Bukhara | Zeravshan | Zeravshan | 40.071012 | 64.778429 | 28/07/2019 |
| 664. | Parabramis_pekinensis_458 | SWU28072019458 | FFU600-21 | MW649790 | Xenocyprididae | *Parabramis pekinensis* | Sheraliev, BM | Sheraliev, BM | Bukhara | Zeravshan | Zeravshan | 40.071012 | 64.778429 | 28/07/2019 |
| 665. | Parabramis_pekinensis_481 | SWU30072019481 | FFU601-21 | MW649791 | Xenocyprididae | *Parabramis pekinensis* | Sheraliev, BM | Sheraliev, BM | Khorazm | Amu Darya | Unnamed canal | 41.282666 | 60.803916 | 30/07/2019 |
| 666. | Parabramis_pekinensis_482 | SWU30072019482 | FFU602-21 | MW649792 | Xenocyprididae | *Parabramis pekinensis* | Sheraliev, BM | Sheraliev, BM | Khorazm | Amu Darya | Unnamed canal | 41.282666 | 60.803916 | 30/07/2019 |

**TABLE S2.** Fish species identification from GenBank and BOLD databases

| No | Species name | GenBank | | BOLD | |
|---|---|---|---|---|---|
| | | Species name | Identity (%) | Species name | Identity (%) |
| 1. | *Rhodeus ocellatus* | *Rhodeus ocellatus* | 100 | *Rhodeus ocellatus* | 100 |
| 2. | *Rhodeus* sp. | ***Rhodeus ocellatus*** | **92.44** | **No match** | **0** |
| 3. | *Acipenser baerii* | *Acipenser baerii* | 99.71-99.85 | *Acipenser baerii* | 100 |
| 4. | *Pseudoscaphirhynchus hermanni* | ***Pseudoscaphirhynchus kaufmanni*** | **99.71** | ***Pseudoscaphirhynchus kaufmanni*** | **99.85-100** |
| 5. | *Pseudoscaphirhynchus kaufmanni* | *Pseudoscaphirhynchus kaufmanni* | 100 | *Pseudoscaphirhynchus kaufmanni* | 99.85-100 |
| 6. | *Channa argus* | *Channa argus* | 100 | *Channa argus* | 100 |
| 7. | *Sabanejewia aurata* | *Sabanejewia aurata* | 98.58-98.74 | *Sabanejewia aurata* | 98.77-99.23 |
| 8. | *Cottus spinulosus* | ***Cottus ricei*** | **98.47** | ***Cottus ricei*** | **98.48** |
| 9. | *Capoeta heratensis* | *Capoeta capoeta heratensis* | 100 | *Capoeta heratensis* | 100 |
| 10. | *Carassius auratus* | *Carassius auratus* | 100 | *Carassius auratus* | 100 |
| 11. | *Carassius gibelio* | *Carassius gibelio* | 100 | *Carassius gibelio* | 100 |
| 12. | *Cyprinus carpio* | *Cyprinus carpio* | 100 | *Cyprinus carpio* | 100 |
| 13. | *Luciobarbus brachycephalus* | *Luciobarbus brachycephalus* | 99.70 | *Luciobarbus brachycephalus* | 99.54-99.69 |
| 14. | *Luciobarbus conocephalus* | ***Luciobarbus capito*** | **98.83-100** | *Luciobarbus conocephalus* | 99.85-100 |
| 15. | *Schizothorax eurystomus* | *Schizothorax eurystomus* | 100 | *Schizothorax eurystomus* | 100 |
| 16. | *Schizothorax fedtschenkoi* | ***Schizothorax davidi*** | **95.29** | **No match** | **0** |
| 17. | *Schizothorax* sp. | ***Schizothorax biddulphi*** | **95.31** | **No match** | **0** |
| 18. | *Esox lucius* | *Esox lucius* | 99.55 | *Esox lucius* | 99.55 |
| 19. | *Neogobius melanostomus* | *Neogobius melanostomus* | 100 | *Neogobius melanostomus* | 100 |
| 20. | *Neogobius pallasi* | ***Neogobius fluviatilis*** | **93.42** | *Neogobius pallasi* | 99.38 |
| 21. | *Rhinogobius* sp. | ***Rhinogobius sp*** | **97.18** | *Rhinogobius sp.* | 100 |
| 22. | *Abbottina rivularis* | *Abbottina rivularis* | 99.71 | *Abbottina rivularis* | 99.71 |
| 23. | *Gobio lepidolaemus* | *Gobio lepidolaemus* | 100 | *Gobio lepidolaemus* | 100 |
| 24. | *Gobio nigrescens* | *Gobio nigrescens* | 100 | *Gobio nigrescens* | 99.54 |
| 25. | *Gobio sibiricus* | *Gobio sibiricus* | 99.54-100 | *Gobio sibiricus* | 99.54 |
| 26. | *Pseudorasbora parva* | *Pseudorasbora parva* | 100 | *Pseudorasbora parva* | 100 |
| 27. | *Abramis brama* | *Abramis brama* | 100 | *Abramis brama* | 100 |
| 28. | *Alburnoides holciki* | *Alburnoides holciki* | 100 | *Alburnoides holciki* | 100 |
| 29. | *Alburnus taeniatus* | ***Alburnus escherichii*** | **98.82** | ***Alburnus escherichii*** | **98.92** |
| 30. | *Alburnus chalcoides* | *Alburnus chalcoides* | 99.85 | *Alburnus chalcoides* | 100 |
| 31. | *Alburnus oblongus* | ***Alburnus escherichii*** | **98.39** | ***Alburnus escherichii*** | **98.62** |
| 32. | *Capoetobrama kuschakewitschi* | ***Vimba melanops*** | **96.47** | **No match** | **0** |
| 33. | *Leuciscus aspius* | *Leuciscus aspius* | 99.70 | *Leuciscus aspius* | 99.85 |
| 34. | *Leuciscus lehmanni* | ***Leuciscus baicalensis*** | **99.85** | ***Leuciscus baicalensis*** | **100** |
| 35. | *Leuciscus squaliusculus* | ***Leuciscus baicalensis*** | **99.71** | ***Leuciscus baicalensis*** | **99.85** |
| 36. | *Pelecus cultratus* | *Pelecus cultratus* | 100 | *Pelecus cultratus* | 100 |
| 37. | *Rutilus lacustris* | *Rutilus lacustris* | 100 | *Rutilus lacustris* | 100 |
| 38. | *Dzihunia amudarjensis* | ***Schistura longa*** | **86.66** | **No match** | **0** |
| 39. | *Dzihunia* sp.1 | ***Schistura reticulofasciata*** | **87.85** | **No match** | **0** |
| 40. | *Dzihunia* sp. 2 | ***Schistura longa*** | **87.98** | **No match** | **0** |
| 41. | *Dzihunia* sp. 3 | ***Schistura longa*** | **88.42** | **No match** | **0** |
| 42. | *Paracobitis longicauda* | ***Paracobitis persa*** | **94.02** | **No match** | **0** |

| | | | | | |
|---|---|---|---|---|---|
| 43. | *Triplopysa ferganaensis* | ***Triplophysa orientalis*** | **94.69** | **No match** | **0** |
| 44. | *Triplophysa strauchii* | *Triplophysa strauchii* | 99.27 | *Triplophysa strauchii* | 99.12 |
| 45. | *Triplohysa* sp.1 | ***Triplophysa aliensis*** | **98.37** | ***Triplophysa aliensis*** | **98.36** |
| 46. | *Triplopysa* sp. 2 | ***Triplophysa scleroptera*** | **95.43** | **No match** | **0** |
| 47. | *Sander lucioperca* | *Sander lucioperca* | 99.85 | *Sander lucioperca* | 100 |
| 48. | *Gambusia holbrooki* | *Gambusia holbrooki* | 99.85-100 | *Gambusia holbrooki* | 100 |
| 49. | *Oncorhynchus mykiss* | *Oncorhynchus mykiss* | 99.85 | *Oncorhynchus mykiss* | 100 |
| 50. | *Silurus glanis* | *Silurus glanis* | 99.27-99.67 | *Silurus glanis* | 99.69 |
| 51. | *Glyptosternon oschanini* | *Glyptosternon oschanini* | 99.84-100 | **No match** | **0** |
| 52. | *Glyptosternon* sp. | ***Glyptosternon oschanini*** | **97.74-97.90** | **No match** | **0** |
| 53. | *Ctenopharyngodon idella* | *Ctenopharyngodon idella* | 100 | *Ctenopharyngodon idella* | 100 |
| 54. | *Hemiculter leuciscuslus* | *Hemiculter leuciscuslus* | 99.85 | *Hemiculter leucisculus* | 99.85 |
| 55. | *Hypophthalmichthys molitrix* | *Hypophthalmichthys molitrix* | 99.56-100 | *Hypophthalmichthys molitrix* | 100 |
| 56. | *Hypophthalmichthys nobilis* | *Hypophthalmichthys nobilis* | 100 | *Hypophthalmichthys nobilis* | 100 |
| 57. | *Mylopharyngodon piceus* | *Mylopharyngodon piceus* | 99.85 | *Mylopharyngodon piceus* | 100 |
| 58. | *Opsariichthys bidens* | *Opsariichthys bidens* | 98.97-100 | *Opsariichthys bidens* | 100 |
| 59. | *Parabramis pekinensis* | *Parabramis pekinensis* | 99.85 | *Parabramis pekinensis* | 100 |

**TABLE S3.** Barcode Index Number (BIN), average and maximum intraspecific distance and distance to nearest neighbor (NN).

| No. | Species name | BINs | Average intraspecific distance (%) | Maximum intraspecific distance (%) | Nearest neighbor (Process ID) | Distance to nearest neighbor (%) |
|---|---|---|---|---|---|---|
| 1. | *Rhodeus ocellatus* | BOLD:ADC7506 | 0.08 | 0.15 | *Rhodeus* sp. (FFU022-20) | 9.78 |
| 2. | *Rhodeus* sp. | BOLD:AEH1279 | N/A | 0.00 | *Rhodeus ocellatus* (FFU018-20) | 9.78 |
| 3. | *Acipenser baerii* | BOLD:AAA3850 | 0.00 | 0.00 | *Pseudoscaphirhynchus kaufmanni* (FFU029-20) | 5.07 |
| 4. | *Pseudoscaphirhynchus hermanni* | BOLD:ABY4771 | 0.00 | 0.00 | *Pseudoscaphirhynchus kaufmanni* (FFU029-20) | 0.00 |
| 5. | *Pseudoscaphirhynchus kaufmanni* | | 0.49 | 0.74 | *Pseudoscaphirhynchus hermanni* (FFU026-20) | 0.00 |
| 6. | *Channa argus* | BOLD:ABW0047 | 0.05 | 0.15 | *Silurus glanis* (FFU529-21) | 21.21 |
| 7. | *Sabanejewia aurata* | BOLD:ADL3084 | 0.26 | 1.04 | *Paracobitis longicauda* (FFU427-21) | 18.07 |
| 8. | *Cottus spinulosus* | BOLD:ABX6144 | 0.18 | 0.44 | *Rhinogobius* sp. (FFU210-210-21) | 17.76 |
| 9. | *Capoeta heratensis* | BOLD:ABV5006 | 0.26 | 0.74 | *Luciobarbus conocephalus* (FFU159-21) | 5.73 |
| 10. | *Carassius auratus* | BOLD:AAA7176 | 0.25 | 0.59 | *Carassius gibelio* (FFU116-20) | 0.00 |
| 11. | *Carassius gibelio* | | 0.32 | 0.89 | *Carassius auratus* (FFU096-20) | 0.00 |
| 12. | *Cyprinus carpio* | BOLD:AAA7175 | 0.07 | 0.29 | *Schizothorax fedtschenkoi* (FFU203-21) | 10.38 |
| 13. | *Luciobarbus brachycephalus* | BOLD:ADC7698 | 0.25 | 0.44 | *Luciobarbus conocephalus* (FFU159-21) | 2.40 |
| 14. | *Luciobarbus conocephalus* | BOLD:AEH2022 | 0.08 | 0.15 | *Luciobarbus brachycephalus* (FFU141-20) | 2.40 |
| 15. | *Schizothorax eurystomus* | BOLD:AAG2555 | 0.24 | 0.59 | *Schizothorax* sp. (FFU666-21) | 5.73 |
| 16. | *Schizothorax fedtschenkoi* | BOLD:AEH2828 | 0.09 | 0.15 | *Schizothorax* sp. (FFU666-21) | 2.71 |
| 17. | *Schizothorax* sp. | BOLD:AEH5960 | 0.35 | 0.59 | *Schizothorax fedtschenkoi* (FFU203-21) | 2.71 |
| 18. | *Esox lucius* | BOLD:AAA5988 | 0.00 | 0.00 | *Paracobitis longicauda* (FFU441-21) | 20.65 |
| 19. | *Neogobius melanostomus* | BOLD:AAC0218 | N/A | 0.00 | *Neogobius pallasi* (FFU207-21) | 11.78 |
| 20. | *Neogobius pallasi* | BOLD:ADL4983 | N/A | 0.00 | *Neogobius melanostomus* (FFU207-21) | 11.78 |
| 21. | *Rhinogobius* sp. | BOLD:ACB4145 | 0.00 | 0.00 | *Cottus spinulosus* (FFU066-20) | 17.76 |
| 22. | *Abbottina rivularis* | BOLD:ACM1972 | 0.16 | 0.44 | *Leuciscus squaliusculus* (FFU394-21) | 15.89 |
| 23. | *Gobio nigrescens* | BOLD:ADM0400 | 0.00 | 0.00 | *Gobio sibiricus* (FFU283-21) | 5.55 |
| 24. | *Gobio lepidolaemus* | BOLD:AAC5606 | 0.28 | 0.59 | *Gobio sibiricus* (FFU283-21) | 0.89 |
| 25. | *Gobio sibiricus* | | 0.00 | 0.00 | *Gobio lepidolaemus* (FFU271-21) | 0.89 |
| 26. | *Pseudorasbora parva* | BOLD:AAD0138 | 0.63 | 1.48 | *Gobio lepidolaemus* (FFU271-21) | 15.33 |
| 27. | *Abramis brama* | BOLD:AAC8592 | 0.00 | 0.00 | *Capoetobrama kuschakewitschi* (FFU365-21) | 6.37 |
| 28. | *Alburnoides holciki* | BOLD:ADK0904 | 0.36 | 1.19 | *Abramis brama*(FFU296-21) | 10.37 |
| 29. | *Alburnus chalcoides* | BOLD:AAB6908 | 0.13 | 0.29 | *Alburnus taeniatus* (FFU339-21) | 3.64 |
| 30. | *Alburnus oblongus* | BOLD:AAB6906 | 0.11 | 0.29 | *Alburnus taeniatus* (FFU339-21) | 2.25 |
| 31. | *Alburnus taeniatus* | | 0.00 | 0.00 | *Alburnus oblongus* (FFU361-21) | 2.25 |
| 32. | *Capoetobrama kuschakewitschi* | BOLD:AEH1599 | 0.00 | 0.00 | *Abramis brama*(FFU296-21) | 6.37 |
| 33. | *Leuciscus aspius* | BOLD:AAC8137 | N/A | 0.00 | *Leuciscus lehmanni* (FFU377-21) | 6.03 |
| 34. | *Leuciscus lehmanni* | BOLD:AAE3495 | 0.21 | 0.59 | *Leuciscus squaliusculus* (FFU391-21) | 0.00 |
| 35. | *Leuciscus squaliusculus* | | 0.37 | 0.74 | *Leuciscus lehmanni* (FFU380-21) | 0.00 |
| 36. | *Pelecus cultratus* | BOLD:AAF5575 | 0.07 | 0.15 | *Abramis brama*(FFU296-21) | 10.04 |
| 37. | *Rutilus lacustris* | BOLD:AAA5492 | 0.21 | 0.44 | *Capoetobrama kuschakewitschi* (FFU365-21) | 8.02 |
| 38. | *Dzihunia amudarjensis* | BOLD:AEH1522 | 0.00 | 0.00 | *Dzihunia* sp.1 (FFU603-21) | 7.16 |
| 39. | *Dzihunia* sp.1 | BOLD:AEH3205 | 0.16 | 0.44 | *Dzihunia* sp.2 (FFU643-21) | 5.05 |
| 40. | *Dzihunia* sp. 2 | BOLD:AEH3654 | 0.00 | 0.00 | *Dzihunia* sp.1 (FFU638-21) | 5.05 |
| 41. | *Dzihunia* sp. 3 | BOLD:AEH8655 | 0.04 | 0.15 | *Dzihunia* sp.1 (FFU603-21) | 8.69 |
| 42. | *Paracobitis longicauda* | BOLD:AEH0433 | 0.33 | 1.18 | *Dzihunia* sp.3 (FFU651-21) | 13.42 |
| 43. | *Triplopysa ferganaensis* | BOLD:AEH3382 | 0.00 | 0.00 | *Triplophysa* sp.2 (FFU610-21) | 5.86 |
| 44. | *Triplophysa strauchii* | BOLD:AAH2273 | 0.22 | 0.74 | *Triplophysa* sp.2 (FFU610-21) | 6.54 |
| 45. | *Triplohysa* sp.1 | BOLD:AEH5574 | 0.00 | 0.00 | *Triplophysa strauchii* (FFU453-21) | 8.32 |
| 46. | *Triplopysa* sp. 2 | BOLD:AEH6371 | 0.00 | 0.00 | *Triplophysa ferganaensis* (FFU614-21) | 5.86 |
| 47. | *Sander lucioperca* | BOLD:AAD1749 | 0.10 | 0.29 | *Cottus spinulosus* (FFU066-20) | 18.79 |
| 48. | *Gambusia holbrooki* | BOLD:AAC2757 | 0.15 | 0.29 | *Rhinogobius* sp. (FFU210-21) | 22.19 |
| 49. | *Oncorhynchus mykiss* | BOLD:AAA1627 | 0.15 | 0.15 | *Rhodeus* sp. (FFU022-20) | 20.99 |
| 50. | *Silurus glanis* | BOLD:ACL1933 | 0.24 | 0.59 | *Gobio lepidolaemus* (FFU269-21) | 18.32 |
| 51. | *Glyptosternon oschanini* | BOLD:ADW1603 | N/A | 0.00 | *Glyptosternon* sp. (FFU660-21) | 1.94 |
| 52. | *Glyptosternon* sp. | BOLD:AEH7649 | 0.00 | 0.00 | *Glyptosternon oschanini* (FFU-653-21) | 1.94 |
| 53. | *Ctenopharyngodon idella* | BOLD:ACL1923 | 0.14 | 0.29 | *Hypophthalmichthys molitrix* (FFU570-21) | 9.43 |

| | | | | | | |
|---|---|---|---|---|---|---|
| 54. | *Hemiculter leuciscuslus* | BOLD:ACB5189 | 0.44 | 1.04 | *Hypophthalmichthys molitrix* (FFU570-21) | 8.37 |
| 55. | *Hypophthalmichthys molitrix* | BOLD:AAF6633 | 0.27 | 0.74 | *Hypophthalmichthys nobilis* (FFU577-21) | 3.95 |
| 56. | *Hypophthalmichthys nobilis* | BOLD:ADK6840 | 0.00 | 0.00 | *Hypophthalmichthys molitrix* (FFU572-21) | 3.95 |
| 57. | *Mylopharyngodon piceus* | BOLD:AAD9723 | 0.44 | 0.44 | *Parabramis pekinensis* (FFU599-21) | 8.29 |
| 58. | *Opsariichthys bidens* | BOLD:ADG6045<br>BOLD:ADG6046 | 0.71 | 1.49 | *Ctenopharyngodon idella* (FFU541-21) | 12.35 |
| 59. | *Parabramis pekinensis* | BOLD:ACU1740 | 0.30 | 0.74 | *Mylopharyngodon piceus* (FFU582-21) | 8.29 |
